# Molecular grammars of intrinsically disordered regions that span the human proteome

**DOI:** 10.1101/2025.02.27.640591

**Authors:** Kiersten M. Ruff, Matthew R. King, Alexander W. Ying, Vicky Liu, Avnika Pant, Whitney E. Lieberman, Min Kyung Shinn, Xiaolei Su, Cigall Kadoch, Rohit V. Pappu

## Abstract

Intrinsically disordered regions (IDRs) of proteins are defined by functionally relevant molecular grammars. This refers to IDR-specific non-random amino acid compositions and non-random patterning of distinct pairs of amino acid types. Here, we introduce GIN (<u>G</u>rammars Inferred using <u>N</u>ARDINI+) as a resource, which we have used to extract the molecular grammars of all human IDRs and classified them into thirty distinct clusters. Unbiased analyses of IDRome-spanning grammars reveals that specialized IDR grammar features direct biological processes, cellular localization preferences, and molecular functions. IDRs with exceptional grammars, defined as sequences with high-scoring non-random features, are harbored in proteins and complexes that enable spatial and temporal sorting of biochemical activities. Protein complexes within the nucleus recruit specific factors through top-scoring IDRs. These IDRs are frequently disrupted via cancer-associated mutations and fusion oncoproteins. Overall, GIN enables the decoding of sequence-function relationships of IDRs and can be deployed in IDR-specific and IDRome-wide analyses.

## Introduction

Eukaryotic proteomes are composed of proteins with folded domains ^1^ and intrinsically disordered regions (IDRs) that contribute jointly to protein functions ^2–8^. In the human proteome, more than fifty percent of proteins contain at least one IDR, and are at least 30-residues in length ^4,9–11^. Proteins with IDRs are found in every cellular compartment, and IDRs are implicated in a range of different molecular functions and cellular processes ^3,12–14^.

The functions of folded domains can be inferred from evolutionary analysis of conserved or covarying sequence-structure relationships ^1,15,16^. However, this approach is not easily transferred to IDRs because their conformational heterogeneity imparts fewer constraints on sequences, thus often leading to poor sequence conservation ^4,17–23^. A different approach to uncovering sequence-function relationships comes from the finding that IDRs are defined by distinct *molecular grammars* ^20,24–36^. The amino acid alphabet is used in a biased manner and the non-random subset of the alphabet that is used by an IDR is arranged in non-random ways along the linear sequence. Accordingly, IDR-specific molecular grammars are defined jointly by the non-random amino acid composition and the non-random patterning of distinct pairs of amino acid types with respect to one another ^20,25,26,29,30,32,33,37–43^.

Molecular grammars of IDRs have been used to explain or quantify the driving forces for condensation ^24,27,32,35,41,42,44–47^, the protein compositions of condensates ^27,36,44,45,48–50^, and the molecular functions of IDRs ^21,33,37,51–53^. The interplay between short linear motifs ^54–56^ and their modulation by the sequence contexts afforded by IDRs can also be explained using molecular grammars ^21,25,28,57–63^. Building on these observations, recent efforts have focused on extracting and analyzing molecular grammars to infer IDR functions from their sequences ^64^. These efforts include unsupervised ^65^ or supervised machine-learning aided analysis of IDR sequences ^33,47,66^. Here, we pursue an approach based on the NARDINI+ ^21,27,44,48^ algorithm that can be readily deployed to analyze a single IDR or multiple IDRomes. NARDINI+ combines the NARDINI algorithm of Cohan et al.,^20^ and compositional analyses inspired by the work of Zarin et al.^18,65,67^. NARDINI+ has proven to be useful for extracting molecular grammars and sequence-function relationships of specific IDRs or families of IDRs ^21,44,48,68^.

Here, we use an unsupervised learning approach to enable the unbiased clustering and characterization of IDRs spanning the entire human proteome based on their molecular grammars. Within this IDRome-spanning basis set, we identify thirty distinct clusters, each with a unique fingerprint defined by standout features of the cluster-specific grammars. We have built and integrated this IDRome spanning grammars known as GIN (**G**rammars **I**nferred using **N**ARDINI+) into a user-friendly resource, supported by two Google Colab notebooks. The use of GIN enables IDRome-wide and IDR-specific analyses, predictions regarding IDRs of unknown localization and function, de novo design of new IDR sequences, and the testing of hypotheses regarding IDR roles in normal and disease states. We demonstrate the use of GIN by analyzing different clusters to show how distinct grammars contribute to distinct cellular localization preferences, temporally organized biological processes, molecular functions, and disruption of the above in the context of human cancers.

## Results

### Extraction of molecular grammars using NARDINI+

The NARDINI+ algorithm follows a defined workflow with an IDR sequence as the input. A set of 54 compositional features are analyzed. The usage of the alphabet, as instantiated in the specific IDR sequence, yields 54 distinct compositional z-scores. Next, the NARDINI algorithm ^20^ is used to compute 36 non-random binary patterns within the IDR sequence. Taken together, NARDINI+ generates a 90-component z-score vector (ZSV) as the output (**Figure 1A**).

**Figure 1:**
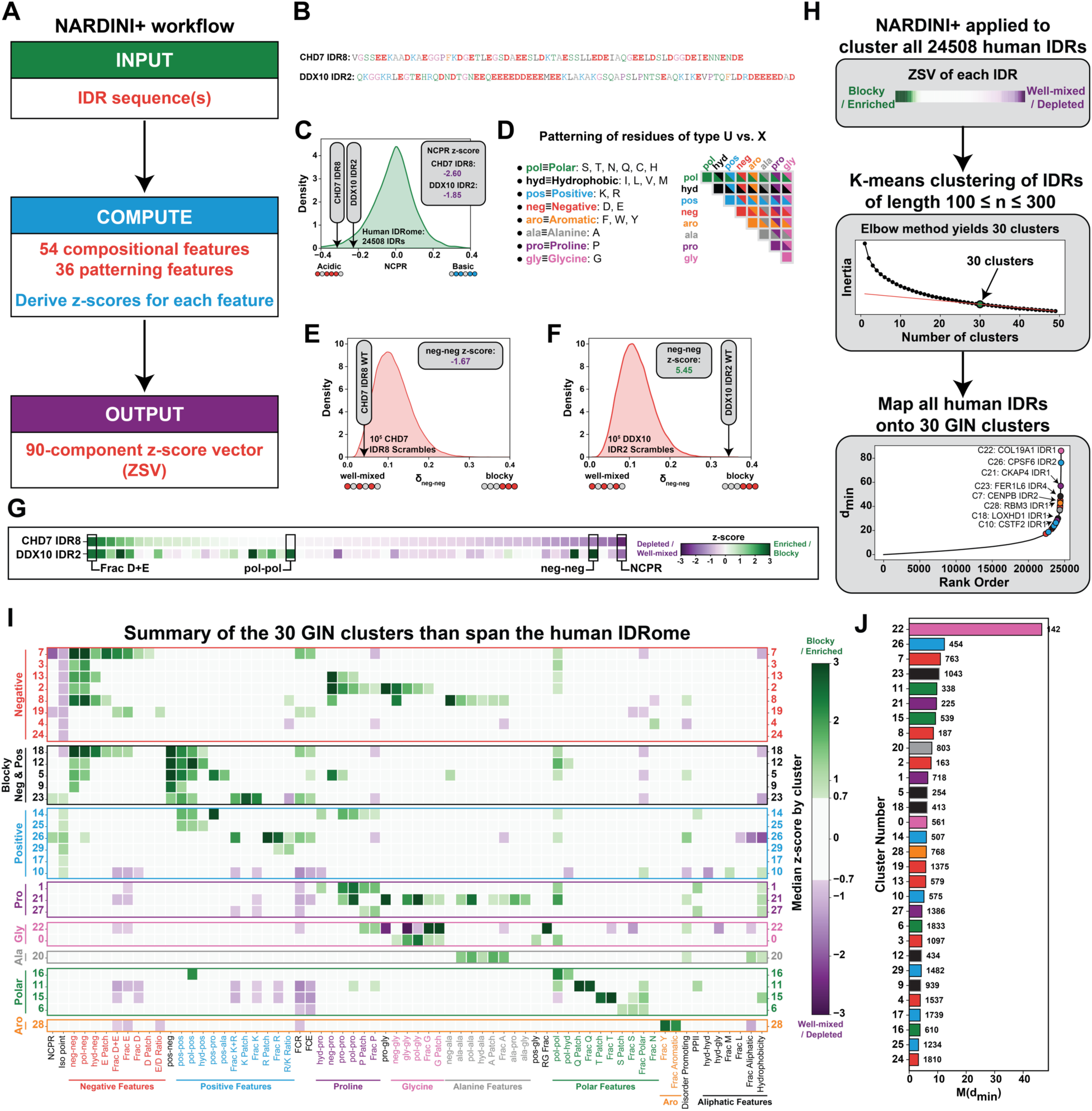
The NARDINI+ algorithm and generation of a basis set of grammars that span the human IDRome. See Figure S1. A. The NARDINI+ workflow. B. The sequence of the IDR or list of IDRs is first used to extract molecular grammars for the input IDR or the set of input IDRs. C. Illustrations of how sequence-specific compositional z-scores are calculated for each IDR. The distribution from the human IDRome is used as the prior distribution. The z-scores are -2.60 and -1.85, respectively for the CHD7 IDR8 and DDX10 IDR2. Negative values imply that the NCPR values observed here are more negative, by 2 or more standard deviations, when compared to the mean of the human IDRome. D. Grouping of amino acid types and the computation of a patterning matrix, where each element δ_UX_ quantifies the degree of mixing / segregation of residue types U and X. E. To derive z-scores for patterning features, the prior distribution is constructed from 10^5^ randomly scrambled sequences while keeping the composition fixed to be that of the IDR of interest. We find that the z-scores for the mixing / segregation of negatively charged residues with respect to one another (neg-neg) are such that these residues are more well-mixed for CHD7 IDR8 (left, z_neg-neg_ = -1.67). F. Same as E, except we find that the z-scores for the mixing / segregation of negatively charged residues with respect to one another (neg-neg) are such that these residues are more segregated for DDX10 IDR2 (z_neg-neg_ = +5.45) than the random prior. G. Examples of ZSVs computed for two different IDRs. Only features in which the z-scores were non-zero for at least one of the IDRs are shown. The compositional (fraction of D+E and NCPR) and patterning parameters (segregation / mixing of polar residues with respect to one another, pol-pol, and segregation / mixing of anionic residues with respect to one another, neg-neg) stand out as distinctly non-random features for the pair of IDRs. H. Schematic of the workflow used for the generation of GIN clusters that span the human IDRome. The collection of clusters constitutes a basis set. I. The median ZSV showing the standout features within each of the 30 GIN clusters. The data are shown for GIN clusters generated using IDRs of length 100 ≤ *n* ≤ 300 where *n* refers to the number of residues. For ease of visualization, we grouped the ZSVs into eight super-categories. These groups, which are shown with distinctive colors for the bounding boxes are as follows: (1) negatively charged (red), (2) blocky polyampholytes (black), (3) positively charged (blue), (4) proline-rich (purple), (5) glycine-rich (pink), (6) alanine-rich (gray), (7) polar-rich (green), and (8) aromatic-rich (orange) clusters. J. The minimum inter-cluster distance is denoted as d_min_ and the figure shows the median of d_min_ or M(d_min_) (see **Methods**) that calculates the goodness of mapping of each IDR to a cluster. Each bar, the length of which corresponds to the value of M(d_min_), has a number at the right that refers to the number of IDRs from the human IDRome that map onto the cluster.

The procedural details of NARDINI+, illustrated using two example sequences from the human IDRome (**Figure 1B**), are as follows: For an IDR sequence, the fraction of each of the 20 naturally occurring residues can be calculated for each of the sequences. This information is then used to derive 34 additional compositional sequence features that include ^34,46,48^: 8 values that quantify the fractions of specific types of residues *viz*., polar (Asn, Cys, Gln, Gly, His, Ser, Thr), aliphatic (Ala, Ile, Leu, Met, Val), aromatic (Phe, Trp, Tyr), the fraction of Lys plus Arg (K+R), fraction of Asp plus Glu (D+E), fraction of charged residues (FCR), fraction of residues that promote chain expansion (FCE), and the fraction of disorder promoting residues (Disorder); 20 values that quantify the presence of specific residue or RG “patches”; 2 values that quantify the ratios of Arg-to-Lys (R/K Ratio) and Glu-to-Asp (E/D Ratio) residues; and 4 values that quantify the net charge per residue (NCPR), the apparent isoelectric point (pI), the Kyte-Doolittle hydrophobicity ^69^, and the overall propensity for adopting polyproline II conformations. The choice of compositional parameters covers all known and established parameters that are used to classify IDRs and delineate them from folded domains ^7,8,12,13,70^ or describe the conformational or functional preferences of IDRs ^3,6,10,11,18,21,24–34,37,44,47,48,53,59,60,65,67,71–86^.

To quantify whether any of the 54 compositional sequence features are non-random for a given IDR, NARDINI+ compares each feature to the distribution of that feature for all IDRs in the human IDRome (**Figure 1C**). Specifically for each feature, a z-score is computed as the signed deviation of the feature from the mean of the distribution, referenced to the standard deviation of the distribution. If the z-score for a compositional feature lies in the interval -1 ≤ z ≤ +1, then the feature in question conforms to the random prior defined by the human IDRome. However, if |z| > 1, then the feature, as represented in the IDR of interest, is non-random and the extent of non-randomness is set by the magnitude of the z-score. In general, a negative z-score implies that the compositional feature is depleted in the IDR when compared to the reference IDRome, whereas a positive z-score implies that the feature is enriched when compared to the reference IDRome. For features such as the isoelectric point (pI), a negative z-score implies a more acidic pI, and a positive z-score implies a more basic pI than the mean of the human IDRome. Similar considerations apply to parameters such as net charge per residue.

In addition to compositions, the linear patterning of distinct pairs of residue types also contribute to sequence-function relationships ^25,26,29,31,72,87^. Cohan et al.,^20^ introduced the NARDINI algorithm to compute 36 distinct binary patterning features for a given IDR. These features are computed based on the grouping of residues into eight types *viz*., polar, hydrophobic, positively charged, negatively charged, aromatic, alanine, glycine, and proline (**Figure 1D**). This grouping was inspired by discoveries of the types of binary patterns that influence IDR-encoded conformational ensembles and functions ^7,25,26,29–32,41–43,49,50,88^. For each unique pair of residue types U and X, the NARDINI algorithm computes a parameter δ_UX_ that quantifies the extent to which U and X are mixed or segregated along the linear sequence ^29,25,26^ (**Figure 1E-F**). To compute the z-score for each patterning feature, 10^5^ randomly scrambled sequences are generated while keeping the composition fixed to be that of the IDR of interest. The scrambled sequences yield a full distribution of realizable δ_UX_ values for a fixed composition. The mean and standard deviation for the values of δ_UX_ are then used in conjunction with the actual value of δ_UX_ to compute the z-score for this feature. The z-score is set to zero if the fraction of U or X constitutes less than 10% of the IDR. If the z-score for a specific δ_UX_ lies in the interval -1 ≤ z ≤ +1, then the patterning of U and X with respect to one another is random. If |z| > 1, then the U and X residue types are patterned non-randomly in the linear sequence. The extent of non-randomness is set by the magnitude of the z-score. A negative z-score implies that the U and X residue types are well-mixed i.e., uniformly distributed within the linear sequence ^20^ (**Figure 1E**). Conversely, a positive z-score implies that the U and X residue types are segregated into distinct blocks within the linear sequence when compared to the ensemble of scrambled sequences ^20^ (**Figure 1F**).

Taken together, for a given IDR, the NARDINI+ algorithm yields 54 compositional z-scores and 36 z-scores for distinct binary patterns. These are organized into a 90-component z-score vector (ZSV) (**Figure 1G**).

### Generation of a basis set of grammars that spans the human IDRome

Next, we used the ZSVs, computed for every IDR in the human IDRome, and clustered them to ask if **G**rammars **I**nferred using **N**ARDINI+ (GIN) can be used to generate a basis set of grammars that spans the human IDRome (**Figure 1H**). We clustered ZSVs across the human IDRome. For this, we initially focused on IDRs of length 100 ≤ *n* ≤ 300 from the human IDRome. This choice was made to: (1) minimize artifacts due to finite lengths that can affect the analysis of compositional features; (2) focus on IDR lengths for which grammars have been shown to modulate localization, function, and phase separation in the context of the full protein ^18,44,48,49^; (3) maximize the ability to separate grammars into distinct clusters by minimizing signal-to-noise issues that crop up when analyzing short IDRs.

The ZSVs from NARDINI+ were extracted for 4,529 IDRs that meet the stipulated length range. Next, we used K-means clustering as an unsupervised learning approach ^89^ to cluster the 4,529 ZSVs that pass the stipulated length filter (**Figure 1H**). Thirty clusters were found to be sufficient to span the 4,529 IDRs, which we refer to as GIN clusters (**Figure 1I, S1A**). Similar GIN clusters were extracted when all IDRs are chosen, although the goodness of categorization of IDRs decreased (**Figure S1B-S1C**). To summarize the main inferences, the cluster-specific signatures of ZSVs are depicted in terms of the median ZSV for each cluster (**Figure 1I**).

Next, we mapped all remaining 19,979 IDRs from the human IDRome using the centroids of each of the 30 GIN clusters. Following this mapping, each GIN cluster is defined by the distinctiveness of the cluster (**Figure 1J**). Here, we calculated the distance between the ZSV of an IDR and the centroid of the cluster onto which the IDR was mapped. We also calculated the distance between ZSV of the IDR and centroids of all other clusters to quantify the next closest distance. We refer to the difference between these two values as the Minimum Inter-Cluster Distance (d_min_). Values of d_min_ that are larger than 5 imply that the IDR is well-mapped onto its given cluster, with larger values implying stronger mappings.

GIN clusters with large numbers of IDRs (>10^3^) often contain more weakly mapped IDRs. An exception to this is the cluster defined by non-random blocky patterning of positive residues with Lys patches (cluster 23). Other distinct clusters include cluster 7 defined by the presence of long D/E-tracts, cluster 26 defined by the presence of Arg patches, and cluster 11 defined by the presence of Q-tracts ^90^ (**Figure 1J, S1A**). Other clusters of significance include cluster 21, which is defined by blocks of Pro, Gly, Ala, and polar residues; cluster 20, which is defined by Ala patches; and cluster 28, which is defined by the high fraction of aromatic residues, specifically Tyr (**Figure 1J, S1A**). Finally, cluster 22 IDRs are defined by the uniform linear distribution of Pro and Gly with respect to one another. This feature strongly segregates cluster 22 IDRs from all other GIN clusters and is a defining grammar of elastomers and collagens ^91^.

### As a resource, GIN can be used to extract and analyze IDR grammars

We have created two Google Colab notebooks to facilitate the use of GIN as a resource (see Methods). The first notebook takes advantage of the fact that GIN was created by analyzing 24,508 IDRs from the human IDRome. Using this notebook, users can input a list of proteins using either gene names or UniProt accessions (**Figure S1D**). The lists can be comma separated or they can be one gene / accession per line file (**Figure S1E**). The analysis produces two outputs (**Figure S1D**): (1) a summary of GIN annotations (**Figure S1F**) and (2) NARDINI+ z-score vectors for each of the IDRs in the gene / accession list (**Figure S1G**). The user has several optional outputs, and these include sequence schematics of proteins from the gene / accession list (**Figure S1H**). Here, IDRs are colored and labeled by their GIN cluster. UniProt extracted domains are shown in yellow and labeled by the domain type ^92^. Users can also plot hierarchically clustered heatmaps of the ZSVs, which can be used for identifying exceptional grammars defined as features with absolute values of z-scores that are higher than a user-defined threshold. Information on exceptional grammars can be used to design IDRs that manipulate these grammars in order to help unmask or probe the consequences sequence-function relationships of IDRs.

The second notebook allows users to input their chosen IDR or a list of IDRs in FASTA format ^93^. This notebook is helpful if users prefer to specify bespoke sequence regions for the IDRs of interest. For the given list of IDRs, this notebook also extracts GIN clusters and ZSVs. However, since compositional and patterning ZSVs have to be calculated anew, the analyses are initially split. This is due to the fact that the extraction of patterning ZSVs requires that thousands of scrambles be generated per IDR, and this can take a few minutes to complete, depending on the length and number of IDRs. To facilitate reasonable run times, we have set the maximum number of IDRs to 20 and the maximum length of the IDR to 10^3^ for the calculation of the patterning ZSVs. For more or longer IDRs, we encourage users to perform the NARDINI analysis as part of localCIDER distribution ^34^. In this notebook, users have the option to download just the computational ZSVs, just the patterning ZSVs, or the full 90 feature ZSVs to either a .excel or .csv file.

Having summarized NARDINI+ and used it to generate GIN clusters, the union of which spans the human IDRome, we set out to analyze sequence-function relationships of IDRs through the lens of GIN clusters.

### GIN clusters with specific grammars are predictive of sub-nuclear localization

Biological processes are often spatially organized into distinct membrane-bound ^94^ or membraneless compartments ^95^. We asked if there was an enrichment of specific GIN clusters within specific organelles or specific membraneless compartments. For this analysis, we extracted annotations of subcellular locations of proteins with IDRs from the Human Protein Atlas (HPA) ^96^. We focused on IDRs of length ≥ 70 and non-linker IDRs of length ≥ 50 that have with high d_min_ values, which means that their mapping onto GIN clusters is unique and strong. This allowed us to analyze IDRs that unambiguously confer functional preferences within the context of full-length proteins because the number of residues is equivalent to that of a canonical folded domain ^1,18,44,48,49^.

We find that IDRs from distinct GIN clusters are enriched in different subcellular locations (**Figure 2A-2B**). Previous studies have also uncovered distinct IDR sequence signatures for proteins that localize to nucleoli and nuclear speckles ^45,48^. Accordingly, we tested whether these grammars were recovered in an unbiased analysis based on GIN clusters. We find that nucleoli are enriched in cluster 23 K-block IDRs and furthermore, all experimentally tested nucleolar localization peptides map to this cluster ^97^ (**Figure 2B**). Similarly, nuclear speckles are enriched in cluster 26 R-patch rich IDRs and all speckle localizing IDRs ^45^ map to this cluster (**Figure 2B**). These results suggest that IDRs, annotated based on the GIN cluster to which they are mapped, can be used to predict localization of proteins ^24,36,45,48,49^. This was recently referred to as protein codes ^66^.

**Figure 2:**
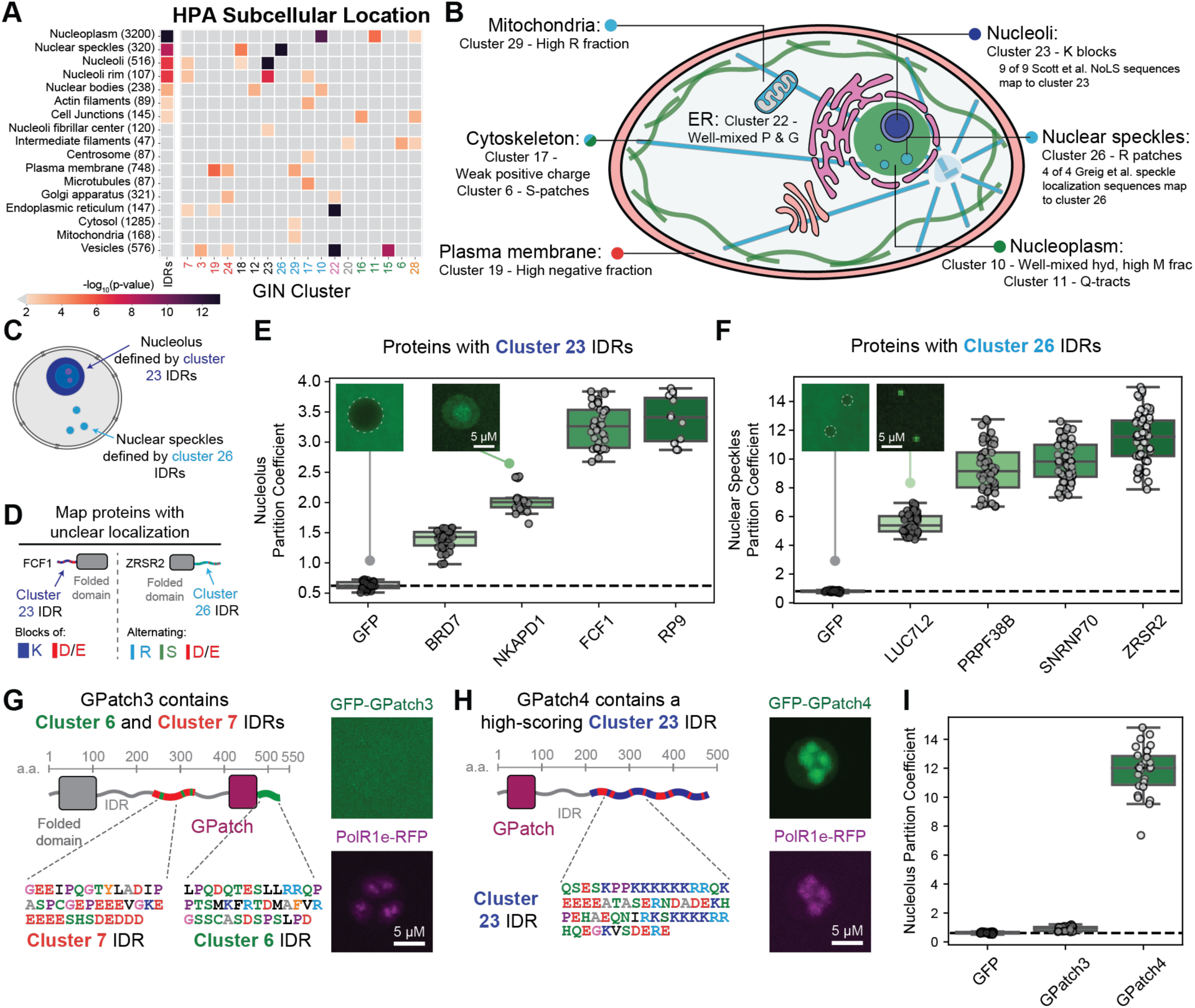
GIN clusters with specific grammars are predictive of sub-nuclear localization. See Figure S2. A. Enrichment of proteins containing IDRs (column 1) and IDRs in GIN clusters (remaining columns) for HPA subcellular location. Numbers in parentheses indicate the number of IDRs associated with the given HPA subcellular location annotation. Labels along the abscissa show the GIN cluster IDs. The heat map shows the -log_10_(p-value). Larger values imply higher statistical significance. The p-values were derived from Fisher’s exact test. B. Schematic of GIN clusters enriched in subcellular locations. Location specific GIN cluster enrichment is consistent with previous studies ^45,97^. C. Schematic showing that nucleolar proteins harbor IDRs defined by GIN cluster 23 and nuclear speckle proteins harbor IDRs defined by GIN cluster 26. D. Example proteins with either cluster 23 (FCF1) or cluster 26 (ZRSR2) IDRs that have ambiguous or unknown sub-cellular localization according to HPA. E. Data for four proteins, with cluster 23 IDRs and ambiguous HPA localization show partitioning into nucleoli of living *Xenopus* oocyte nuclei. F. Data for four proteins, with ambiguous HPA localization annotations featuring IDRs from cluster 26, show partitioning into nuclear speckles of living *Xenopus* oocyte nuclei. G. Features of the GPatch3 protein include its RNA-binding folded domain and two IDRs corresponding to GIN clusters 6 and 7. A representative confocal image of a nucleolus from a living *Xenopus* oocyte nucleolus expressing GFP-GPatch3 and Polymerase I subunit E tagged with RFP H. Features of the GPatch4 protein include its RNA-binding folded domain and a long IDR corresponding to GIN cluster 23. A representative confocal image of a nucleolus from a living *Xenopus* oocyte nucleolus expressing GFP-GPatch4 and Polymerase I subunit E tagged with RFP. I. Partition coefficients for GPatch3 and GPatch4 partitioning into nucleoli. For all partition coefficient plots: grey circles correspond to individual nucleoli or nuclear speckles and standard box and whisker statistics are used to show all the data that were collected (see methods for details).

To further test whether nucleolar and nuclear speckle specific GIN clusters are reasonable predictors of partitioning into these bodies, we studied the spatial localization preferences of full-length proteins containing cluster 23 and 26 IDRs using germinal vesicles (GVs) from live *Xenopus laevis* oocytes (**Figure 2C-D, S2A**). We focused on monomeric proteins that contain IDRs that map onto clusters 23 and 26 (**Figure S2B-S2C**). For these proteins, the HPA annotation regarding localization was ambiguous with the annotations being nucleoplasmic, cytosolic, or unknown. Note that HPA is the most comprehensive annotation of intracellular localization preferences. While GFP is excluded from nucleoli, with a partition coefficient that is less than one, all four human proteins containing cluster 23 IDRs partitioned into nucleoli in *Xenopus laevis* GVs (**Figure 2E, S2D**). Likewise, all four human proteins containing cluster 26 IDRs partitioned into nuclear speckles in *Xenopus laevis* GVs (**Figure 2F, S2E**). For nucleoli, we used RNA polymerase I as a fiduciary marker. For nuclear speckles, we took advantage of the fact that speckles and Cajal bodies in *Xenopus laevis* GVs are readily identifiable using differential interference contrast (DIC) microscopy as shown by Gall and coworkers ^98^. It is worth noting that the partition coefficients we measure here are larger than what has been recently reported in attempts to use IDR grammars, derived from machine learning, to map sub-nuclear protein localization ^66^. Our results suggest that, in *Xenopus laevis* GVs, the defining molecular grammars of cluster 23 and 26 IDRs function as strong signals that determine localization to nucleoli versus nuclear speckles, respectively.

IDRs alone cannot unambiguously determine sub-cellular, sub-nuclear, or sub-compartmental localization. Instead, localization preferences of proteins are jointly determined by IDR grammars and the folded domains to which the IDRs are tethered ^48^. Accordingly, we investigated the synergy between folded domains and IDRs using two naturally occurring GPatch proteins, which have been proposed to regulate the functions of DEAH/RHA RNA helicases ^99^. The folded GPatch domains are partially folded RNA binding domains with six well-conserved glycine residues ^100^. The proteins GPatch3 and GPatch4 each contain a single GPatch domain. GPatch3 features a disordered linker that maps onto cluster 7 (D/E tracts), and a C-terminal cluster 6 IDR (**Figure 2G**). In contrast, GPatch4 contains a C-terminal cluster 23 IDR (**Figure 2H**). For proteins with similar folded domains, the expectation would be that the IDR grammars are the primary determinants of localization preferences. Direct measurements within GVs from live *Xenopus laevis* oocytes are consistent with this hypothesis (**Figures 2I**). This is in accord with observations that the presence of a cluster 23 IDR is a driver of localization to nucleoli (**Figure 2E**) ^48^.

Overall, our data and recent work ^66^ suggest that the presence of IDRs with specific grammars is necessary and can be sufficient for determining the localization preferences of proteins ^24,36,45,48,49^. Previous studies showed that the localization preferences of proteins with IDRs derived from a specific GIN cluster will be governed by the folded domains to which they are tethered ^48^. Here, we show that the complement of this is true as well because the localization preferences of proteins featuring identical folded domains are governed by the IDR grammars. Taken together with the corpus of published work ^24,36,45,48,49,66^, our findings suggest that there might be designable or evolvable zip codes whereby folded domains, clustered using CATH ^101^ or SCOP ^102^, can be combined with GIN clusters to program, analyze, or evolve sub-cellular localization patterns of proteins.

### Distinct functions are enriched with IDRs from distinct GIN clusters

We next asked if specific GIN clusters are associated with specific molecular functions and biological processes that go beyond determining localization preferences. We first assessed the enrichment of each GIN cluster in all high-level Gene Ontology (GO) ^103,104^ Slim molecular function terms with at least 25 IDRs (**Figure 3A**). Proteins involved in DNA and RNA binding are enriched in IDRs and they house IDRs with different grammars. The GIN clusters with IDRs annotated as being enriched in RNA binding show a bias towards a high fraction of charged residues (FCR). Accordingly, we calculated the FCR distribution for all IDRs and compared these to IDRs housed in proteins associated with RNA binding and IDRs housed in proteins associated with DNA binding (**Figure 3B**). IDRs housed in proteins associated with RNA binding have higher FCR values than IDRs housed in proteins associated with DNA binding (Mann-Whitney-Wilcoxon p-value=3.5×10^-26^) and all IDRs (Mann-Whitney-Wilcoxon p-value= 2.4×10^-33^). In contrast, we did not observe a significant difference in FCR values between IDRs housed in proteins associated with DNA binding when compared to all IDRs (Mann-Whitney-Wilcoxon p-value=0.49). Similar results were obtained in examining high-level GO Slim biological process terms that have at least 25 IDRs associated with these processes (**Figure 3C**). Biological processes that are enriched in IDRs and are associated with distinct IDR grammars include the regulation of DNA-templated transcription, chromatin organization, and mRNA metabolism. Cluster 11 IDRs, which are defined by Q-tracts, are most enriched in the regulation of DNA-templated transcription ^90^; cluster 18 IDRs, defined by the co-occurrence of large negative and positive blocks, are most enriched in chromatin organization; and the R-patch-rich IDRs of cluster 26 are most enriched in mRNA metabolism. These results are consistent with the observed enrichment of Q-rich repeats in transcriptional regulators ^90^, the enrichment of D, E, and K repeats in histone- and chromatin-binding IDRs ^65^, and splicing proteins housing R-rich IDRs ^45,65^. Our analysis also identifies additional GIN clusters in these IDR-enriched biological processes, implying that multiple types of molecular grammars may influence specific sets of biological processes. For example, regulation of DNA-templated transcription is highly enriched in the well-mixed hydrophobic cluster 10 as well as the blocky positive, negative, and polar cluster 12. These data suggest the utility of studies aimed to define sequence-function relationships of IDR grammars rather than larger deletions or replacements with IDRs that are not semantically interoperable ^36^.

**Figure 3:**
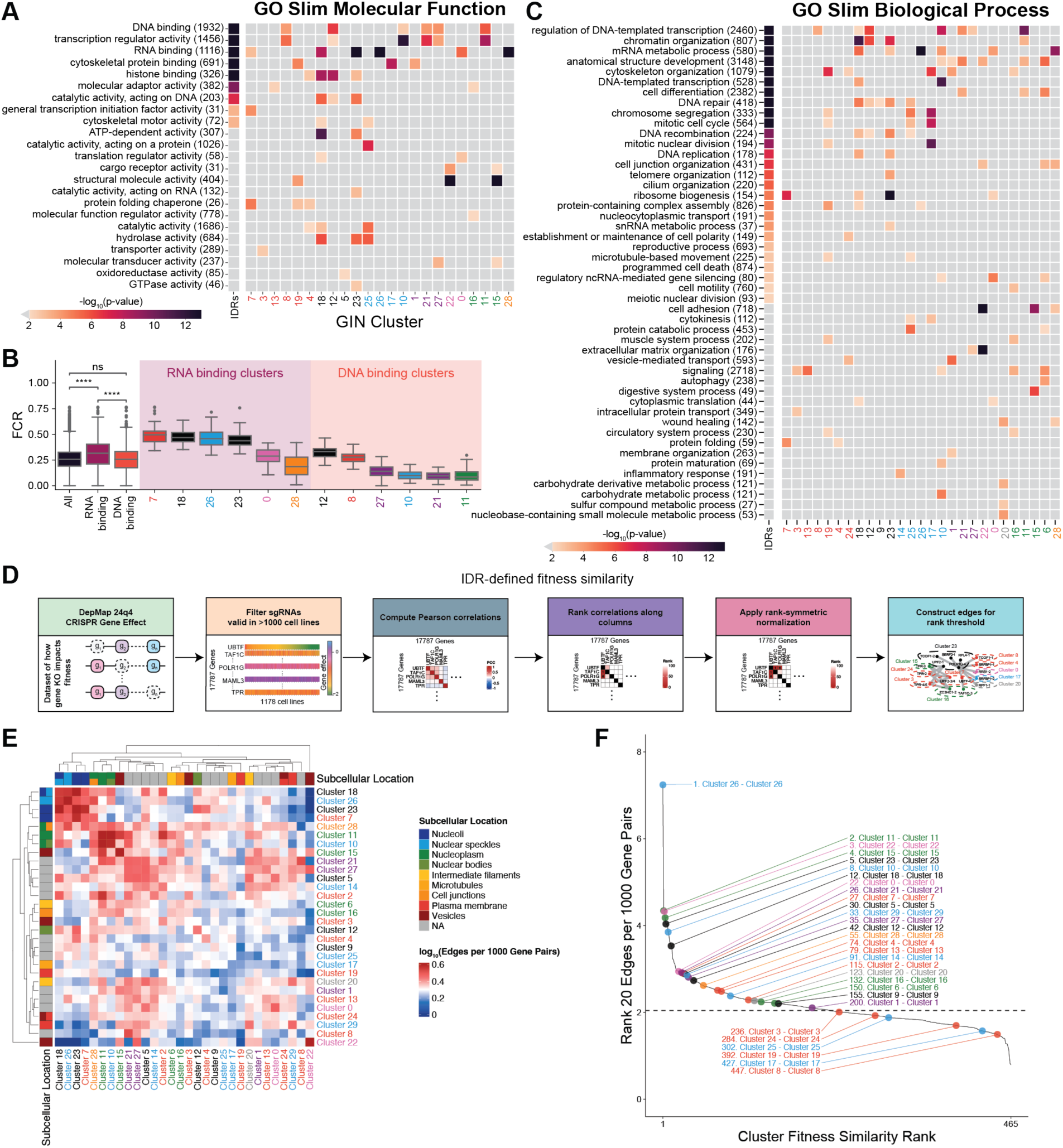
Distinct functions are enriched with IDRs from distinct GIN clusters. A. Enrichment of proteins containing IDRs (column 1) and IDRs in GIN clusters (remaining columns) for GO Slim molecular functions. Numbers in parentheses indicate the number of IDRs associated with the given molecular function. Labels along the abscissa show the GIN cluster IDs. The heat map shows the -log_10_(p-value). The larger this number, the higher is the statistical significance of the annotation. The p-values were derived from Fisher’s exact test. B. FCR distribution in IDRs of RNA and DNA binding proteins by their GIN cluster. Here, **** signifies p-values that are ≤ 10^-4^ computed via a Mann-Whitney-Wilcoxon t-test. C. Same as A, except for GO Slim biological processes. D. Schematic for analysis performed on the DepMap dataset to quantify functional relationships between pairs of genes as functional relationships between genes corresponding to proteins within pairs of GIN clusters. E. Heatmap of scores quantifying functional relationships between GIN clusters that were derived from the DepMap 24Q4 dataset. A high score is defined by a large number of edges and the colors bar shows annotation of the scores as log_10_ (edges per 10^3^ gene pairs). Subcellular locations enriched in specific GIN clusters are indicated on the plot. F. Plot of the ranked functional fitness correlations of gene pairs within (intra-cluster) or between (inter-cluster) GIN clusters. Points are shown for all intra-cluster pairs. Dashed line represents the overall edges per 1000 gene pairs, indicating that most intra-cluster networks are more highly connected than randomly selected average.

Next, we next sought to define whether proteins with IDR grammars that fall within distinct GIN clusters exhibit functional similarities (or differences). Functional similarity between a given pair or set of genes, and by extension, proteins, can be derived by the analysis of data from genome-scale CRISPR gene knockout screens performed across a sufficient genetic diversity of cellular contexts. For example, DepMap (Broad Institute) ^105^ has performed genome-scale CRISPR knockout screening in over 1000 cancer cell lines. Here, two genes (proteins) may be functionally linked if their fitness effects upon knockout across the cell lines are correlated. Using this DepMap dataset ^105,106^ and the methods of Pan et al., ^107^ we asked whether proteins housing IDRs with specific GIN clusters show functional relationships with proteins housing IDRs of the same and different GIN clusters (**Figure 3D**).

Applying this approach across all functional genetic relationships corresponding to protein IDRs within the defined GIN clusters, we identified inter- and intra-cluster correlations. Notably, high-scoring cluster fitness correlations included those defined by subcellular localization. These include GIN clusters 7, 23, and 18 that are enriched in nucleoli, GIN clusters 26 and 18 that are enriched in nuclear speckles, and GIN clusters 10 and 11 that are enriched in the nucleoplasm. However, high-scoring cluster fitness correlations also included those with distinct localization or with GIN clusters not enriched in a particular subcellular localization (cluster 2 and cluster 11; cluster 8 and cluster 21) (**Figure 3E**). Conversely, cluster 22 had zero detectable functional correlations with genes encoding protein IDRs in clusters localizing to the nucleus, such as 18, 26, and 23 (**Figure 3E**). Strikingly, the top ranked intra-cluster correlations are within clusters 26, 11, 22, 15, 23, and 10, respectively. These clusters are also highly enriched for specific subcellular locations (**Figure 2A**), thus suggesting that a major signal for the fitness data might be subcellular localization preferences driven, in part, by IDRs.

Examining ranks of gene pairs within GIN clusters and between GIN clusters reveals that 6 of the top 10 top-ranked gene (protein) fitness correlations were within GIN clusters (**Figure 3F**). These include four nuclear enriched clusters, specifically clusters 26, 11, 23, and 10. Together, these data reveal that functional relationships of genes, and their protein products correlate with selected IDR grammars. Fitness similarity analyses ^107,108^ can resolve gene perturbation effects based on whole gene knockouts or knockdowns; here, we used these data to uncover correlations within the GIN framework of IDR segments within protein products of perturbed genes.

The precise nature of IDR-mediated interactions that contribute to the fitness effects of pairs of GIN clusters obviates the need for new experimental investigations. The correlations we observe could involve direct interactions among IDRs with complementary grammars or they might also be the result of colocalization of genes encoding IDRs enriched in specific subcellular locations (**Figure 3E**). Investigations of functional relationships, through the lens of IDR-mediated interactions will be needed to obtain a clear understanding of causal relationships that underlie the correlations between GIN clusters. In the remainder of this work, we use collated information about specific biological processes and protein complexes in order to illustrate how GIN can be used to generate testable functional hypotheses and biological insights.

### GIN clusters enriched in sub-nuclear regions are associated with discrete processes

Nucleoli (p-value=2.9×10^-7^), nuclear speckles (p-value=8.0×10^-9^), and the nucleoplasm (p-value=3.4×10^-61^) are the subcellular locations most enriched in IDRs (**Figure 2A**). Furthermore, all three locations have specific functions and distinct enrichments of GIN clusters (**Figure 2A, 4A-4B**). This suggests that IDR grammars play specific functional role within sub-nuclear regions. Here, we highlight the specific functions and GIN clusters of nucleoli, nuclear speckles, and nucleoplasm. We then assess the prevalence of enriched GIN clusters in the discrete processes that are localized to three sub-nuclear regions.

We find that nucleolar IDRs are enriched in GIN clusters 23, 7, and 18 (**Figure 2A, 4A-4B**). Cluster 23 is defined by blocks of Ks, cluster 7 is defined by D/E-tracts, and cluster 18 is defined by large, negative blocks with positive blocks (**Figure 1I, 4B**). IDRs from each of these clusters are enriched in proteins that are involved in ribosome biogenesis – the main function of the nucleolus (**Figure 3C**). The nucleolus, however, is multifunctional ^109^ and proteins with IDRs from clusters 7, 18, and 23 are also enriched in other known functions of the nucleolus ^68^ (**Figure 3C**). These functions include DNA repair (clusters 18 and 23), DNA replication (clusters 18 and 23), and stress response, which include functions as protein folding chaperones (cluster 7) ^110–112^.

Ribosome biogenesis is a complex process whereby hundreds of proteins come together to organize 42 discrete and synergistic processes ^113^. Of the 42 discrete processes, 35 (83%) involve a protein with at least one IDR from cluster 7, 18, or 23 (**Figure 4C**). This is greater than an average of 21% for any other triplet of GIN clusters that excludes clusters 7, 18, and 23 (**Figure 4D**). Cluster 23 IDRs are most commonly found in proteins that are associated with discrete nucleolar processes (71%). These processes include the box C/D and H/ACA ribonucleoprotein complexes, nucleolar exosome, and the UTP-A and UTP-B complexes. We find a striking sorting of GIN cluster preferences based on the temporal ordering of processes involved in ribosomal biogenesis. IDRs in cluster 23 are prevalent in proteins associated with early-stage processes. In contrast to cluster 23, we find that that cluster 18 IDRs are preferred in proteins that are involved in later-stage ribosome biogenesis. Overall, the presence of cluster 7, 18, or 23 IDRs in ribosome biogenesis complexes / sub-processes suggests that the IDR grammars corresponding to these GIN clusters are important in nearly every step of ribosome biogenesis, with clear preferences based on the time points for ribosomal assembly.

**Figure 4:**
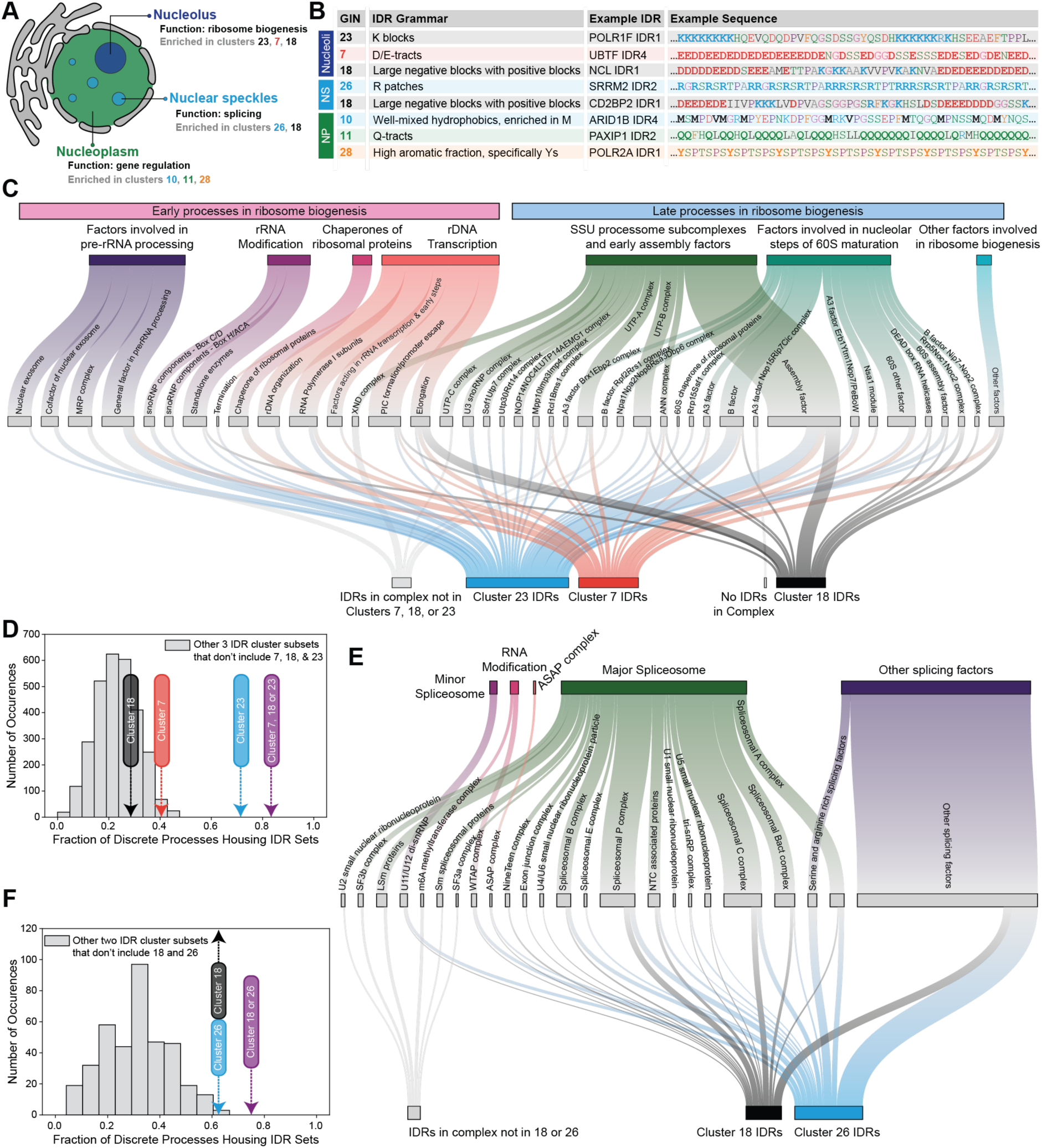
GIN clusters enriched in sub-nuclear regions are associated with discrete processes. See Figure S3. A. Schematic of the three sub-nuclear regions analyzed in this work. The schematic shows an annotation of each region by their main functions. It also lists the GIN clusters that are enriched within each region. B. Examples of IDRs, annotated by their GIN clusters and sequences that are enriched in specific sub-nuclear regions. Also shown are examples of specific proteins that harbor these IDRs. Residues that highlight the non-random grammars are shown in bold face. C. Sankey plot ^114^ showing the relationship between discrete ribosome biogenesis processes and whether the proteins associated with these processes house nucleolar-specific GIN clusters (clusters 23, 7, and 18). Edge widths between the general biological processes and the discrete processes denote the number of proteins in the given discrete process. Edge widths between the discrete processes and the GIN cluster IDR type denote the number of IDRs in the discrete process that are categorized by the given GIN cluster. IDRs that are not part of GIN clusters 7, 18, 23 are only shown if the discrete process does not involve proteins with IDRs classified into GIN clusters 7, 18, or 23. General ribosomal biogenesis processes with their discrete processes were collated from Dörner et al.^113^ (**Table S4**). Visualization was enabled by the use of D3Blocks (github.com/d3blocks). D. Comparison of the fraction of discrete processes ^113^ housing different sets of GIN clusters (**Table S4**). Here, 83% of discrete complexes have at least one IDR from cluster 7, 18, or 23 (purple arrow), compared to an average of 21% for any other triplet of clusters that do not include clusters 7, 18, or 23 (grey histogram). E. Sankey plot ^114^ showing the relationship between discrete splicing processes and whether the proteins in those processes house nuclear speckle specific GIN clusters (clusters 26 and 18). Edge widths between the general biological processes and the discrete processes denote the number of proteins in the given discrete process. IDRs that are not part of GIN clusters 18 or 26 are only shown if the discrete process does not involve proteins with IDRs classified into GIN clusters 18 or 26. General splicing processes with their corresponding discrete processes were collated from various sources in **Table S4** ^92,115,116^. Visualization was enabled by the use of D3Blocks (github.com/d3blocks).F. Comparison of the fraction of discrete splicing complexes housing different sets of GIN clusters. Here, 75% of discrete complexes have at least one IDR from cluster 18 or 26 (purple arrow), compared to an average of 30% for any other doublet of clusters that does not include either cluster 18 or 26 (grey histogram).

Next, we analyzed IDRs in nuclear speckle proteins. Nuclear speckles are membraneless compartments that function as storage and modification centers for pre-mRNA splicing factors ^117^. Speckles are formed near sites of active transcription and this helps couple transcription with splicing ^118^. Additionally, IDRs and their grammars have been shown to be important for nuclear speckle formation and localization ^45,119,120^.

IDRs localized to nuclear speckles are enriched in clusters 26 and 18 (**Figure 2A, 4A-4B**). Consistent with known functions of nuclear speckles, proteins with IDRs from clusters 18 and 26 are enriched in RNA binding, mRNA metabolism, and the spliceosome pathway ^121^ (**Figure 3A, 3C, S3A**). IDRs of cluster 26 are uniquely enriched in nuclear speckles and they are defined by the presence of Arg patches (R Patch) and a high fraction of Arg residues (Frac R) (**Figure 1I, 4B**). Importantly, the distribution of Arg residues in these IDRs cannot be described as being either highly blocky or well-mixed.

To determine the relevance of cluster 18 and 26 IDRs in general splicing functions, we collated proteins involved in splicing ^92,115–117,122^. Eighteen of the 24 discrete splicing categories have at least one cluster 18 or 26 IDR (75%). This contrasts with an average of 30% for any other doublet of clusters that does not include clusters 18 or 26 (**Figure 4E-F**). Early spliceosome assembly members including the serine- and arginine-rich splicing factors, the U1 small nuclear ribonucleoprotein, and other proteins that comprise the E complex lack cluster 18 IDRs and prefer cluster 26 IDRs ^123^.

Unlike speckles or nucleoli, the nucleoplasm is the key location for the regulation of gene expression, including RNA polymerase II regulated transcription ^44,64,124–126^. The nucleoplasm houses many components of transcriptional regulation machineries including the Mediator complex, general transcription factors, chromatin remodeling complexes, histone modifying complexes, transcription factors, and transcriptional co-factors ^125,127–133^. Components of the transcriptional regulation machinery are known to interact with one another to form transcriptional condensates at distinct genomic loci ^49,64,134^. Sequence grammars of IDRs can modulate the condensation of and recruitment to transcriptional condensates ^36,49,50,66,124,134–139^. Furthermore, IDR grammars can also modulate promoter binding preferences by changing the grammars of well-mixed hydrophobic residues ^64,140,141^.

Nucleoplasmic proteins are enriched in IDRs from clusters 10, 11, and 28 (**Figure 2A, 4A-4B**). Proteins that house cluster 10 and cluster 11 IDRs are enriched in DNA binding proteins and proteins that regulate DNA-templated transcription processes (**Figure 3A, 3C**). In contrast, proteins that house cluster 28 IDRs ^50^ are enriched in processes that involve RNA binding (**Figure 3A, 3C**). Cluster 10 IDRs are defined by well-mixed hydrophobic residues with a preference for Met residues, cluster 11 IDRs are defined by Q-tracts, and cluster 28 IDRs are defined by high fractions of aromatic residues with a preference for Tyr ^25,28,50^ (**Figure 1I, 4B**).

Next, we examined the relevance of the nucleoplasm-specific GIN clusters in discrete complexes that regulate gene expression (**Table S4**) ^92,115,125,127–129,131–133,142–144^. Most complexes that regulate gene expression house at least one IDR from clusters 10, 11, or 28 (**Figure S3B**). Specifically, 89% of discrete complexes have at least one IDR from clusters 10, 11, or 28, when compared to an average of 70% for any other triplet set of clusters that exclude these clusters (**Figure S3C**). Of the multiprotein complexes found in the nucleoplasm, the canonical BAF complex is unique in that it houses IDRs from all three GIN clusters (10, 11, and 28) that are enriched in the nucleoplasm. The presence of these IDRs has been shown to be important for diverse partner recruitment and targeting of the BAF complex to specific genomic locations^135,145^.

Overall, we find that proteins from nucleoli, nuclear speckles, and the nucleoplasm have IDRs from distinct GIN clusters *viz*., clusters 7, 18, and 23 (nucleoli), clusters 18 and 26 (nuclear speckles), and clusters 10, 11, and 28 (nucleoplasm). The involvement of IDRs from specific GIN clusters in specific molecular functions within distinct sub-nuclear regions suggests routes for targeting these IDRs via mutagenesis or design to probe sequence-function relationships enabled by these IDRs.

### Proteins involved in early processes and / or core complexes feature IDRs with exceptional grammars

Given that GIN clusters are defined by a few strong features we sorted IDRs in the clusters by their prominent features (**Figures S4-S5**). For GIN clusters defined by compositional features, we used the valence (number) rather than fraction of the given residue or patch because valence is a determinant of multivalency of ligand binding and the organization of protein interaction networks ^24,25,146–148^. When we sorted nucleolar IDRs, the top scoring cluster 7 and cluster 18 IDRs were found to be within the three key scaffolds UBTF, NPM1, and NCL^48,113,128,149–153^ (**Figure S4A-S4B**). Additionally, two RNA polymerase I specific subunits, POLR1F and POLR1G, contain the highest scoring cluster 23 IDRs (**Figure S4C**). Likewise, when we sorted nuclear speckle IDRs, the highest scoring cluster 26 IDR was found to be in SRRM2 (**Figure S4E**), which is a well-known splicing factor that is a component of the spliceosome. SRRM2 and SON, a splicing factor protein that houses another top scoring cluster 26 IDR, are essential for nuclear speckle formation ^119,154^. Lastly, for nucleoplasmic IDRs, the highest scoring cluster 28 IDR is the C-terminal domain of RNA polymerase II – an important player in all steps of the transcriptional process ^126,128^ (**Figure S5C**).

To test for evidence of functionality associated with exceptional features the full human proteome, we sorted all 24,508 IDRs by a given GIN cluster feature and examined if any protein in specific functional complexes were in the 99^th^ percentile. We find that proteins involved in the earliest ribosome biogenesis processes, rDNA transcription and rRNA modification, house exceptional K-block grammars, the defining feature of cluster 23 IDRs, relative to the entire human IDRome (**Figure 5A**). The transcription of rDNA has three main stages: initiation, elongation, and termination ^113^. Each of these stages involves proteins with IDRs that are among sequences with the top 80 pos-pos z-scores when sorted by the full human IDRome (**Figure 5B**). These include POLR1F and POLR1G of the pre-initiation complex, CTR9 and RTF1 of the Paf1C elongation complex, and TTF1 of the termination complex. These results suggest that exceptional K-block grammars are important for nucleolar functions that go beyond driving condensation and include upstream processes of ribosome biogenesis.

**Figure 5:**
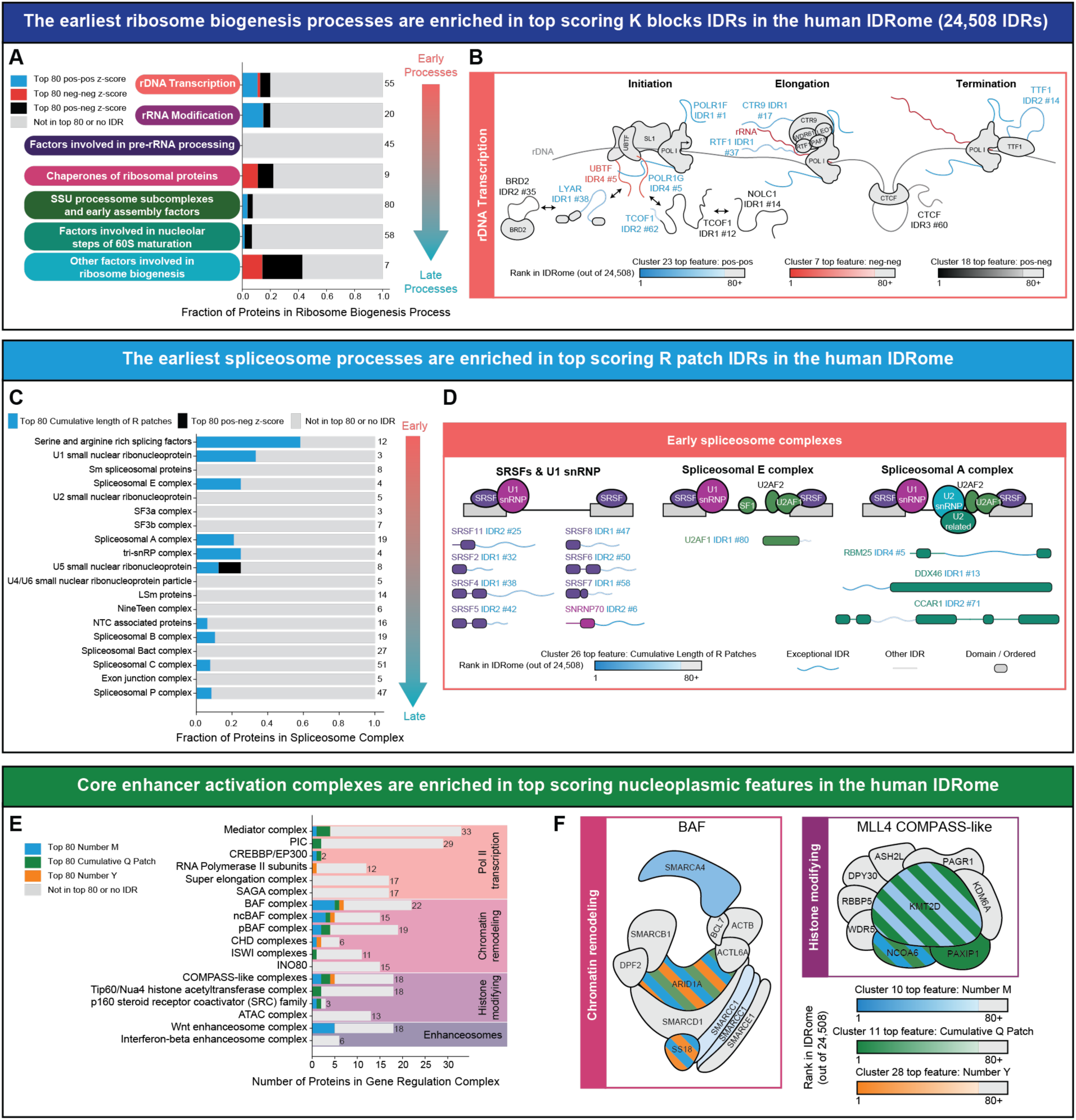
Proteins involved earliest processes and / or core complexes feature IDRs with exceptional grammars. See Figures S4-5. A. For each of the processes involved in ribosome biogenesis, the fraction of proteins housing IDRs with a top 80 pos-pos z-score (a cluster 23 feature), a top 80 neg-neg z-score (a cluster 7 feature), and a top 80 pos-neg z-score (a cluster 18 feature), extracted by sorting the full human IDRome (24,508 IDRs), are shown. Proteins involved in ribosome biogenesis are sorted by processes in early versus late stages of assembly. B. Schematic of all IDRs involved in rDNA transcription that are among the top 80 IDRs in terms of z-scores for pos-pos, neg-neg, or pos-neg patterns when the full human IDRome is sorted. Domains / complexes are shown as filled grey shapes. The color bars provide annotations for the coloring scheme used to depict the exceptional IDRs. C. For each spliceosome complex, the fraction of proteins housing IDRs with a top 80 z-score of cumulative length of R patches (a cluster 26 feature) and top 80 pos-neg z-score (a cluster 18 feature) when the full human IDRome is sorted (24,508 IDRs) are shown. Discrete spliceosome complexes are sorted in terms of their involvement in early versus late stages of splicing. D. Schematic of all IDRs involved in spliceosome processes through the spliceosomal A complex that are in the top 80 for cumulative length of R patches when the full human IDRome is sorted. The main complexes are shown as a schematic across the top and the individual proteins housing exceptional IDRs are shown as schematics below. Here, the domains in the proteins are colored to match the spliceosome complex they are a part of, and the color bar provides annotations for the scheme used to depict exceptional IDRs. E. For each gene regulation complex, the number of proteins housing IDRs with a top 80 number of M (cluster 10 feature), top 80 cumulative length of Q patches (cluster 11 feature), and top 80 number of Y (cluster 28 feature) when the full human IDRome is sorted (24,508 IDRs) are shown. F. Schematics of the canonical BAF and the MLL4 compass-like complexes. Protein subunits are colored if an IDR within that subunit is within the top 80 when the full human IDRome was sorted for number of M, cumulative Q Patch length, or number of Y. Striped subunits imply IDRs within that subunit have multiple top 80 grammar features. The color bars provide annotations for the coloring scheme used to depict the exceptional IDRs.

The splicing process also has a temporal ordering and so we examined if the earliest spliceosome complexes are enriched in top scoring R-patches, which is the defining feature of cluster 26 IDRs. The most upstream spliceosome complexes house some of the longest cumulative R-patches in the full human IDRome (**Figure 5C**). These include serine and arginine rich splicing factors (SRSFs), SNRNP70 of the U1 small nuclear ribonucleoprotein, U2AF1 of the spliceosomal E complex, and RBM25, DDX46, and CCAR1 of the spliceosomal A complex (**Figure 5D**) ^115,123^. These results suggest that long stretches of R-rich regions, the defining feature of cluster 26 IDRs, are important for the function of the earliest spliceosomal processes.

Next, we analyzed specific complexes that are critical for chromatin and gene regulation. Two trithorax (Trx) family complexes that play critical activating roles include the mammalian SWI/SNF, specifically, canonical BAF (cBAF) chromatin remodeling complexes, and the histone modifying complex, MLL4 COMPASS-like complex^133,155,156^. We find that only the cBAF and COMPASS-like MLL complexes contain IDRs that score in the top 80 for the defining features of nucleoplasm-specific GIN clusters. These features are number of Met residues, cumulative length of Q-patches, and number of Tyr residues (**Figure 5E**) that define clusters 10, 11, and 28, respectively. In particular, cBAF components stand out as having exceptional grammars in the context of the full human IDRome. IDR1 of the cBAF subunit, ARID1A, has the 3^rd^ highest valence of Met residues, the 5^th^ highest valence of Tyr residues, and the 26^th^ longest cumulative length of Q patches in the human IDRome. Other subunits, including SS18 (6^th^ in valence of Met residues, 9^th^ in valence of Tyr residues), also show exceptional grammars (**Figure 5F**).

In terms of the cumulative lengths of Q patches, IDRs within the MLL4 COMPASS-like complex stand out as being exceptional. Specifically, KMT2D, PAXIP1, and NCOA6 house the 1^st^, 2^nd^, and 10^th^ longest cumulative Q patches in the human IDRome, respectively (**Figure 5F**). KMT2D and NCOA6 also show exceptional grammars in terms of the valence of Met residues (**Figure 5F**). ARID1A and KMT2D are among the ten most commonly mutated driver genes in cancer ^157,158^. Removal of ARID1A IDR1 or truncation of KMT2D IDR10 does not affect the assembly of BAF and MLL4 COMPASS-like complexes, respectively ^44,159^, however, both IDRs drive condensation and modulate heterotypic interactions of their respective complexes. These IDRs, which house exceptional grammars, appear to play important roles for the function of BAF and MLL4 COMPASS-like complexes.

### RNA Pol I and II house evolutionarily conserved exceptional and distinct IDR grammars

RNA polymerases are essential for transcription. RNA polymerase I (Pol I) catalyzes the transcription of rDNA in the nucleolus whereas RNA polymerase II (Pol II) catalyzes the transcription of mRNAs and regulatory non-coding RNAs in the nucleoplasm ^128^. Pol I and Pol II comprise 13 and 12 subunits, respectively, and share five core subunits ^132^ (**Figure 6A**). From the Pol I and Pol II specific subunits we extracted all IDRs of length ≥ 70 and non-linker IDRs of length ≥ 50. Pol I contains three subunit-specific IDRs namely, POLR1F IDR1, POLR1G IDR4, and POLR1A IDR1. The grammars of POLR1F IDR1 and POLR1G IDR4 map them onto cluster 23 and the grammar of POLR1A IDR1 maps it onto cluster 7. Proteins with IDRs from these clusters are enriched in the nucleolus where Pol I transcription takes place. In contrast, Pol II contains one subunit-specific IDR, POLR2A IDR1, which features an IDR from cluster 28.

**Figure 6:**
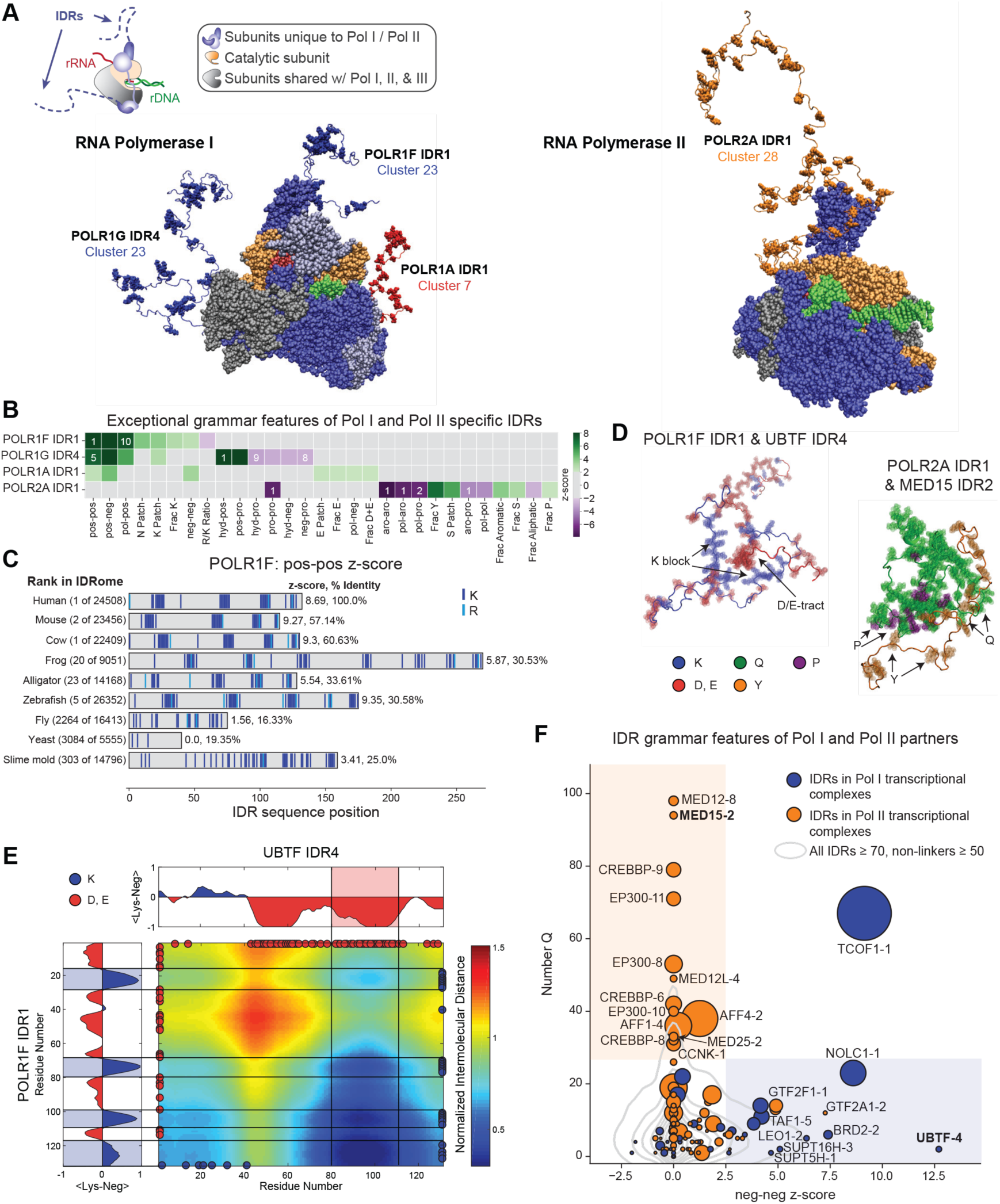
RNA Pol I and II house evolutionarily conserved exceptional and distinct IDR grammars. See Figure S6. A. Structures of RNA polymerase I and II (PDB 70b9 and 6exv, respectively) overlayed with representative conformations of all IDRs of length ≥ 70 and non-linker IDRs of length ≥ 50 amino acids, which are drawn from atomistic simulations. For Pol I F and G IDRs, lysine residues are shown, and negatively charged residues are shown for Pol I A. For Pol II IDR, aromatic amino acids are shown. B. Exceptional features of Pol I and Pol II specific IDRs. All features with |z-scores| ≥ 2 are shown. If a feature is in the top ten of the human IDRome, its rank is shown by the number in the box. C. Sequence features of POLR1F IDR1. On the left, the rank of the feature in the species-specific IDRome is given. On the right, the z-score of the sequence feature followed by the percent identity with the human IDR is shown ^92^. D. Representative snapshots are shown from all-atom simulations with pairs of molecules that were used to quantify interaction patterns, via scaled inter-residue distances, as shown in panel E. E. Normalized inter-residue distance maps extracted from atomistic simulations that model the association-dissociation equilibria of a pair of POLR1F IDR1 and UBTF IDR4 molecules. Inter-residue distances were normalized by an ideal prior such that blue colors imply enhanced attractions, and red colors imply enhanced repulsions when compared to pairs of ideal chains. The color bar provides an annotation of the normalized inter-residue distances. Distributions on the top and right show the average of the fraction of lysine residues minus the fraction of acidic residues over sliding windows of length five. This is used to highlight predicted charge-driven interactions. F. Complementary grammars of IDRs in Pol I and Pol II complexes^92,113,115,128,129,132^. Circle sizes are proportional to IDR length. Numbers after the gene name refer to the IDR number. Two-dimensional density plot for all IDRs of length ≥ 70 and non-linker IDRs of length ≥ 50 is shown by contours with a 0.05 threshold. Interactions of the bolded IDRs with Pol I and Pol II IDRs were assessed with atomistic simulations.

The subunit-specific IDRs of Pol I and Pol II have grammars with exceptional features that define clusters 23 and 28, respectively. The C-terminal POLR1F and the C-terminal POLR1G IDRs are 1^st^ and 5^th^ in the full human IDRome in terms of pos-pos patterning z-score (**Figure 6B**). This is a defining feature of IDRs from cluster 23 IDRs. Likewise, the C-terminal POLR2A IDR has the most well-mixed aromatic patterning (most negative aro-aro z-score) and the 3^rd^ highest number of Tyr residues in the full human IDRome, which are features that define cluster 28 IDRs (**Figure 6B**).

The case for the functional relevance of IDR grammars encoding exceptional features can be made via evolutionary analysis. We analyzed the IDRome of eight species and calculated pos-pos z-scores for IDRs from orthologs of POLR1F and POLR1G. We also analyzed the aro-aro z-scores for IDRs from orthologs of POLR2A. For POLR1F orthologs we find high pos-pos z-scores are conserved from humans to zebrafish with the POLR1F IDR being at least in the top 25 of all IDRs in each IDRome (**Figure 6C**). Interestingly, in this evolutionary range, the percent identity with human POLR1F IDR1 decreases to ∼30%. This suggests that while the exact sequence of POLR1F orthologs can vary considerably, the exceptional feature of high pos-pos z-score is maintained across evolution. Furthermore, we find a significant preference for Lys versus Arg residues in all POLR1F orthologous IDRs. Similar results were observed for the C-terminal IDR of POLR1G (**Figure S6A**). Interestingly, in yeast it is this IDR rather than the POLR1F IDR that has blocky patterning of positively charged residues.

For the C-terminal POLR2A IDR, all eight IDRs from the orthologous proteins have one of the most significant extents of uniform distribution of aromatic residues in their IDRomes (**Figure S6B**). In general, this IDR is more conserved than the Pol I IDRs with high percent identities compared to the human IDR. However, the length of the POLR2A IDR varies considerably across evolution. Overall, the conservation of exceptional grammars in RNA polymerase subunit specific IDRs suggests that these grammars play a role in the diverse functions of Pol I and Pol II.

Extant evidence suggests that subunit-specific IDRs can interact with IDRs found in pre-initiation and other transcriptional complexes ^48,139^. Additionally, truncation of the C-terminal IDR of the POLR2A homolog in yeast decreases the bound fraction of various pre-initiation complex components^160^. We hypothesized that certain IDRs with grammars that are enriched in nucleoli and the nucleoplasm should enable interactions with the Pol I and Pol II IDRs. To test this hypothesis, we performed atomistic simulations. The results from these simulations suggest that blocks of negative residues within an IDR found in Pol I transcriptional complexes make complementary interactions with the K blocks of POLR1F (**Figure 6E**). Likewise, glutamine residues within an IDR found in Pol II transcriptional complexes make complementary interactions with polar and aromatic residues within the POLR2A C-terminal IDR (**Figure S6C**). In contrast, repulsive or weakly attractive interactions are found when Pol I and II IDRs are analyzed with IDRs found in the orthogonal transcriptional complexes (**Figure S6D-S6E**).

To cast the simulations in terms of sequence-function relationships, we analyzed how the neg-neg z-scores correlate with the number of Gln residues in Pol I and Pol II transcriptional complex IDRs^92,113,115,128,129,132^ (**Figure 6F**). The contours show the two-dimensional distribution of all IDRs of length ≥ 70 and non-linker IDRs of length ≥ 50 in the human IDRome. This analysis suggests that the grammars of specific Pol I and II factors have exceptional and separable features in their IDRs. Specifically, IDRs that are specific to the Pol I-specific transcriptional complex have exceptional neg-neg z-scores. These include UBTF of the pre-initiation complex, TCOF1 and NOLC1 involved in rDNA organization, and LEO1, SUPT16H, and SUPT5H which are involved in transcription elongation. Likewise, IDRs that are specific to the Pol II transcriptional complex have exceptional Q valence. These include components of the Mediator complex and IDRs of the transcriptional co-activators EP300 and CREBBP.

### Cancer mutations disrupt exceptional IDR grammars within distinct GIN clusters

We hypothesized that disruption of IDR grammars, especially those pertaining to exceptional features, can modify IDR-driven protein targeting, interactions, and function to play key roles in disease. We tested this hypothesis by analyzing cancer driver genes ^161^. These genes encode proteins that are enriched in IDRs from cluster 10, cluster 11, and cluster 12 (**Figure 7A**). The association of cluster 12 with cancer driver genes is relevant given that blocks of negative and positive residues have previously been shown to localize proteins to bodies scaffolded by the Mediator component MED1^49^. The implication is that IDRs in proteins that drive cancer are non-random and may play a role in mediating nucleoplasmic specific interactions.

**Figure 7:**
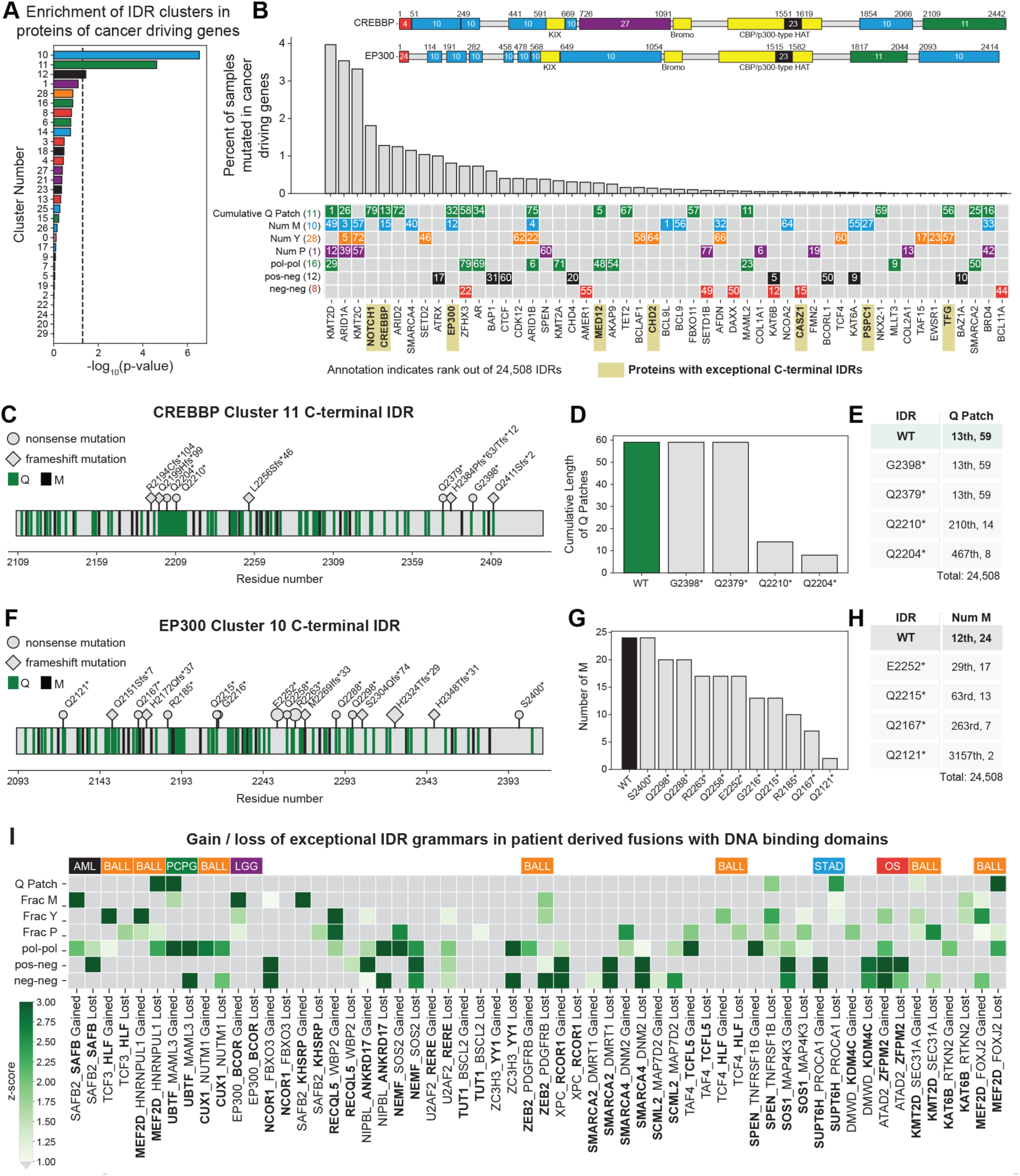
Cancer mutations disrupt exceptional IDR grammars within distinct GIN clusters. See Figure S7. A. Enrichment of GIN clusters in proteins of IntOGen cancer driver genes^161^ using the Fisher’s exact test. B. Within the IntOGen cancer driver genes^161^, genes encoding top 80 IDRs in terms of cumulative Q Patch, number of Met, Tyr, or Pro residues, and patterning of polar residues with respect to one another, patterning of oppositely charged residues, or the clustering versus mixing of negatively charged residues are shown. Colored boxes imply that a given protein contains a top 80 grammar feature, and the rank is annotated whereas the numbers in parentheses are the GIN cluster identities. Grey boxes imply that a top 80 IDR with that feature is not found in the protein. Inset: full protein sequence schematics of CREBBP and EP300. IDRs are colored and labeled by their GIN cluster. UniProt domains are shown in yellow. C. Sequence schematic of CREBBP C-terminal IDR. Marker size denotes number of patients with the mutation. D. Cumulative length of Q patches in WT CREBBP C-terminal IDR and truncation mutants. E. Rank and cumulative length of Q patches in WT CREBBP C-terminal IDR and truncation mutants. F. Sequence schematic of EP300 C-terminal IDR. Marker size denotes number of patients with the mutation. G. Number of methionines in WT EP300 C-terminal IDR and truncation mutants. H. Rank and number of methionines in WT EP300 C-terminal IDR and truncation mutants. I. Exceptional grammars of IDRs that are lost / gained in the Tripathi et al.^164^ patient derived fusion dataset were analyzed through the lens of the parent protein housing the DNA binding domain (bold). Cancer types associated with these fusions are shown across the top.

Next, we organized the cancer driving genes by their IDRs (**Figure 7B**). This shows that the IDRs in cancer driving genes have exceptional features. Specifically, 51 of the 619 cancer driver genes encode at least one of the top scoring IDRs (top 80 out of 24,508) in terms of cumulative length of a Q Patch, number of Met, Tyr, or Pro residues, patterning of polar residues with one another, patterning of oppositely charged residues with respect to one another, or the clustering versus mixing of negatively charged residues. Two exceptional IDRs encoded in cancer driver genes are the CREBBP C-terminal cluster 11 IDR and EP300 C-terminal cluster 10 IDR (**Figure 7B-7H**). Specifically, CREBBP C-terminal IDR is 13^th^ in cumulative length of Q patches (**Figure 7C**) and EP300 C-terminal IDR is 12^th^ in the number of Ms (**Figure 7H**). Both IDRs house cancer driving nonsense and frameshift mutations (**Figures 7C, 7F**). We find that nonsense mutations, which lead to truncations, cause a decrease in the exceptionality of the defining feature, such as cumulative length of the Q patch (**Figures 7D-7E**) or the number of Met residues (**Figures 7G-7H**). This reduction in the exceptionality of a specific feature is likely to have functional consequences that are disease-associated and merit closer scrutiny.

A more dramatic alteration of exceptional grammars comes from the subset of cancer driver genes, which are fusion oncogenes. Here, chromosomal translocations give rise to hybrid genes ^162–165^ known as fusion oncoproteins (FOs) that often link a structured DNA-binding domain of one parent protein with an IDR of another (**Figure S7A**). These types of FOs can modify the recruitment of transcriptional and / or chromatin remodeling complexes and lead to aberrant gene expression ^164,166^. Using the patient-derived dataset of Tripathi et al.,^164^ we identified IDR grammars and probed for the gain or loss of exceptional features upon fusion. This analysis was performed through the lens of the parent protein housing the DNA binding domain (**Figure S7A-S7B, 7I**). All 29 FOs examined lose and / or gain at least one non-random feature associated with IDRs in the nucleoplasm (Q Patch and fraction Met or fraction Tyr) or with unmodified cancer driver proteins (fraction Pro and patterning of polar, positive, and negative residues), thereby generating an aberrant IDR grammar (**Figure 7I**). This suggests that FOs are likely to rewire gene regulation protein interaction networks by swapping IDR grammars.

Next, we analyzed UBTF fusions, which are archetypes of grammars being altered and novel grammars being generated. These fusions are drivers of B-cell acute lymphoblastic leukemia (B-ALL)^167^ and pheochromocytomas and paragangliomas (PCC/PGL)^168^. In both cases, the exceptional C-terminal D/E-tract IDR of UBTF is replaced upon gene fusion and the formation of the chimeric protein (**Figure S7C**). Removal of the D/E-tract should weaken the driving forces for phase separation of UBTF with rDNA^48^. Furthermore, the D/E-tracts of UBTF can interact with the K blocks of POLR1F IDR1. Thus, replacement of the D/E-tract is likely to modulate the scaffolding behavior, interactions with POLR1F, and the DNA-binding activities which contribute to rDNA transcription^169^.

In B-ALL, UBTF forms an in-frame fusion with ATXN7L3. The ATXN7L3 portion encodes two IDRs, both of which are enriched in blocks of positively charged residues. Atomistic simulations show that the ATXN7L3 IDR2 interacts weakly with the POLR1F IDR, when compared to the UBTF IDR4 (**Figure S6F**). This points to a disruption of cognate interaction patterns upon alteration of the native IDR grammar. In PCC/PGL, UBTF forms a fusion with an IDR of the transcription co-activator MAML3 that is enriched in Q patches. Furthermore, in the context of the full human IDRome, ATXN7L3 IDR2 and MAML3 IDR are exceptional in pos-pos and Q patch z-scores, respectively (**Figure S7C**). Thus, one exceptional IDR grammar is replaced with a different exceptional IDR grammar upon gene fusion. These disruptions of the native grammar, appear to weaken cognate interactions, and generate aberrant interactions.

Next, we asked if there is evidence for IDR grammar swaps in FOs modifying partner recruitment. If so, we would expect genes that are functionally similar to FO parent genes to be enriched in IDR grammars that complement the grammars of the parent FO. We utilized the DepMap dataset^106^ to extract genes with correlated gene fitness effect profiles to UBTF and MAML3, the two FO parents for UBTF:MAML3. 213 genes have a Pearson’s correlation ≥ 0.1 with UBTF and 584 genes have a correlation ≥ 0.1 with MAML3 (**Figure S7D**).

The UBTF and MAML3 correlated gene sets are enriched for proteins with very different IDR grammars (**Figure S7E**). The UBTF set shows an enrichment in features such as FCR and K blocks when compared to the remaining human IDRome. Conversely, the MAML3 set shows a clear depletion of charged resides and enrichment in glycine, hydrophobic residues, and polar residues. The implication is that IDR-mediated interactions of UBTF and MAML3 are likely to be distinct. Based on this analysis, we hypothesize that creation of the UBTF::MAML3 fusion engenders a loss of interactions for UBTF through its cluster 7 IDR and a gain of very different interactions through the Q-tract features of the cluster 11 IDR of MAML3.

## Discussion

We have created a resource, which we refer to as GIN, that uncovers molecular grammars across all IDRs in the human IDRome and uses the grammars to organize IDRs into IDRome-spanning clusters. The complete dataset that includes the z-scores for the 90 sequence grammar features, as well as GO Slim and HPA classifications, can be found in **Table S1**. To facilitate the use of GIN as a resource, we have created two Google Colab notebooks. This should enable the usage of GIN for extracting IDR-specific grammars, the analysis of these grammars referenced to proteomes of interest, and the design of IDRs with preferred grammars.

Through the creation of the GIN resource, we discovered that cellular locations, functions, and processes are associated with specific GIN clusters. Our work organizes the IDRome by IDR grammars and uncovers how these grammars contribute to molecular functions, localization, and biological processes. We find that the highest scoring IDRs are housed in proteins that are involved in the earliest / core functions of a given location thus suggesting that IDR grammars may be more than just localization signals. Analysis of cellular fitness datasets highlight the potential importance of IDR-mediated contributions to functional dependencies between gene pairs that merit in-depth investigations.

Location- and function-specific grammars are consistent with results of previous studies ^18,65,67^. Our assessments of localization specificities enabled by the extraction of IDR grammars show stronger partition coefficients into nucleoli and nuclear speckles when compared to recent machine learning based efforts ^66^. There appear to be clear and discernible rules, which we have summarized, that determine how IDRs enable protein localization into membraneless bodies. We show how IDRs can alter the localization of proteins that share identical folded domains. This builds on previous work ^48^, which showed that the localization specificities proteins with similar IDRs are determined by the folded domains to which they are tethered. Taken together, we conclude that the synergies between IDRs and folded domains contribute jointly to the sub-cellular localization specificities of proteins.

We go beyond the contribution of IDR grammars to localization and use a GIN-driven analysis of proteins and complexes involved in specific molecular functions. We find that the grammars of IDRs likely enable complementary intermolecular interactions. We tested these predictions via simulations, which suggest that sequence-function relationships between IDRs might derive from the complementarity of IDR grammars. Additionally, we found that the IDRs with the highest scoring subcellular location specific grammars are housed in proteins involved in the core functions of that location. Our analysis suggests that the exceptional grammars of these IDRs specify function through complementary and location-specific IDR-mediated intermolecular interactions. Of particular interest are the IDRs of RNA polymerases I and II which share a majority of structural subunits but house the most exceptional location specific IDR grammars of the nucleolus and nucleoplasm, respectively.

Using GIN clusters as fingerprints for individual IDRs or a set of IDRs within a cluster will enable the design of mutations that allow for unmasking sequence-function relationships of specific IDRs in different contexts or of different IDRs across similar contexts. This is useful since it is often difficult to discern or identify sequence features of IDRs that are functionally important. The traditional approach of evolutionary analysis becomes challenging because extracting patterns of conservation and covariation from multiple sequence alignments ^2,15,16,170–174^ is confounded by large sequence variations of IDRs across orthologs and paralogs ^19,20,22,80^. Zarin et al., ^18,67^ have adapted heuristics from studies of sequence-ensemble relationships and combined these with models for mutational rates to develop methods that enable evolutionary analyses of IDRs. The organization afforded by GIN complements the approaches of Zarin et al. and can be used to generate testable hypotheses regarding determinants of sequence-function relationships of IDRs. While this has been done in recent one-off studies ^21,44,48^, we expect that the organization of the IDRome afforded by GIN will potentiate proteome- or sub-proteome-wide assessments of sequence-function relationships.

Considerable effort has been invested in studies that characterize conformational ensembles of IDRs, either as autonomous units or in the context of being tethered to folded domains^,3,5,6,9–11,17,21,26,29,31,34,40,53,58–60,70–73,75,77,78,81–86,175–203^. Conformations of an IDR can be characterized in terms of global measures such as average sizes and shapes, moments of distribution functions that quantify the amplitudes of spontaneous local and global conformational fluctuations, and the preferences for specific types of secondary and tertiary structural motifs ^10,11,22,29,40,56,71–73,75,82–85,175,178,181,183,192,203,204^. Conformational dynamics of IDRs can span a range of timescales ^176,180,182,195,196,205^, and these too are sequence-specific. The totality of these features emphasizes how conformational heterogeneity is a defining hallmark of IDRs ^17,81,83,86,87,175,184,185,206–211^.

There is interest in building on biophysical characterizations of conformational ensembles to codify the information within ensembles to generate sequence-ensemble relationships and use these to extract sequence-ensemble-function relationships^6,22,27,33,37,48,53,70,72,78,79,81,82,87,212,213^. Two recent efforts have focused on the codification of sequence-ensemble relationships across the entire human IDRome ^10,11^. From simulated ensembles, sequence-ensemble relationships can be derived by reducing sequence information to specific numerical parameters and the properties of conformational ensembles to numerical values such as scaling exponents ^10,11,71,84,202^. The dimensions of IDRs, measured in terms of radii of gyration (R_g_), for example, can be written as R_g_ ∼ N^ν^, where N is the number of residues in the IDR of interest, and ν is the scaling exponent ^84,192,202,203^. This parameter is used to infer the nature of IDR-solvent interactions and the length scale over which conformational fluctuations are correlated with one another ^84^. Larger values of ν (with 0.3 ≤ ν ≤ 1) are taken to imply favorable interactions with the solvent ^74^.

We used the deep learning-based prediction tool ALBATROSS developed by Lotthammer et al.,^10^ to assess whether GIN clusters, which are organized by distinct molecular grammars, are defined by cluster-specific values of the apparent scaling exponent ν. Irrespective of whether we used all IDRs (**Figure S7F**) or well mapped IDRs of length ≥ 70 and non-linker IDRs of length ≥ 50 (**Figure S7G**), we note that the ν values, sorted by GIN clusters, are not cluster-specific. The computed median ν values span a narrow range between 0.51 and 0.58, although the spread of ν values is cluster-specific, being largest for IDRs in clusters 28, 18, and 11. This analysis of sequence-ensemble relationships of IDRs with very different grammars suggests that the values of scaling exponents do not appear to show grammar specificity.

At this juncture, it is clear that while sequence-ensemble-function relationships can be extracted for a range of specific systems ^6,22,27,33,37,48,53,70,72,78,79,81,82,87,212,213^, this might be difficult to achieve on the proteome-scale, at least with the current descriptors being used for quantifying sequence-ensemble relationships. Instead, the current work makes it clear that considerable progress is likely to be made on understanding and manipulating sequence-function relationships of IDRs, and this is likely to advance rapidly through resources such as GIN.

## Methods

### Animal handling

*Xenopus laevis* (frog) oocytes were used for analysis of protein localization in nucleoli. Mature female *Xenopus laevis* frogs (3-7 years old) are housed in the professional animal facility. Oocyte harvesting is conducted by trained personnel, within the animal preparation room of this animal facility. All procedures are under the supervision of the Washington University Institutional Animal Care and Use Committee Office. All procedures are approved by the Washington University Institutional Animal Care and Use Committee (Animal Welfare Assurance #: A-3381-01). The animal care facility provides additional technical assistance where needed.

### Analysis of IDR grammars using NARDINI+

NARDINI+^20,44,48^ is an IDR sequence grammar analysis tool that combines the NARDINI (Non-random Arrangement of Residues in Disordered Regions Inferred using Numerical Intermixing) analysis with analyses of compositional grammar features. Together this yields a sequence specific z-score vector of 90 unique sequence features: 36 from NARDINI^20^ and 54 from compositional analyses. For the NARDINI patterning analyses, a given sequence is compared to 10^5^ random scrambles of its same composition to determine if a given patterning parameter is more well-mixed or blockier than by random chance. Here, residues are divided into eight groups: pol = (S, T, N, Q, C, H), hyd = (I, L, M, V), pos = (K, R), neg = (E, D), aro = (F, W, Y), ala = (A), pro = (P), and gly = (G). For two residue types U and X, we extract a U-X z-score. Z-scores less than zero imply the input sequence is more well-mixed than expected, whereas z-scores greater than zero imply the input sequence is blockier than expected.

For compositional features, z-scores are determined by comparing to the distribution from the species specific IDRome. In this paper, we analyzed the IDRomes of nine different species: Human (*Homo sapiens*, ID: 9606), Mouse (*Mus musculus*, ID: 10090), Cow (*Bos taurus*, ID: 9913), Frog (*Xenopus tropicalis*, ID: 8364), Alligator (*Alligator mississippiensis*, ID: 8496), Zebrafish (*Danio rerio*, ID: 7955), Fly (*Drosophila melanogaster*, ID: 7227), Yeast (*Saccharomyces cerevisiae*, ID: 559292), and Slim Mold (*Dictyostelium discoideum*, ID 44689). In all cases, proteomes were downloaded from the *Swissprot* database^92^. MobiDB was utilized to extract disordered regions from each sequence in the proteome^215,216^. A region is defined as an IDR if the consecutive consensus prediction is of length greater than or equal to 30. This yielded 24508 IDRs for human, 23460 IDRs for mouse, 22409 IDRs for cow, 9051 IDRs for frog, 14168 IDRs for alligator, 26351 IDRs for zebrafish, 16413 IDRs for fly, 5555 IDRs for yeast, and 14796 IDRs for slime mold. The tool localCIDER^34^ was utilized to calculate most compositional features. Specifically, the compositional features extracted were the fraction of each amino acid type (20 features); the fraction of polar, aliphatic, aromatic, positive, negative, charged residues (FCR), chain expanding residues (FCE), disorder promoting residues (8 features); the ratios of Ks to Rs and Es to Ds (2 features); the net charge per residue (NCPR), the hydrophobicity, the isoelectric point, and the polyproline-II propensity (4 features). We also analyzed “Patch” features which denotes the fraction of the IDR comprised of patches of a particular amino acid or arginine-glycine (RG) pairs (20 features). Specifically, a patch is defined by having at least four occurrences of the given residue or two occurrences of RG and could not extend past two interruptions. W patches are excluded since none exist in the human IDRome. For a given sequence, the specific compositional feature z-score is determined by utilizing the mean and standard deviation from the entire species specific IDRome

### K-means clustering of IDRs based on NARDINI+ z-score vectors

The z-score vectors from NARDINI+ were extracted for all human IDRs of length 100 ≤ *n* ≤ 300 (4529 IDRs, ∼18% of the human proteome). Seventy-nine percent of IDRs are below this cutoff and ∼2% of IDRs are above this cutoff. This length cutoff was used to minimize length dependent effects, particularly in the z-scores that depend on fractions of particular amino acids, as well as to focus on long IDRs that are more likely to play functional roles. In order to avoid large distances between z-score vectors mostly based on one or a few large z-score values, all z-scores > 3 were set to 3 and all z-scores < -3 were set to -3. A |z-score| cutoff of three was chosen since anything above three should be significant (p-value < 0.01).

To determine how many clusters the IDRs should be categorized into, we calculated the mean silhouette coefficient upon K-means clustering for cluster numbers ranging from 10 to 100 and five different initial random states. We found that the mean silhouette coefficient was generally flat regardless of number of clusters and initial random state chosen. Additionally, we found that the inertia started decreasing linearly around a cluster size of 30. Thus, we settled on a cluster size of 30 to allow for tractability.

Next, we categorized all remaining 19979 human IDRs into one of the 30 clusters. This was done by first restricting the maximum z-score value to 3 and minimum z-score value to -3, as was done for the human IDRs of length 100 ≤ *n* ≤ 300. Then the distance between each IDR z-score vector and each cluster centroid vector was calculated. The given IDR was then assigned to the cluster corresponding to the centroid with the smallest distance. Furthermore, we calculated the difference of the distance to the next best cluster centroid and the distance to the mapped cluster centroid. We refer to this value as the “Min Inter Clust Dist”. A large “Min Inter Clust Dist” value implies that the IDR is well mapped into its given cluster, whereas a small “Min Inter Clust Dist” implies that the IDR could be similarly mapped into another cluster. For analyses in which we only wanted to include well mapped IDRs, i.e., for Gene Ontology and Human Protein Atlas enrichment, a “Min Inter Clust Dist” cutoff of 1.5 was used.

Lastly, we compared GIN types derived from all IDRs of length 100 ≤ *n* ≤ 300 to GIN types derived from all IDRs of length *n* ≥ 30. K-means clustering was performed on this set of IDRs for a range of cluster sizes. Similarly to the IDRs of length 100 ≤ *n* ≤ 300 set, a GIN type size of 30 seemed reasonable based on mean silhouette coefficients and inertia. To determine if similar GIN types were extracted for each of these two IDR sets, the Euclidean distance between median z-score vectors was calculated for all GIN types derived from all IDRs of length 100 ≤ *n* ≤ 300 versus all GIN types derived from all IDRs of length *n* ≥ 30. Two GIN types were said to be similar if the Euclidean distance was ≤ 3. Twenty-three of the thirty GIN types derived from IDRs of length 100 ≤ *n* ≤ 300 mapped well to at least one GIN type derived from all IDRs of length *n* ≥ 30 (Figure S1B). Thus, in a majority of cases similar GIN types were derived regardless of which set of IDRs were used for K-means clustering.

### Enrichment of molecular functions, biological processes, subcellular locations, and KEGG pathways by GIN cluster

For molecular functions and biological processes, high-level Gene-Ontology parent terms (GO Slim) and their corresponding protein accessions were downloaded from https://go.princeton.edu/cgi-bin/GOTermMapper in June of 2023. Here, organism was set to Homo sapiens (GOA @EBI) and ontology was set to Generic slim. For subcellular location, all human protein atlas^96^ (HPA) subcellular locations and their corresponding protein accessions were downloaded from https://www.proteinatlas.org/about/download in June of 2023. Specifically, only main locations were extracted. For KEGG pathways^121^ the KEGG API was utilized in March 2024 to extract pathway information (www.kegg.jp./kegg/rest/keggapi.html). Then, we determined whether proteins housing IDRs were enriched in each of the molecular functions, biological processes, and subcellular locations using the Fisher’s exact test. Specifically, only the subset of proteins that were both in the original downloaded human proteome and either the full list of GO Slim molecular function, GO Slim biological process, or HPA subcellular location proteins were used. This criterion was utilized to ensure that we were only analyzing proteins for which functional or localization information has been gathered to ensure a protein was not missing a categorization just because it had not been studied. Overall, these criteria yielded 14016 total proteins analyzed for molecular function, 16570 total proteins analyzed for biological process, and 12887 total proteins analyzed for subcellular localization. Furthermore, a protein was said to house an IDR if an IDR of length ≥ 70 or a non-linker IDRs of length ≥ 50 existed within the protein. This choice was made as IDRs of these lengths should be more likely to confer functional and localization preferences in the context of the full protein.

For enrichment of molecular functions, biological processes, subcellular location, and KEGG pathways by specific GIN clusters, we also used an additional criterion for what defines a cluster specific IDR. Here, the “Min Inter Clust Dist” (d_min_) had to be ≥ 1.5 to ensure that the IDR was well-mapped onto the assigned GIN cluster (8560 IDRs). Again, we used the Fisher’s exact test to determine what functions, processes, locations, and pathways were enriched in a given GIN cluster. Here, only the subset of IDRs that had a length ≥ 70 or a non-linker IDRs of length ≥ 50 and a d_min_ ≥ 1.5, as well as their proteins were in either the full list of GO Slim molecular function, GO Slim biological process, HPA subcellular location, or KEGG pathway proteins were used. Thus, 5955 IDRs were analyzed for molecular function, 7058 IDRs were analyzed for biological process, 6619 IDRs were analyzed for subcellular location, and 7736 IDRs were analyzed for KEGG pathway.

### *Xenopus laevis* localization studies

Harvested oocytes are manually de-flocculated with forceps then subjected to collagenase digestion (gentle rocking) for 2 hours at 22°C. Oocytes are stored in ND96 Buffer (0.005M HEPES, 0.096M NaCl, 0.002M KCl, 0.0018M CaCl2, 0.001M MgCl2) that was filter sterilized and supplemented with 2.5mM Sodium Pyruvate C3H3NaO3 (Thermo 11360070) and 1x Penicillin-Streptomycin (Sigma P4333) at 18°C. Healthy stage 6 oocytes were selected, residual vasculature removed, and injected with messenger RNAs (mRNAs) using glass micro-needles (Drummond 3-000-203-G/X) and a Drummond Nanoinject II (3-000-204). Messenger RNAs encoding a C-terminally RFP tagged RNA polymerase I E subunit (PolIe-RFP) and mRNAs encoding an N-terminally GFP tagged protein of intertest (GFP-POI) were injected in a single oocyte. Each injection comprises 23 nLs of mRNA in ddH2O corresponding to 5 ng each of PolIe-RFP mRNA and GFP-POI mRNA. Injected oocytes were stored individually in wells in a 48-well Polystyrene SterileTissue Culture Plates (Fisher FB012930) in supplemented ND-96 buffer at 18°C for 24 hours to allow expression and localization of the exogenous proteins. Immediately prior to imaging, germinal vesicles (i.e. nuclei) were manually dissected in mineral oil and mounted on a glass slide with 6 µL of mineral oil. A 22 x 22mm glass coverslip was gently overlaid onto the sample and fields of view nucleoli or nuclear speckles were then immediately imaged. This procedure was carried out for all proteins of interest for at least two separate harvests of oocytes and similar results were obtained across these biological replicates.

Fields of view with nucleoli or nuclear speckles were imaged as a confocal Z-stack. All images shown are of a single Z-slice that is representative and is at least 2 µm above the coverslip. Demarcated sub-phases were used to identify which sub-phase contains the peak signal of the target protein.

Confocal imaging of all samples was carried out on a Nikon Eclipse Ti2 microscope with a Yokogawa CSU X1 disk module and a LunF laser launch equipped with 405 nm, 488 nm, 561 nm, and 647 nm lasers. Images shown were taken using a 63X, 1.4 NA Apo oil immersion objective (Nikon) and a Hamamatsu Orca Flash 4.0 CMOS camera. All experiments were carried out at room temperature. NIS-Elements software was used for all image acquisition. Images within a data set were taken with identical imaging parameters ensuring that signal was not saturated (averaging, binning, and projecting were not used). Images were taken and saved as 16-bit ‘.nd2’ files. ImageJ, Python3, and MATLAB were used for all quantitative image analysis.

Individual crops of fields of view with several nucleoli were Otsu thresholded in the PolR1e channel to create a mask corresponding to the location of a given nucleolus. Nuclear speckles were identified by their characteristic morphology using brightfield microscopy, namely their roughly unform size (∼2µm) and their tendency to be attached to a larger Cajal Body. Mask corresponding to the locations of nuclear speckles were made by Otsu thresholding the GFP-POI signal for bodies that were morphologically identified as nuclear speckles. Using the analyze particle and particle manager tools on ImageJ, masks were used to obtain the mean intensity value (AU) the GFP-tagged protein of interest channels. The mean intensity values of the GFP-POI channel were used to obtain partition coefficients (PCs). PCs are the difference of the GFP signal in the masked regions divided by the GFP signal in the non-masked regions (i.e. background or nucleoplasm). All images shown are representative crops of one or a few nucleoli or nuclear speckles and the brightness and contrast have been optimized.

### Analysis of RNA and DNA binding associated proteins

We extracted the FCR for all 8560 IDRs with a length ≥ 70 or a non-linker IDRs of length ≥ 50 and a “Min Inter Clust Dist” ≥ 1.5. We further categorized each IDR as to whether they belonged to a protein in the GO Slim molecular function category “RNA binding” or “DNA binding”. Lastly, FCRs of IDRs belonging to proteins in the GO Slim molecular function categories “RNA binding” and “DNA binding” were collected and split by their GIN cluster. To determine if the FCR distributions were significantly different between different subsets of IDRs the Mann-Whitney test was performed.

### Extraction of functional relationships between GIN clusters using DepMap

Information regarding functional relationships between GIN clusters was extracted from the DepMap 24q4 dataset^106^. The dataset contains gene fitness effect scores for thousands of genes, generated by knocking out each gene one at a time using CRISPR and measuring the fitness effect of the knockdown on > 1000 cancer cell lines. Two genes are said to be functionally linked if their effects on cellular fitness are correlated. Genes were filtered to include only those in which valid gene effect scores were determined for at least 1000 cell lines (n=17787). Correlation was assessed by calculating the Pearson correlation coefficient of gene effect profiles between all gene pairs. For each gene, the correlation coefficient for every other partner gene was ranked, and this ranking was symmetrized by taking the minimum ranking for a given gene pair. An undirected edge was then constructed between two different gene nodes if the correlation rank was less than the stated threshold for a given figure.

To measure global associations between GIN clusters, all well mapped IDRs of length ≥ 70 and non-linker IDRs of length ≥ 50 were mapped to its gene node. Then, edges between genes were counted in order to convert between gene nodes and GIN cluster nodes. Here, edges were drawn between GIN clusters X and Y if the correlation rank for the given gene pair was less than the given threshold and one gene encodes an IDR mapped to cluster X and the other an IDR mapped to cluster Y. This is repeated for each pair of unique IDRs in the given gene pair. The total number of edges between two GIN clusters was then normalized by the number of combinations of member genes. This procedure reduces the order of the original graph, defined by the number of genes, down to the number of GIN clusters. This is because multiple genes map onto a single GIN cluster. Each node in the reduced graph represents an agglomeration of gene nodes, and an individual gene contributes to multiple nodes in the lower order graph.

### GIN cluster IDR enrichment in location specific biological processes

For each protein in the list of location specific biological processes (Table S4) we determined whether that protein contained an IDR of length ≥ 70 or a non-linker IDRs of length ≥ 50 and whether that IDR was categorized to a location specific GIN cluster. D3blocks was utilized to create Sankey plots (github.com/d3blocks). Here, the first layer corresponds to the general biological process and width of the edges connecting to the discrete biological processes (second layer, grey) correspond to how many proteins are part of the given discrete biological process. The width of the edges between the discrete biological processes and IDR type correspond to the number of IDRs of the given GIN cluster in the given discrete biological process. IDRs that are not in the given set of location specific GIN clusters are only shown if the full list of proteins in the discrete biological process contain no location specific GIN cluster IDRs. This choice was made so that way the size of the grey boxes in the second layer corresponds to the total number of proteins in the discrete biological process.

To determine if the location specific GIN cluster IDRs are enriched in location specific discrete biological processes, we determined the fraction of location specific discrete biological processes that contain at least one location specific GIN cluster IDR. For the nucleolus and nucleoplasm, we then compared this fraction to the fraction extracted for all other subsets of three GIN cluster IDRs that do not include the location specific GIN clusters. For nuclear speckles the same analysis is done except all other subsets of two GIN clusters IDRs are analyzed. This is because nuclear speckle IDRs are enriched in only two types of GIN clusters.

### Analysis of localization specific GIN cluster IDRs

For z-score sorted IDR plots all IDRs with a length ≥ 70 or a non-linker IDRs of length ≥ 50, a d_min_ ≥ 1.5, and belong to the given GIN cluster are shown. HeLa cell proteome wide abundance values were collected from Nagaraj et al.^217^. Proteins involved in specific processes of ribosome biogenesis, splicing, and gene regulation were collated from various sources and are listed in Tables S4^92,113,115,116,125,127–129,131–133,142–144^.

### Identification of top IDRs in the full human IDRome

For identification of top IDRs in the full human IDRome, all IDRs (length ≥ 30) of the human IDRome are sorted by a particular grammar feature (24508 IDRs). For determining whether general / discrete processes have top IDRs of a given grammar feature, we examined all proteins in each general / discrete process and counted the number of proteins that contained at least one IDR of the given grammar feature in the top 80 of the human IDRome. The choice of top 80 was made to allow for easy color scaling in our schematics showing top IDRs in early processes / important functional complexes.

### Atomistic simulations

All atom simulations were performed using the ABSINTH^218^ implicit solvation model and forcefield paradigm as implemented in the CAMPARI simulation engine (http://campari.sourceforge.net). The parameter set was based on abs3.2_opls.prm with the radius of the sodium ions increased to 1.81 Å to improve sampling with highly acidic sequences. Simulations were performed in spherical droplets of radii 350 Å at 340 K with explicitly modeled counterions and an excess of 2.5mM NaCl.

Two types of simulations were performed. First isolated IDR simulations were performed on Pol I and II IDRs. For these simulations, five independent simulations were conducted with 10^7^ equilibration steps and 5.15x10^7^ production steps. Since, the POLR1A IDR1 is a linker IDR, a harmonic restraint was applied between the first and last Cα atoms to restrain this distance to the distance observed in the Pol I structure (7ob9.pdb)^219^. Here, the distance was set to 37 Å and the spring constant to 10 kcal mol^-1^ Å^-2^.

The second type of simulations performed were two molecule IDR simulations. In all simulations a harmonic distance restraint was applied between central Cα atoms in the two IDRs in order to restrain their distance if it exceeded the target maximum distance. The maximum distance was set to 100 Å for Pol I and II IDR simulations. To initialize the two molecule simulations, ten independent simulations with 10^5^ production steps were performed. Then the end PDBs from these simulations were used as starting structures for full simulations with 10^7^ equilibration steps and 3x10^7^ production steps. Additionally, three independent reference two molecule simulations were performed. Here, only the repulsive Lennard-Jones term is kept on with an exponent of six and a linear scaling factor of 0.001. We term this type of reference simulation as an effective Flory random coil. For the reference simulations, 5x10^6^ equilibration and 5x10^7^ production steps were performed.

To quantify the degree of interaction compared to the reference simulations distance maps were calculated using the SOURSOP analysis package using the get_interchain_distance_map() function^220^. Then the mean distance map from the full simulations was divided by the mean distance map from the reference simulations. Thus, values less than one imply enhanced interactions compared to the reference simulations. Interaction regions are related to the sequence grammars by calculating residue type profiles. Here, we consider two types of residues: set1 and set2. For a given sequence, we calculate the fraction of set1 residues minus the fraction of set2 for each sliding window of length five. Then, for each residue the values from all sliding windows that contain that residue are averaged to obtain a residue specific mean net fraction of set1 vs set2 residues.

### Pol I and II Evolutionary Analyses

NARDINI+ analyses were performed for eight additional species: Mouse (*Mus musculus*, ID: 10090), Cow (*Bos taurus*, ID: 9913), Frog (*Xenopus tropicalis*, ID: 8364), Alligator (*Alligator mississippiensis*, ID: 8496), Zebrafish (*Danio rerio*, ID: 7955), Fly (*Drosophila melanogaster*, ID: 7227), Yeast (*Saccharomyces cerevisiae*, ID: 559292), and Slim Mold (*Dictyostelium discoideum*, ID 44689)^92^. In some cases, Pol I and Pol II IDRs were not in the original accession list, not predicted from MobiDB, or were predicted to have small interruptions of ordered regions from MobiDB. In these cases, the IDRs were added manually using a combination of MobiDB predictions^215^, AlphaFold predictions^2^, and information from alignments^92,221^. These sequences and their corresponding NARDINI+ values were then added to the full species IDRome for analysis. Percent identities with the human IDRs were extracted from UniProt after multiple sequence alignment of all nine species IDRs.

### IDR grammar features of Pol I and Pol II partners

For Pol I complex partners, all proteins that participate in rDNA transcription as listed by Dörner et al.^113^ (Table S4) but were not Pol I subunits were extracted. For Pol II complex partners, all proteins that were categorized under the general process “RNA polymerase II-mediated transcription” but were not Pol II subunits were extracted (Table S4). Then, the IDRs of length ≥ 70 and non-linker IDRs of length ≥ 50 were extracted for each of these proteins lists and their properties were compared to the remaining IDRome. For Pol I partner IDRs neg-neg z-scores were enriched compared to the remaining IDRome. For Pol II partner IDRs number Q z-scores were enriched compared to the remaining IDRome. Thus, given these enrichments and that these features are predicted to engage in complementary interactions with the blocky positive and well-mixed aromatic patterning of Pol I and Pol II IDRs, respectively, we plotted neg-neg z-score vs number Q z-score for all Pol I and Pol II partner IDRs.

### Enrichment of GIN clusters in cancer driver genes

The list of cancer driver genes was extracted from IntOGen^161^ (https://www.intogen.org/ <u>search</u>). Using this gene list the enrichment of IDRs in each GIN cluster was determined using a Fisher’s exact test. In Figure 7B, only cancer driver genes encoding at least one top 80 IDR in the features listed are shown.

### Cancer driving mutations of CREBBP and EP300

Driver mutations were extracted from cBioPortal (https://www.cbioportal.org/). Driver mutations were selected by filtering for mutations with Annotation = OncoKB Likely Oncogenic. For CREBBP and EP300 C-terminal IDRs all nonsense and frameshift mutation following this filtering are shown in Figures 7C and 7F. Marker sizes are proportional to the number of patients with the given nonsense / frameshift mutation.

### Gain / loss of exceptional IDR grammars in patient derived fusions with DNA binding domains

The FOdb-II database of 3174 fusion oncogenes were extracted from Tripathi et al.^164^. These fusions had verified data on cancer type and number of patient occurrences. This fusion list was then filtered for DNA binding domain (DBD) containing proteins. Briefly, the fusion sequences were subject to analysis using RPS-BLAST to extract DBDs using the PFAM and CDD domain list generated in King et al.^48,222^. This filtering generated 451 fusion oncogenes with DBDs. Next, the full sequences of the parent proteins were extracted from UniProt ^92^ and the longest intersecting sequences with the fusion were extracted for each parent protein. Only fusions in which both parent sequences contributed to at least 50 residues in the fusion were kept. Then, the fusion sequences were further filtered for fusions in which only one parent had a DBD(s), parent_DBD_. This filtering was performed in order to focus on IDR changes. Next, the fusions were filtered to find fusions in which the parent_DBD_ lost at least one IDR upon fusion, quantified by having less than 25% of an IDR sequence still in the fusion sequence, and the parent_DBD_ gained at least one IDR from the non-DBD parent protein. Twenty-nine fusions satisfied all of these criteria. Lastly, the neg-neg, pos-neg, pol-pol, Frac P, Frac Y, Frac M, and Q Patch z-scores were extracted for all lost and gained IDRs. These grammar features were chosen given their enrichment in IDRs in nucleoplasmic proteins and cancer driver proteins. If more than one IDR was gained / lost than the z-score recorded was the maximum of all the IDRs gained / lost.

### DepMap 24Q2 fitness correlation analyses of UBTF and MAML3

Pairwise fitness correlation values were downloaded from DepMap for UBTF and MAML3 for 1150 cancer cell lines in August 2024 (CRISPR (DepMap Public 24Q2+Score, Chronos), https://depmap.org/portal/interactive/custom_analysis)^106^. Positively correlated genes were kept if they had a Pearson correlation of ≥ 0.1 to our gene of interest. This generated a set of positively correlated fitness genes. The proteins of these genes were then analyzed for their IDR grammars. Specifically, z-score distributions for each of the 90 grammar features of the IDRs within the positively correlated fitness set were compared to z-score distributions of the remaining human IDRome. This generated a signed log_10_(p-value) using the two-sample Kolmogorov-Smirnov test. Here, the log_10_(p-value) was positive if the p-values was < 0.05, the mean of the z-scores for the set was greater than the mean of the z-scores of the remaining IDRome and greater than zero. The log_10_(p-value) was negative if the p-values was < 0.05, the mean of the z-scores for the set was less than the mean of the z-scores of the remaining IDRome and less than zero. All other values were set to zero. These choices were made to distinguish whether a significant change in distribution was due to an increase or decrease in z-score values.

### Relationship of GIN clusters to apparent scaling exponent

To extract apparent scaling exponents for all human IDRs the ALBATROSS Google Colab was utilized (https://colab.research.google.com/github/holehouse-lab/ALBATROSS-colab/ blob/main/example_notebooks/polymer_property_predictors.ipynb)^10^. Then, distributions of the apparent scaling exponents were shown by GIN cluster in Figures S7F-S7G.

### Resource Availability and Access

The Google Colab notebooks, code, and data that provides full access to GIN is available via https://github.com/kierstenruff/RUFF_KING_Grammars_of_IDRs_using_NARDINI-.

## Acknowledgments

We thank Anthony Hyman, Tanja Mittag, and Pappu lab members and alumni especially Megan Cohan, Rahul Das, Jeong-Mo Choi, Mina Farag, Alex Holehouse, and Dylan Tomares for helpful discussions and insights regarding molecular grammars. This work was funded by the US Air Force Office of Scientific Research grant (FA9550-20-1-0241 to RVP), the St. Jude Research Collaborative on the Biology and Biophysics of RNP granules (to RVP), the US National Science Foundation (MCB-2227268 to RVP), the US National Institutes of Health (F32GM146418-01A1 to MRK, K99GM152778 to MKS, and R01NS121114 to RVP), and the Howard Hughes Medical Institute (to CK).

## Author Contributions

Conceptualization: KMR, MRK, and RVP; NARDINI+ Methodology: KMR, MRK, AWY, VL, MKS, CK, and RVP; Creation of GIN: KMR; *Xenopus* Measurements: MRK, AP; Curation of datasets and analyses: KMR, MRK, VL, AWY, WEL; Critical Assessments: KMR, MRK, XS, CK, RVP; Funding acquisition: CK, RVP; Project administration: KMR, CK, and RVP; Supervision: KMR, CK, and RVP; Preparation of Figures: KMR, MRK, AP, AWY, CK, and RVP; Writing: KMR and RVP; Reviewing, additions, and editing: All authors.

---

**Figure S1:**
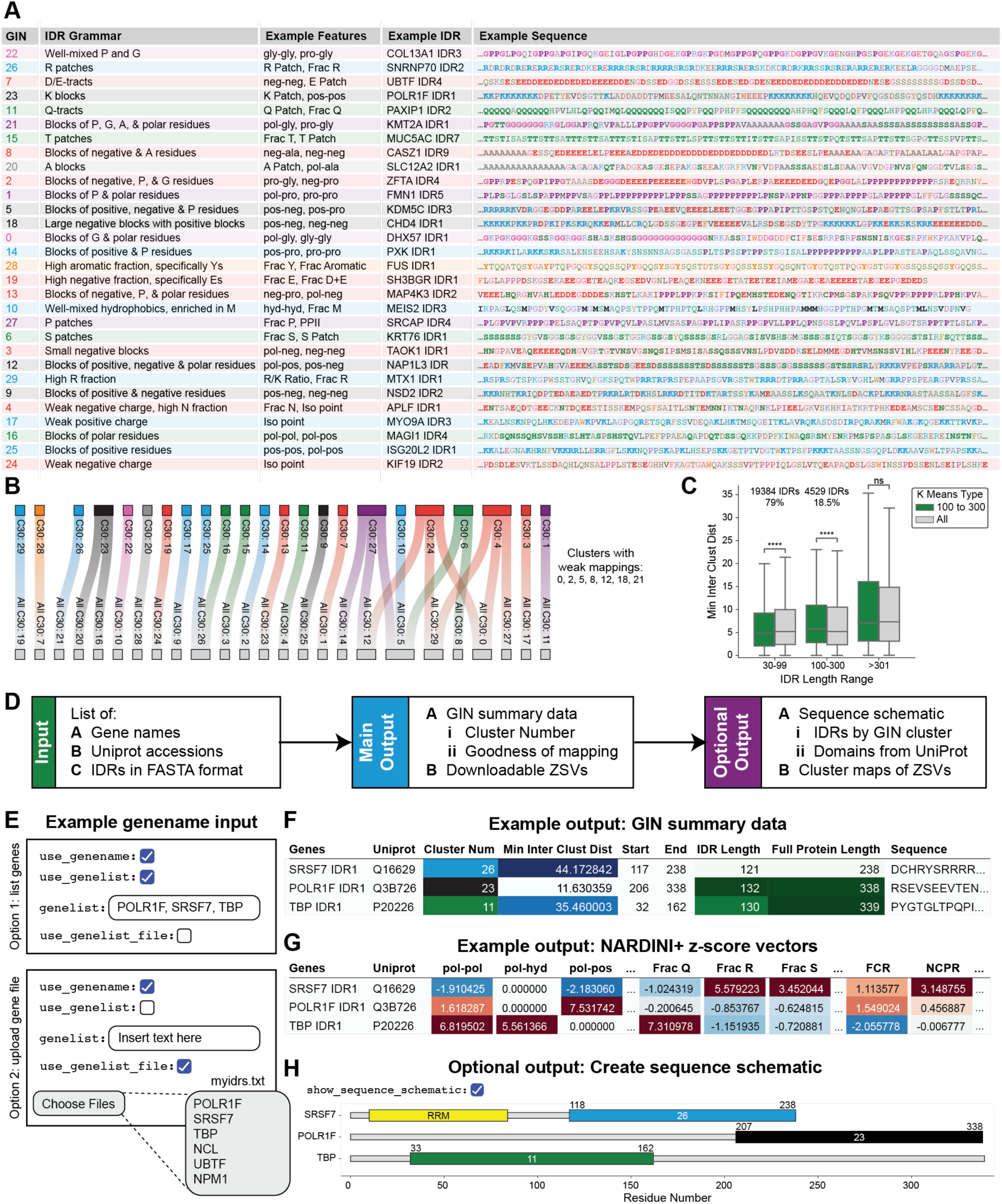
Description and test of GIN clusters, related to Figure 1. A. GIN clusters sorted by their M(d_min_) values which provides a goodness of the cluster. Here, the IDR grammar is described, and exceptional grammar features are listed for each GIN cluster. Additionally, an example IDR and its sequence is shown for each GIN cluster. B. Mapping of GIN clusters identified using all IDRs of length 100 ≤ *n* ≤ 300 (top) to GIN clusters identified if all IDRs ≥ 30 were used (bottom). Lines are drawn if the Euclidean distance between the medians ZSVs of the two clusters was ≤ 3. Similar GIN clusters are identified in both cases. C. Box and whisker plots comparing the d_min_ distributions for IDRs of various length ranges depending on whether the k-means clustering was used to extract GIN clusters on IDRs of length 100 ≤ *n* ≤ 300 (green) or all IDRs ≥ 30 (grey). Here, **** signifies p-values that are ≤ 10^-4^ computed via a paired t-test. D. General workflow of GIN resource in Google Colab notebooks. E. Example Google Colab notebook input parameters that allows users to input a comma separated gene list (Option 1) or a file of gene names (Option 2). F. Example GIN output provided by the input gene list of POLR1F, SRSF7, and TBP. Here, the Min Inter Clust Dist (see methods) provides a measure of how well the given IDR maps to the identified GIN cluster. G. The notebook also outputs the ZSVs for each of the IDRs in the gene list. These can be downloaded as a .csv or .excel file for further analyses. H. Users have the option to create a sequence schematic of the genes in their gene list. Here, IDRs are colored and labeled by their GIN cluster. Domains, downloaded from UniProt, are shown in yellow with the domain name.

**Figure S2:**
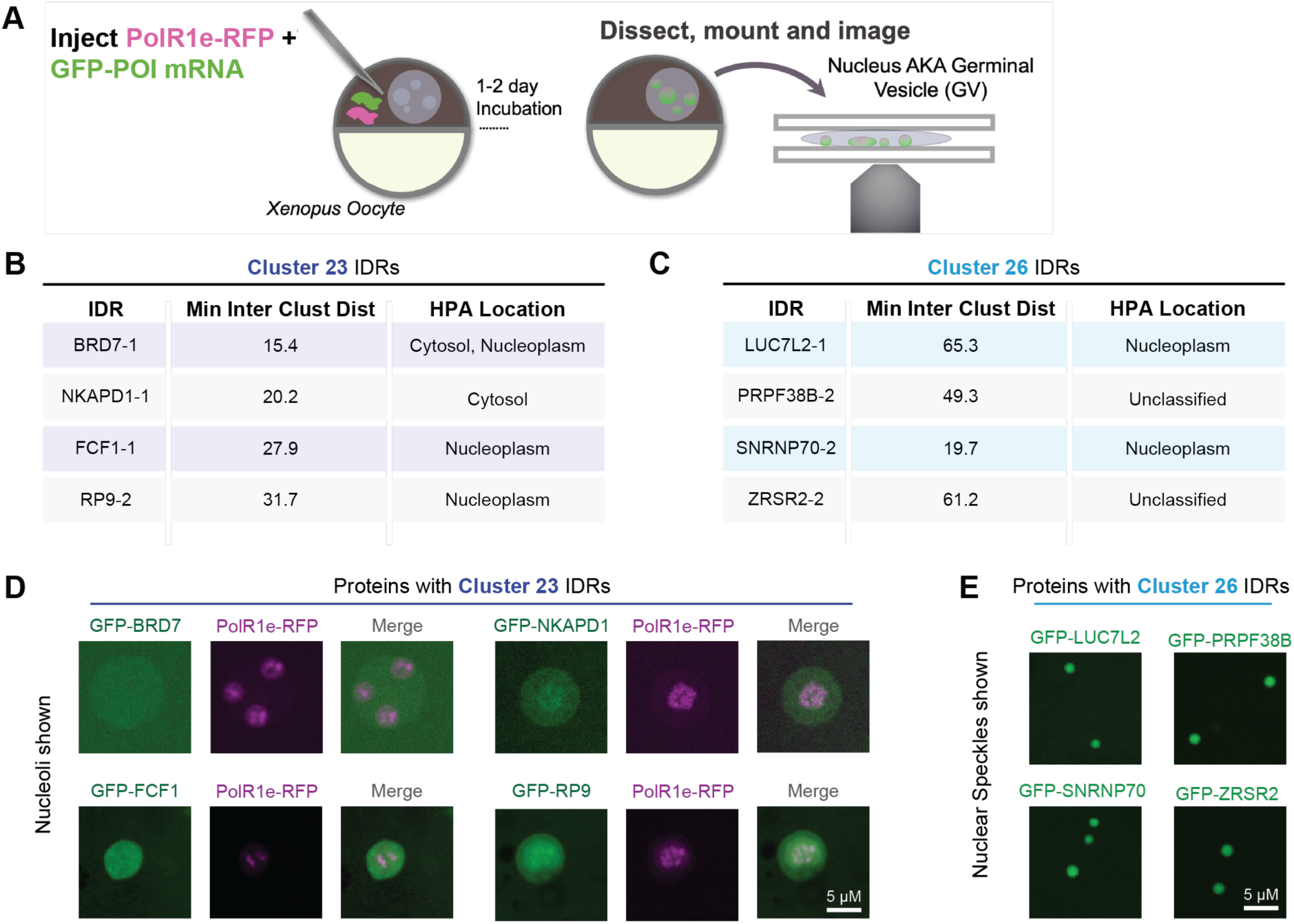
Distinct GIN clusters drive localization and are enriched in specific functions, related to Figure 2. A. Schematic of obtaining and imaging living nuclei (i.e. germinal vesicles) from *Xenopus* oocytes that express a given GFP-tagged protein of interest (GFP-POI) to identify their sub-nuclear localization. B. Table of cluster 23 IDRs whose full protein localization was analyzed. Here, Min Inter Clust Dist is a quantification of the goodness of mapping to that cluster (see Methods). C. Table of cluster 26 IDRs whose full protein localization was analyzed. Here, Min Inter Clust Dist is a quantification of the goodness of mapping to that cluster (see Methods). D. Representative confocal images of individual nucleoli in *Xenopus* oocyte GVs expressing PolRIe-RFP and a GFP-tagged protein that contains a cluster 23 IDR. E. Representative confocal images of nuclear speckles in *Xenopus* oocyte GVs expressing a GFP-tagged protein that contains a cluster 26 IDR.

**Figure S3:**
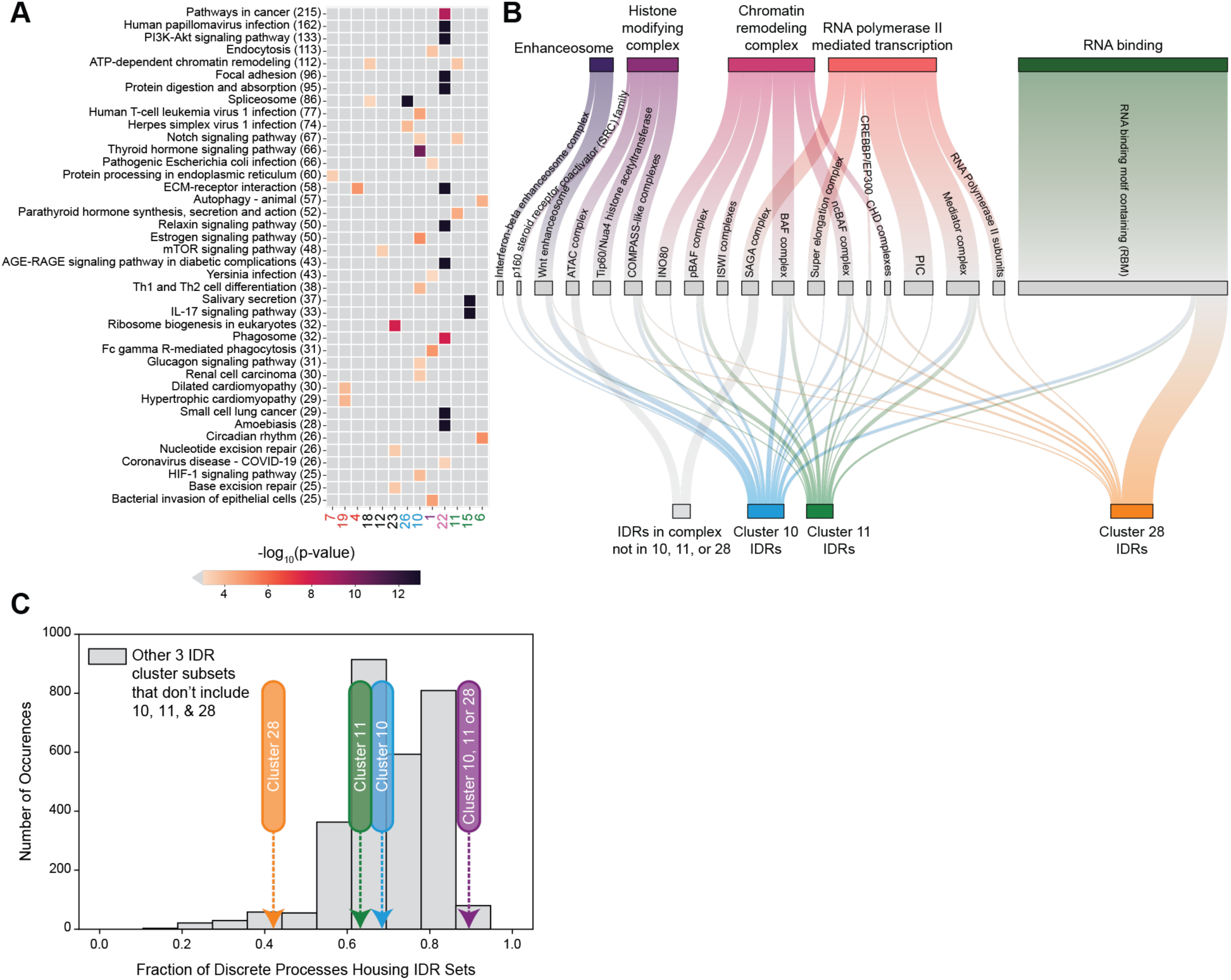
GIN clusters enriched in the nucleoplasm are associated with discrete complexes, related to Figure 4. A. Enrichment of GIN clusters in KEGG pathways^121^ (Fisher’s exact test). B. Sankey plot showing the relationship between discrete gene regulation processes and whether the proteins in those processes house nucleoplasmic specific GIN cluster IDRs (clusters 10, 11 and 28). Edge widths between the general biological processes and the discrete processes denote the number of proteins in the given discrete process. Edge widths between the discrete processes and the GIN cluster IDR type denote the number of IDRs in the discrete process that are categorized by the given GIN cluster. IDRs that are not part of GIN clusters 10, 11, or 28 are only shown if the discrete process contains no IDRs classified into GIN clusters 10, 11, or 28. General gene regulation processes with their corresponding discrete processes were collated from various sources in **Table S4**^92,115,125,127–129,131–133,142–144^. Visualization was created using D3Blocks (github.com/d3blocks). C. Comparison of the fraction of discrete processes housing different sets of GIN cluster IDRs. Here, 89% of discrete complexes have at least one IDR in cluster 10, 11, or 28 (purple arrow), compared to an average of 70% for any other triplet set of clusters excluding these clusters (grey histogram).

**Figure S4:**
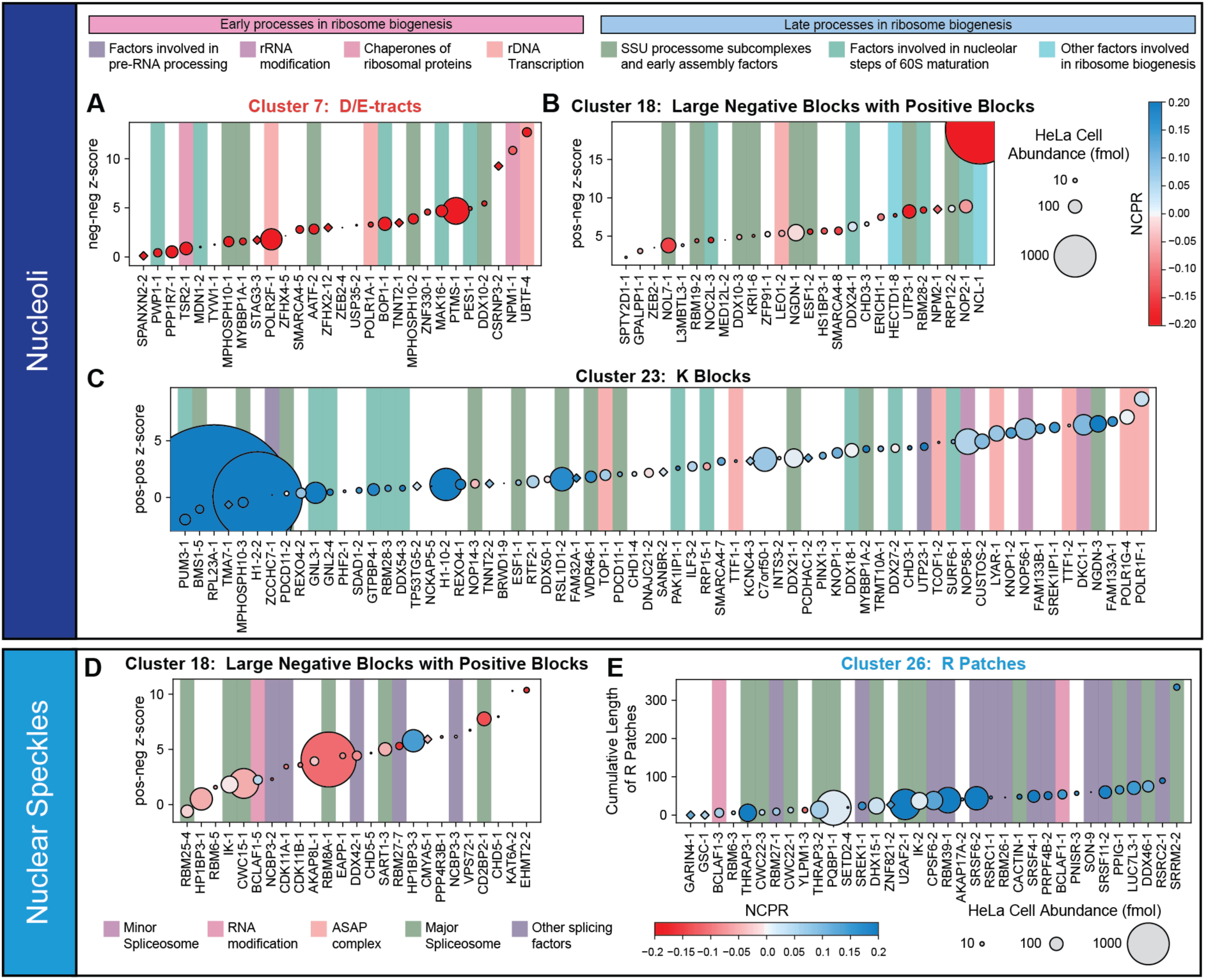
Nucleoli and nuclear speckle IDRs sorted by a prominent feature of the enriched GIN cluster, related to Figure 5. A. Cluster 7 HPA denoted nucleoli IDRs sorted by an enriched cluster 7 feature: neg-neg z-score. The number after the gene name refers to the IDR number. Circle size denotes HeLa cell abundance and diamonds denote that the abundance is not known for that protein. Circle color denotes the NCPR of the IDR. If a protein is involved in a ribosome biogenesis process, then this is denoted by the background bar color. Processes considered are rDNA transcription, rRNA modification, factors involved in pre-RNA processing, chaperones of ribosomal proteins, SSU processome subcomplexes and early assembly factors, factors involved in nucleolar steps of 60S maturation, and other factors involved in ribosome biogenesis. Early ribosome biogenesis processes are shown in pinks /purples, whereas late ribosome biogenesis processes are shown in blues /greens. B. Same as A, except cluster 18 HPA denoted nucleoli IDRs are sorted by an enriched cluster 18 feature: pos-neg z-score. C. Same as A, except cluster 23 HPA denoted nucleoli IDRs are sorted by an enriched cluster 23 feature: pos-pos z-score. Top scoring IDRs are involved in early ribosome biogenesis processes (pink /purple bars) D. Cluster 18 HPA denoted nuclear speckle IDRs sorted by an enriched cluster 18 feature: pos-neg z-score. The number after the gene name refers to the IDR number. Circle size denotes HeLa cell abundance and diamonds denote that the abundance is not known for that protein. Circle color denotes the NCPR of the IDR. If a protein is involved in a splicing process, then this is denoted by the background bar color. Processes considered are major spliceosome, minor spliceosome, RNA modification, ASAP complex, and other splicing factors. E. Same as D, except cluster 26 HPA denoted nuclear speckle IDRs are sorted by an enriched cluster 26 feature: cumulative length of R patches.

**Figure S5:**
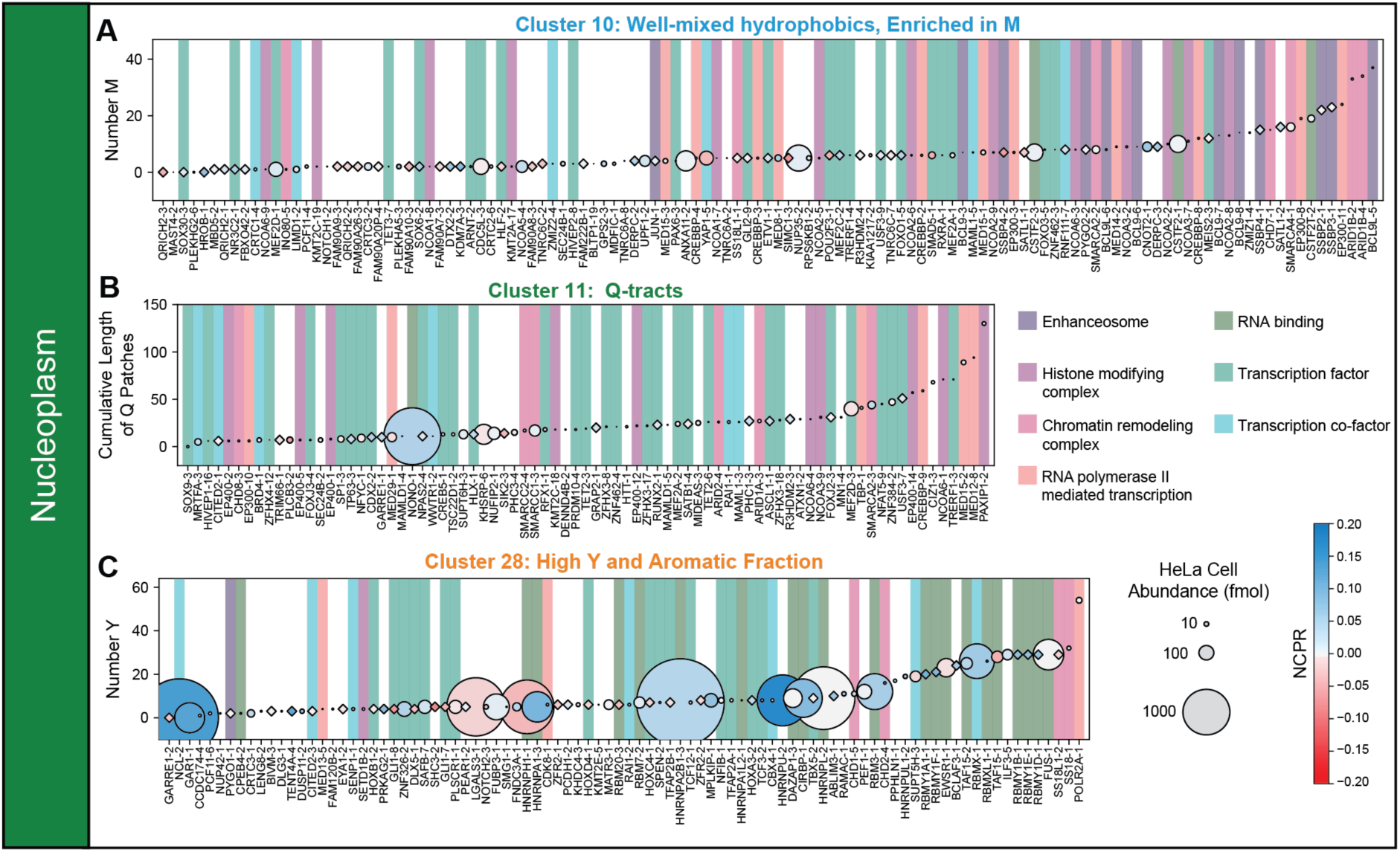
Nucleoplasmic IDRs sorted by a prominent feature of the enriched GIN cluster, related to Figure 5. A. Cluster 10 HPA denoted nucleoplasmic IDRs sorted by an enriched cluster 10 feature: number of M. The number after the gene name refers to the IDR number. Circle size denotes HeLa cell abundance and diamonds denote that the abundance is not known for that protein. Circle color denotes the NCPR of the IDR. If a protein is involved in a gene regulation process, then this is denoted by the background bar color. Processes considered are enhanceosome, histone modifying complex, chromatin remodeling complex, RNA polymerase II mediated transcription, RNA binding, transcription factor, and transcription co-factor. B. Same as A, except cluster 11 HPA denoted nucleoplasmic IDRs are sorted by an enriched cluster 11 feature: cumulative length of Q patches. C. Same as A, except cluster 28 HPA denoted nucleoplasmic IDRs are sorted by an enriched cluster 28 feature: number of Y.

**Figure S6:**
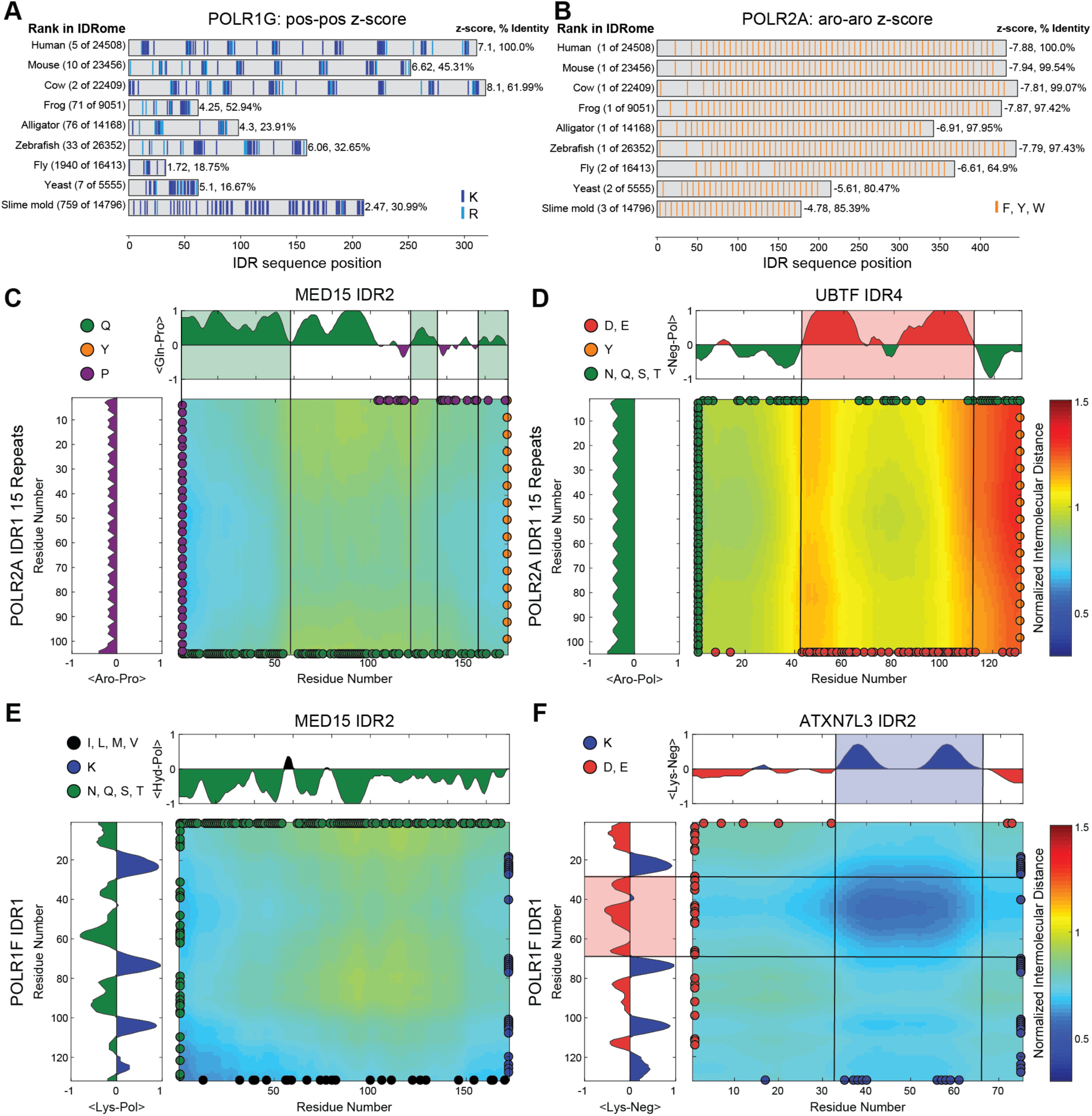
Exceptional Pol I and II IDRs are conserved and interact with distinct exceptional IDRs, related to Figure 6. A. Sequence grammar features of POLR1G IDR4. On the left, the rank of the grammar feature in the species specific IDRome is given. On the right, the z-score of the sequence feature followed by the percent identity with the human IDR is shown. B. Sequence grammar features of POLR2A IDR1. On the left, the rank of the grammar feature in the species specific IDRome is given. On the right, the z-score of the sequence feature followed by the percent identity with the human IDR is shown. C. Normalized inter-residue distance maps extracted from atomistic simulations that model the association-dissociation equilibria of pair of molecules comprising POLR2A IDR1 15 repeats and MED15 IDR2. Inter-residue distances were normalized by an ideal prior such that blue colors imply enhanced attractions, and red colors imply enhanced repulsions when compared to pairs of ideal chains. The color bar provides an annotation of the normalized inter-residue distances. Distributions on the top and left quantify the average of the fraction of residue set 1 minus the fraction of residue set 2 across sliding windows of five residues. This is used to highlight enrichments of certain types of residues and whether these enrichments correspond to regions of interaction. D. Same as C but the pair of molecules is POLR2A IDR1 15 repeats and UBTF IDR4. E. Same as C but the pair of molecules is POLR1F IDR1 and MED15 IDR2. F. Same as C but the pair of molecules is POLR1F IDR1 and ATXN7L3 IDR2

**Figure S7:**
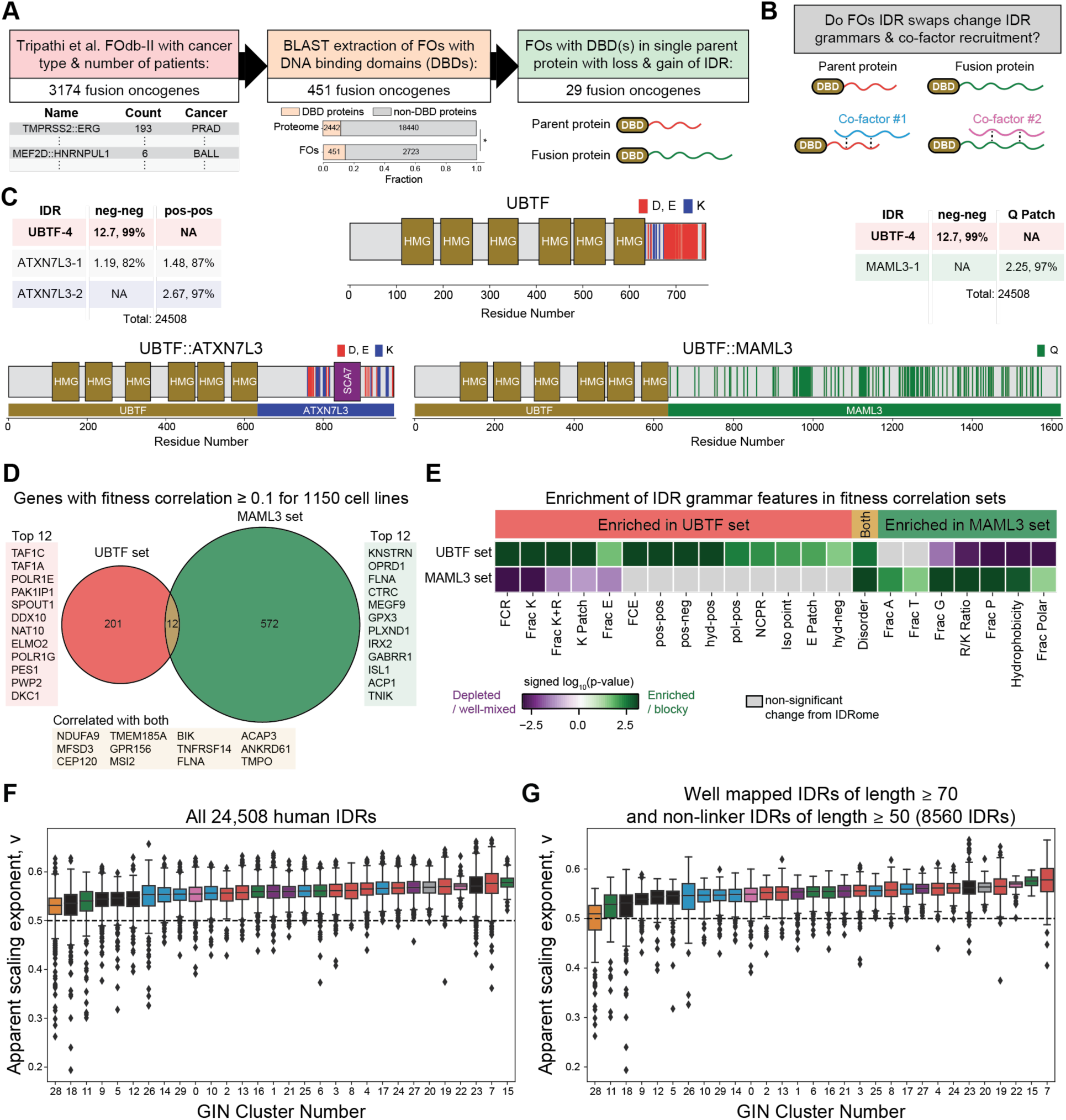
Cancer driver genes are enriched in distinct GIN clusters, related to Figure 7. A. Flowchart for extracting fusions containing DNA binding domains from the patient derived fusion dataset of Tripathi et al.,^164^. DBDs are enriched in fusions when compared to the proteome (Fisher’s exact test, 3.87×10^-5^). B. Schematic depicting the hypothesis that fusions may change IDR grammars and rewire interactions with co-factors. C. Schematics of WT UBTF and UBTF fusions associated with cancer. The exceptional grammar changes in the C-terminal tail for the two fusions are quantified in the left and right tables. D. Venn diagram of genes with positively correlated fitness effects ≥ 0.1 for UBTF and MAML3. E. NARDINI+ analysis of IDRs within the positively correlated fitness effect gene sets for UBTF and MAML3. P-values were calculated using the two-sample Kolmogorov-Smirnov test ^214^ comparing the z-score distribution for IDRs in the given fitness correlated set versus the z-score distribution of the remaining IDRome. F. Box and whisker plots of the ALBATROSS predicted apparent scaling exponent for all human IDRs. Outliers are shown as diamonds. G. Same as E, but for well mapped IDRs of length ≥ 70 and non-linker IDRs of length ≥ 50.H.

## References

1. Chothia, C., Gough, J., Vogel, C., and Teichmann, S.A. (2003). Evolution of the Protein Repertoire. Science 300, 1701–1703. 10.1126/science.1085371.

2. Jumper, J., Evans, R., Pritzel, A., Green, T., Figurnov, M., Ronneberger, O., Tunyasuvunakool, K., Bates, R., Zidek, A., Potapenko, A., et al. (2021). Highly accurate protein structure prediction with AlphaFold. Nature 596, 583–589. 10.1038/s41586-021-03819-2.

3. Holehouse, A.S., and Kragelund, B.B. (2024). The molecular basis for cellular function of intrinsically disordered protein regions. Nature Reviews Molecular Cell Biology 25, 187–211. 10.1038/s41580-023-00673-0.

4. Babu, M.M., Kriwacki, R.W., and Pappu, R.V. (2012). Versatility from Protein Disorder. Science 337, 1460–1461. 10.1126/science.1228775.

5. Wright, P.E., and Dyson, H.J. (1999). Intrinsically unstructured proteins: re-assessing the protein structure-function paradigm. Journal of Molecular Biology 293, 321–331. 10.1006/jmbi.1999.3110.

6. Mittal, A., Holehouse, A.S., Cohan, M.C., and Pappu, R.V. (2018). Sequence-to-Conformation Relationships of Disordered Regions Tethered to Folded Domains of Proteins. Journal of Molecular Biology 430, 2403–2421. 10.1016/j.jmb.2018.05.012.

7. Uversky, V.N., Gillespie, J.R., and Fink, A.L. (2000). Why are “natively unfolded” proteins unstructured under physiologic conditions? Proteins: Structure, Function, and Bioinformatics 41, 415–427. 10.1002/1097-0134(20001115)41:3<415::AID-PROT130>3.0.CO;2-7.

8. van der Lee, R., Buljan, M., Lang, B., Weatheritt, R.J., Daughdrill, G.W., Dunker, A.K., Fuxreiter, M., Gough, J., Gsponer, J., Jones, D.T., et al. (2014). Classification of intrinsically disordered regions and proteins. Chemical Reviews 114, 6589–6631. 10.1021/cr400525m.

9. Dyson, H.J., and Wright, P.E. (2005). Intrinsically unstructured proteins and their functions. Nature Reviews Molecular Cell Biology 6, 197–208. 10.1038/nrm1589.

10. Lotthammer, J.M., Ginell, G.M., Griffith, D., Emenecker, R.J., and Holehouse, A.S. (2024). Direct prediction of intrinsically disordered protein conformational properties from sequence. Nature Methods 21, 465–476. 10.1038/s41592-023-02159-5.

11. Tesei, G., Trolle, A.I., Jonsson, N., Betz, J., Knudsen, F.E., Pesce, F., Johansson, K.E., and Lindorff-Larsen, K. (2024). Conformational ensembles of the human intrinsically disordered proteome. Nature 626, 897–904. 10.1038/s41586-023-07004-5.

12. Dunker, A.K., Lawson, J.D., Brown, C.J., Williams, R.M., Romero, P., Oh, J.S., Oldfield, C.J., Campen, A.M., Ratliff, C.M., Hipps, K.W., et al. (2001). Intrinsically disordered protein. Journal of Molecular Graphics and Modelling 19, 26–59. 10.1016/S1093-3263(00)00138-8.

13. Oldfield, C.J., and Dunker, A.K. (2014). Intrinsically Disordered Proteins and Intrinsically Disordered Protein Regions. Annual Review of Biochemistry 83, 553–584. 10.1146/annurev-biochem-072711-164947.

14. Parchure, A., Tian, M., Stalder, D., Boyer, C.K., Bearrows, S.C., Rohli, K.E., Zhang, J., Rivera-Molina, F., Ramazanov, B.R., Mahata, S.K., et al. (2022). Liquid–liquid phase separation facilitates the biogenesis of secretory storage granules. Journal of Cell Biology 221. 10.1083/jcb.202206132.

15. Lockless, S.W., and Ranganathan, R. (1999). Evolutionarily Conserved Pathways of Energetic Connectivity in Protein Families. Science 286, 295–299. doi:10.1126/science.286.5438.295.

16. Halabi, N., Rivoire, O., Leibler, S., and Ranganathan, R. (2009). Protein Sectors: Evolutionary Units of Three-Dimensional Structure. Cell 138, 774–786. 10.1016/j.cell.2009.07.038.

17. Lemke, E.A., Babu, M.M., Kriwacki, R.W., Mittag, T., Pappu, R.V., Wright, P.E., and Forman-Kay, J.D. (2024). Intrinsic disorder: A term to define the specific physicochemical characteristic of protein conformational heterogeneity. Molecular Cell 84, 1188–1190. 10.1016/j.molcel.2024.02.024.

18. Zarin, T., Strome, B., Peng, G., Pritisanac, I., Forman-Kay, J.D., and Moses, A.M. (2021). Identifying molecular features that are associated with biological function of intrinsically disordered protein regions. Elife 10. 10.7554/eLife.60220.

19. Bertagna, A., Toptygin, D., Brand, L., and Barrick, D. (2008). The effects of conformational heterogeneity on the binding of the Notch intracellular domain to effector proteins: a case of biologically tuned disorder. Biochemical Society Transactions 36, 157–166. 10.1042/bst0360157.

20. Cohan, M.C., Shinn, M.K., Lalmansingh, J.M., and Pappu, R.V. (2022). Uncovering Non-random Binary Patterns Within Sequences of Intrinsically Disordered Proteins. Journal of Molecular Biology 434, 167373. 10.1016/j.jmb.2021.167373.

21. Shinn, M.K., Cohan, M.C., Bullock, J.L., Ruff, K.M., Levin, P.A., and Pappu, R.V. (2022). Connecting sequence features within the disordered C-terminal linker of Bacillus subtilis FtsZ to functions and bacterial cell division. Proceedings of the National Academy of Sciences 119, e2211178119. doi:10.1073/pnas.2211178119.

22. Cohan, M.C., Ruff, K.M., and Pappu, R.V. (2019). Information theoretic measures for quantifying sequence–ensemble relationships of intrinsically disordered proteins. Protein Engineering, Design and Selection 32, 191–202. 10.1093/protein/gzz014.

23. Riley, A.C., Ashlock, D.A., and Graether, S.P. (2023). The difficulty of aligning intrinsically disordered protein sequences as assessed by conservation and phylogeny. PLOS ONE 18, e0288388. 10.1371/journal.pone.0288388.

24. Wang, J., Choi, J.M., Holehouse, A.S., Lee, H.O., Zhang, X., Jahnel, M., Maharana, S., Lemaitre, R., Pozniakovsky, A., Drechsel, D., et al. (2018). A Molecular Grammar Governing the Driving Forces for Phase Separation of Prion-like RNA Binding Proteins. Cell 174, 688–699 e616. 10.1016/j.cell.2018.06.006.

25. Martin, E.W., Holehouse, A.S., Peran, I., Farag, M., Incicco, J.J., Bremer, A., Grace, C.R., Soranno, A., Pappu, R.V., and Mittag, T. (2020). Valence and patterning of aromatic residues determine the phase behavior of prion-like domains. Science 367, 694–699. 10.1126/science.aaw8653.

26. Martin, E.W., Holehouse, A.S., Grace, C.R., Hughes, A., Pappu, R.V., and Mittag, T. (2016). Sequence Determinants of the Conformational Properties of an Intrinsically Disordered Protein Prior to and upon Multisite Phosphorylation. Journal of the American Chemical Society 138, 15323–15335. 10.1021/jacs.6b10272.

27. Ruff, K.M., Choi, Y.H., Cox, D., Ormsby, A.R., Myung, Y., Ascher, D.B., Radford, S.E., Pappu, R.V., and Hatters, D.M. (2022). Sequence grammar underlying the unfolding and phase separation of globular proteins. Molecular Cell 82, 3193–3208 e3198. 10.1016/j.molcel.2022.06.024.

28. Bremer, A., Farag, M., Borcherds, W.M., Peran, I., Martin, E.W., Pappu, R.V., and Mittag, T. (2022). Deciphering how naturally occurring sequence features impact the phase behaviours of disordered prion-like domains. Nature Chemistry 14, 196–207. 10.1038/s41557-021-00840-w.

29. Das, R.K., and Pappu, R.V. (2013). Conformations of intrinsically disordered proteins are influenced by linear sequence distributions of oppositely charged residues. Proceedings of the National Academy of Sciences 110, 13392–13397. 10.1073/pnas.1304749110.

30. Zheng, W., Dignon, G., Brown, M., Kim, Y.C., and Mittal, J. (2020). Hydropathy Patterning Complements Charge Patterning to Describe Conformational Preferences of Disordered Proteins. The Journal of Physical Chemistry Letters 11, 3408–3415. 10.1021/acs.jpclett.0c00288.

31. Sawle, L., and Ghosh, K. (2015). A theoretical method to compute sequence dependent configurational properties in charged polymers and proteins. The Journal of Chemical Physics 143. 10.1063/1.4929391.

32. Pak, C.W., Kosno, M., Holehouse, A.S., Padrick, S.B., Mittal, A., Ali, R., Yunus, A.A., Liu, D.R., Pappu, R.V., and Rosen, M.K. (2016). Sequence Determinants of Intracellular Phase Separation by Complex Coacervation of a Disordered Protein. Molecular Cell 63, 72–85. 10.1016/j.molcel.2016.05.042.

33. Morffy, N., Van den Broeck, L., Miller, C., Emenecker, R.J., Bryant, J.A., Lee, T.M., Sageman-Furnas, K., Wilkinson, E.G., Pathak, S., Kotha, S.R., et al. (2024). Identification of plant transcriptional activation domains. Nature 632, 166–173. 10.1038/s41586-024-07707-3.

34. Holehouse, A.S., Das, R.K., Ahad, J.N., Richardson, M.O., and Pappu, R.V. (2017). CIDER: Resources to Analyze Sequence-Ensemble Relationships of Intrinsically Disordered Proteins. Biophysical Journal 112, 16–21. 10.1016/j.bpj.2016.11.3200.

35. Lin, Y., Currie, S.L., and Rosen, M.K. (2017). Intrinsically disordered sequences enable modulation of protein phase separation through distributed tyrosine motifs. Journal of Biological Chemistry 292, 19110–19120. 10.1074/jbc.M117.800466.

36. Sabari, B.R., Hyman, A.A., and Hnisz, D. (2024). Functional specificity in biomolecular condensates revealed by genetic complementation. Nature Reviews Genetics. 10.1038/s41576-024-00780-4.

37. Staller, M.V., Ramirez, E., Kotha, S.R., Holehouse, A.S., Pappu, R.V., and Cohen, B.A. (2022). Directed mutational scanning reveals a balance between acidic and hydrophobic residues in strong human activation domains. Cell Systems 13, 334–345.e335. 10.1016/j.cels.2022.01.002.

38. Bigman, L.S., Iwahara, J., and Levy, Y. (2022). Negatively Charged Disordered Regions are Prevalent and Functionally Important Across Proteomes. Journal of Molecular Biology 434, 167660. 10.1016/j.jmb.2022.167660.

39. Vuzman, D., and Levy, Y. (2010). DNA search efficiency is modulated by charge composition and distribution in the intrinsically disordered tail. Proceedings of the National Academy of Sciences 107, 21004–21009. doi:10.1073/pnas.1011775107.

40. Marsh, J.A., and Forman-Kay, J.D. (2010). Sequence determinants of compaction in intrinsically disordered proteins. Biophysical Journal 98, 2383–2390. 10.1016/j.bpj.2010.02.006.

41. Quiroz, F.G., and Chilkoti, A. (2015). Sequence heuristics to encode phase behaviour in intrinsically disordered protein polymers. Nature Materials 14, 1164–1171. 10.1038/nmat4418.

42. Dzuricky, M., Rogers, B.A., Shahid, A., Cremer, P.S., and Chilkoti, A. (2020). De novo engineering of intracellular condensates using artificial disordered proteins. Nature Chemistry 12, 814–825. 10.1038/s41557-020-0511-7.

43. Banjade, S., Wu, Q., Mittal, A., Peeples, W.B., Pappu, R.V., and Rosen, M.K. (2015). Conserved interdomain linker promotes phase separation of the multivalent adaptor protein Nck. Proceedings of the National Academy of Sciences 112, E6426–E6435. doi:10.1073/pnas.1508778112.

44. Patil, A., Strom, A.R., Paulo, J.A., Collings, C.K., Ruff, K.M., Shinn, M.K., Sankar, A., Cervantes, K.S., Wauer, T., St Laurent, J.D., et al. (2023). A disordered region controls cBAF activity via condensation and partner recruitment. Cell 186, 4936–4955 e4926. 10.1016/j.cell.2023.08.032.

45. Greig, J.A., Nguyen, T.A., Lee, M., Holehouse, A.S., Posey, A.E., Pappu, R.V., and Jedd, G. (2020). Arginine-Enriched Mixed-Charge Domains Provide Cohesion for Nuclear Speckle Condensation. Molecular Cell 77, 1237–1250 e1234. 10.1016/j.molcel.2020.01.025.

46. Hoffmann, C., Ruff, K.M., Edu, I.A., Shinn, M.K., Tromm, J.V., King, M.R., Pant, A., Ausserwöger, H., Morgan, J.R., Knowles, T.P.J., et al. (2025). Synapsin condensation is governed by sequence-encoded molecular grammars. Journal of Molecular Biology *In press*. 10.1101/2024.08.03.606464.

47. Hadarovich, A., Singh, H.R., Ghosh, S., Scheremetjew, M., Rostam, N., Hyman, A.A., and Toth-Petroczy, A. (2024). PICNIC accurately predicts condensate-forming proteins regardless of their structural disorder across organisms. Nature Communications 15, 10668. 10.1038/s41467-024-55089-x.

48. King, M.R., Ruff, K.M., Lin, A.Z., Pant, A., Farag, M., Lalmansingh, J.M., Wu, T., Fossat, M.J., Ouyang, W., Lew, M.D., et al. (2024). Macromolecular condensation organizes nucleolar sub-phases to set up a pH gradient. Cell 187, 1889–1906 e1824. 10.1016/j.cell.2024.02.029.

49. Lyons, H., Veettil, R.T., Pradhan, P., Fornero, C., De La Cruz, N., Ito, K., Eppert, M., Roeder, R.G., and Sabari, B.R. (2023). Functional partitioning of transcriptional regulators by patterned charge blocks. Cell 186, 327–345 e328. 10.1016/j.cell.2022.12.013.

50. Basu, S., Martinez-Cristobal, P., Frigole-Vivas, M., Pesarrodona, M., Lewis, M., Szulc, E., Banuelos, C.A., Sanchez-Zarzalejo, C., Bielskute, S., Zhu, J., et al. (2023). Rational optimization of a transcription factor activation domain inhibitor. Nature Structural & Molecular Biology 30, 1958–1969. 10.1038/s41594-023-01159-5.

51. Jonas, F., Carmi, M., Krupkin, B., Steinberger, J., Brodsky, S., Jana, T., and Barkai, N. (2023). The molecular grammar of protein disorder guiding genome-binding locations. Nucleic Acids Research 51, 4831–4844. 10.1093/nar/gkad184.

52. Ravarani, C.N., Erkina, T.Y., De Baets, G., Dudman, D.C., Erkine, A.M., and Babu, M.M. (2018). High-throughput discovery of functional disordered regions: investigation of transactivation domains. Molecular Systems Biology 14, e8190. 10.15252/msb.20188190.

53. Staller, M.V., Holehouse, A.S., Swain-Lenz, D., Das, R.K., Pappu, R.V., and Cohen, B.A. (2018). A High-Throughput Mutational Scan of an Intrinsically Disordered Acidic Transcriptional Activation Domain. Cell Systems 6, 444–455.e446. 10.1016/j.cels.2018.01.015.

54. Davey, N.E., Van Roey, K., Weatheritt, R.J., Toedt, G., Uyar, B., Altenberg, B., Budd, A., Diella, F., Dinkel, H., and Gibson, T.J. (2012). Attributes of short linear motifs. Molecular BioSystems 8, 268–281. 10.1039/C1MB05231D.

55. Nguyen Ba, A.N., Yeh, B.J., van Dyk, D., Davidson, A.R., Andrews, B.J., Weiss, E.L., and Moses, A.M. (2012). Proteome-Wide Discovery of Evolutionary Conserved Sequences in Disordered Regions. Science Signaling 5, rs1–rs1. doi:10.1126/scisignal.2002515.

56. Horton, C.A., Alexandari, A.M., Hayes, M.G.B., Marklund, E., Schaepe, J.M., Aditham, A.K., Shah, N., Suzuki, P.H., Shrikumar, A., Afek, A., et al. (2023). Short tandem repeats bind transcription factors to tune eukaryotic gene expression. Science 381, eadd1250. doi:10.1126/science.add1250.

57. Kim, J.-Y., Meng, F., Yoo, J., and Chung, H.S. (2018). Diffusion-limited association of disordered protein by non-native electrostatic interactions. Nature Communications 9, 4707. 10.1038/s41467-018-06866-y.

58. Karlsson, E., Schnatwinkel, J., Paissoni, C., Andersson, E., Herrmann, C., Camilloni, C., and Jemth, P. (2022). Disordered Regions Flanking the Binding Interface Modulate Affinity between CBP and NCOA. Journal of Molecular Biology 434, 167643. 10.1016/j.jmb.2022.167643.

59. Sherry, K.P., Das, R.K., Pappu, R.V., and Barrick, D. (2017). Control of transcriptional activity by design of charge patterning in the intrinsically disordered RAM region of the Notch receptor. Proceedings of the National Academy of Sciences 114, E9243–E9252. 10.1073/pnas.1706083114.

60. Das, R.K., Huang, Y., Phillips, A.H., Kriwacki, R.W., and Pappu, R.V. (2016). Cryptic sequence features within the disordered protein p27^Kip1^ regulate cell cycle signaling. Proceedings of the National Academy of Sciences 113, 5616–5621. 10.1073/pnas.1516277113.

61. Halpin, J.C., and Keating, A.E. (2025). PairK: Pairwise k-mer alignment for quantifying protein motif conservation in disordered regions. Protein Science 34, e70004. 10.1002/pro.70004.

62. Van Roey, K., Uyar, B., Weatheritt, R.J., Dinkel, H., Seiler, M., Budd, A., Gibson, T.J., and Davey, N.E. (2014). Short Linear Motifs: Ubiquitous and Functionally Diverse Protein Interaction Modules Directing Cell Regulation. Chemical Reviews 114, 6733–6778. 10.1021/cr400585q.

63. Das, R.K., Mao, A.H., and Pappu, R.V. (2012). Unmasking Functional Motifs Within Disordered Regions of Proteins. Science Signaling 5, pe17–pe17. doi:10.1126/scisignal.2003091.

64. Ferrie, J.J., Karr, J.P., Tjian, R., and Darzacq, X. (2022). “Structure”-function relationships in eukaryotic transcription factors: The role of intrinsically disordered regions in gene regulation. Molecular Cell 82, 3970–3984. 10.1016/j.molcel.2022.09.021.

65. Pritišanac, I., Alderson, T.R., Kolarić, Đ., Zarin, T., Xie, S., Lu, A., Alam, A., Maqsood, A., Youn, J.-Y., Forman-Kay, J.D., and Moses, A.M. (2024). A Functional Map of the Human Intrinsically Disordered Proteome. bioRxiv, 2024.2003.2015.585291. 10.1101/2024.03.15.585291.

66. Kilgore, H.R., Chinn, I., Mikhael, P.G., Mitnikov, I., Van Dongen, C., Zylberberg, G., Afeyan, L., Banani, S., Wilson-Hawken, S., Lee, T.I., et al. (2025). Protein codes promote selective subcellular compartmentalization. Science 0, eadq2634. 10.1126/science.adq2634.

67. Zarin, T., Strome, B., Nguyen Ba, A.N., Alberti, S., Forman-Kay, J.D., and Moses, A.M. (2019). Proteome-wide signatures of function in highly diverged intrinsically disordered regions. Elife 8. 10.7554/eLife.46883.

68. King, M.R., Ruff, K.M., and Pappu, R.V. (2024). Emergent microenvironments of nucleoli. Nucleus 15, 2319957. 10.1080/19491034.2024.2319957.

69. Kyte, J., and Doolittle, R.F. (1982). A simple method for displaying the hydropathic character of a protein. Journal of Molecular Biology 157, 105–132. 10.1016/0022-2836(82)90515-0.

70. Forman-Kay, Julie D., and Mittag, T. (2013). From Sequence and Forces to Structure, Function, and Evolution of Intrinsically Disordered Proteins. Structure 21, 1492–1499. 10.1016/j.str.2013.08.001.

71. Holla, A., Martin, E.W., Dannenhoffer-Lafage, T., Ruff, K.M., König, S.L.B., Nüesch, M.F., Chowdhury, A., Louis, J.M., Soranno, A., Nettels, D., et al. (2024). Identifying Sequence Effects on Chain Dimensions of Disordered Proteins by Integrating Experiments and Simulations. JACS Au 4, 4729–4743. 10.1021/jacsau.4c00673.

72. Das, R.K., Ruff, K.M., and Pappu, R.V. (2015). Relating sequence encoded information to form and function of intrinsically disordered proteins. Current Opinion in Structural Biology 32, 102–112. 10.1016/j.sbi.2015.03.008.

73. González-Foutel, N.S., Glavina, J., Borcherds, W.M., Safranchik, M., Barrera-Vilarmau, S., Sagar, A., Estaña, A., Barozet, A., Garrone, N.A., Fernandez-Ballester, G., et al. (2022). Conformational buffering underlies functional selection in intrinsically disordered protein regions. Nature Structural & Molecular Biology 29, 781–790. 10.1038s41594-022-00811-w.

74. Moses, D., Ginell, G.M., Holehouse, A.S., and Sukenik, S. (2023). Intrinsically disordered regions are poised to act as sensors of cellular chemistry. Trends in Biochemical Sciences 48, 1019–1034. 10.1016/j.tibs.2023.08.001.

75. Marsh, J.A., Baker, J.M.R., Tollinger, M., and Forman-Kay, J.D. (2008). Calculation of Residual Dipolar Couplings from Disordered State Ensembles Using Local Alignment. Journal of the American Chemical Society 130, 7804–7805. 10.1021/ja802220c.

76. Nott, Timothy J., Petsalaki, E., Farber, P., Jervis, D., Fussner, E., Plochowietz, A., Craggs, T.D., Bazett-Jones, David P., Pawson, T., Forman-Kay, Julie D., and Baldwin, Andrew J. (2015). Phase Transition of a Disordered Nuage Protein Generates Environmentally Responsive Membraneless Organelles. Molecular Cell 57, 936–947. 10.1016/j.molcel.2015.01.013.

77. Crick, S.L., Jayaraman, M., Frieden, C., Wetzel, R., and Pappu, R.V. (2006). Fluorescence correlation spectroscopy shows that monomeric polyglutamine molecules form collapsed structures in aqueous solutions. Proceedings of the National Academy of Sciences 103, 16764–16769. doi:10.1073/pnas.0608175103.

78. Tran, H.T., Mao, A., and Pappu, R.V. (2008). Role of Backbone−Solvent Interactions in Determining Conformational Equilibria of Intrinsically Disordered Proteins. Journal of the American Chemical Society 130, 7380–7392. 10.1021/ja710446s.

79. Dignon, G.L., Zheng, W., Kim, Y.C., Best, R.B., and Mittal, J. (2018). Sequence determinants of protein phase behavior from a coarse-grained model. PLOS Computational Biology 14, e1005941. 10.1371/journal.pcbi.1005941.

80. Chow, C.F.W., Ghosh, S., Hadarovich, A., and Toth-Petroczy, A. (2024). SHARK enables sensitive detection of evolutionary homologs and functional analogs in unalignable and disordered sequences. Proceedings of the National Academy of Sciences 121, e2401622121. doi:10.1073/pnas.2401622121.

81. Mao, A.H., Crick, S.L., Vitalis, A., Chicoine, C.L., and Pappu, R.V. (2010). Net charge per residue modulates conformational ensembles of intrinsically disordered proteins. Proceedings of the National Academy of Sciences 107, 8183–8188. 10.1073/pnas.0911107107.

82. Mao, Albert H., Lyle, N., and Pappu, Rohit V. (2012). Describing sequence–ensemble relationships for intrinsically disordered proteins. Biochemical Journal 449, 307–318. 10.1042/bj20121346.

83. Müller-Späth, S., Soranno, A., Hirschfeld, V., Hofmann, H., Rüegger, S., Reymond, L., Nettels, D., and Schuler, B. (2010). Charge interactions can dominate the dimensions of intrinsically disordered proteins. Proceedings of the National Academy of Sciences 107, 14609–14614. 10.1073/pnas.1001743107.

84. Hofmann, H., Soranno, A., Borgia, A., Gast, K., Nettels, D., and Schuler, B. (2012). Polymer scaling laws of unfolded and intrinsically disordered proteins quantified with single-molecule spectroscopy. Proceedings of the National Academy of Sciences 109, 16155–16160. 10.1073/pnas.1207719109.

85. Thomasen, F.E., and Lindorff-Larsen, K. (2022). Conformational ensembles of intrinsically disordered proteins and flexible multidomain proteins. Biochemical Society Transactions 50, 541–554. 10.1042/bst20210499.

86. Lyle, N., Das, R.K., and Pappu, R.V. (2013). A quantitative measure for protein conformational heterogeneity. The Journal of Chemical Physics 139. 10.1063/1.4812791.

87. Lin, Y.-H., Song, J., Gomes, G.-N., Das, S., Gradinaru, C.C., Forman-Kay, J.D., and Sun Chan, H. (2018). Conformational Heterogeneity and Theory of Sequence-Specific Functional Phase Separation of Intrinsically Disordered Proteins. Biophysical Journal 114, 6a. 10.1016/j.bpj.2017.11.064.

88. Halfmann, R., Alberti, S., Krishnan, R., Lyle, N., O’Donnell, Charles W., King, Oliver D., Berger, B., Pappu, Rohit V., and Lindquist, S. (2011). Opposing Effects of Glutamine and Asparagine Govern Prion Formation by Intrinsically Disordered Proteins. Molecular Cell 43, 72–84. 10.1016/j.molcel.2011.05.013.

89. Hartigan, J.A., and Wong, M.A. (1979). Algorithm AS 136: A K-Means Clustering Algorithm. Journal of the Royal Statistical Society. Series C (Applied Statistics) 28, 100–108. 10.2307/2346830.

90. Gemayel, R., Chavali, S., Pougach, K., Legendre, M., Zhu, B., Boeynaems, S., van der Zande, E., Gevaert, K., Rousseau, F., Schymkowitz, J., et al. (2015). Variable Glutamine-Rich Repeats Modulate Transcription Factor Activity. Molecular Cell 59, 615–627. 10.1016/j.molcel.2015.07.003.

91. Rauscher, S., Baud, S., Miao, M., Keeley, Fred W., and Pomès, R. (2006). Proline and Glycine Control Protein Self-Organization into Elastomeric or Amyloid Fibrils. Structure 14, 1667–1676. 10.1016/j.str.2006.09.008.

92. UniProt, C. (2023). UniProt: the Universal Protein Knowledgebase in 2023. Nucleic Acids Res 51, D523–D531. 10.1093/nar/gkac1052.

93. Lipman, D.J., and Pearson, W.R. (1985). Rapid and Sensitive Protein Similarity Searches. Science 227, 1435–1441. doi:10.1126/science.2983426.

94. Diekmann, Y., and Pereira-Leal, José B. (2012). Evolution of intracellular compartmentalization. Biochemical Journal 449, 319–331. 10.1042/bj20120957.

95. Banani, S.F., Lee, H.O., Hyman, A.A., and Rosen, M.K. (2017). Biomolecular condensates: organizers of cellular biochemistry. Nature Reviews Molecular Cell Biology 18, 285–298. 10.1038/nrm.2017.7.

96. Thul, P.J., Akesson, L., Wiking, M., Mahdessian, D., Geladaki, A., Ait Blal, H., Alm, T., Asplund, A., Bjork, L., Breckels, L.M., et al. (2017). A subcellular map of the human proteome. Science 356. 10.1126/science.aal3321.

97. Scott, M.S., Boisvert, F.M., McDowall, M.D., Lamond, A.I., and Barton, G.J. (2010). Characterization and prediction of protein nucleolar localization sequences. Nucleic Acids Research 38, 7388–7399. 10.1093/nar/gkq653.

98. Handwerger, K.E., Cordero, J.A., and Gall, J.G. (2005). Cajal Bodies, Nucleoli, and Speckles in the Xenopus Oocyte Nucleus Have a Low-Density, Sponge-like Structure. Molecular Biology of the Cell 16, 202–211. 10.1091/mbc.e04-08-0742.

99. Robert-Paganin, J., Réty, S., and Leulliot, N. (2015). Regulation of DEAH/RHA Helicases by G-Patch Proteins. BioMed Research International 2015, 931857. 10.1155/2015/931857.

100. Aravind, L., and Koonin, E.V. (1999). G-patch: a new conserved domain in eukaryotic RNA-processing proteins and type D retroviral polyproteins. Trends in Biochemical Sciences 24, 342–344. 10.1016/s0968-0004(99)01437-1.

101. Sillitoe, I., Bordin, N., Dawson, N., Waman, V.P., Ashford, P., Scholes, H.M., Pang, C.S.M., Woodridge, L., Rauer, C., Sen, N., et al. (2020). CATH: increased structural coverage of functional space. Nucleic Acids Research 49, D266–D273. 10.1093/nar/gkaa1079.

102. Andreeva, A., Kulesha, E., Gough, J., and Murzin, A.G. (2019). The SCOP database in 2020: expanded classification of representative family and superfamily domains of known protein structures. Nucleic Acids Research 48, D376–D382. 10.1093/nar/gkz1064.

103. Ashburner, M., Ball, C.A., Blake, J.A., Botstein, D., Butler, H., Cherry, J.M., Davis, A.P., Dolinski, K., Dwight, S.S., Eppig, J.T., et al. (2000). Gene ontology: tool for the unification of biology. The Gene Ontology Consortium. Nature Genetics 25, 25–29. 10.1038/75556.

104. Gene Ontology, C., Aleksander, S.A., Balhoff, J., Carbon, S., Cherry, J.M., Drabkin, H.J., Ebert, D., Feuermann, M., Gaudet, P., Harris, N.L., et al. (2023). The Gene Ontology knowledgebase in 2023. Genetics 224. 10.1093/genetics/iyad031.

105. Arafeh, R., Shibue, T., Dempster, J.M., Hahn, W.C., and Vazquez, F. (2025). The present and future of the Cancer Dependency Map. Nature Reviews Cancer 25, 59–73. 10.1038/s41568-024-00763-x.

106. Tsherniak, A., Vazquez, F., Montgomery, P.G., Weir, B.A., Kryukov, G., Cowley, G.S., Gill, S., Harrington, W.F., Pantel, S., Krill-Burger, J.M., et al. (2017). Defining a Cancer Dependency Map. Cell 170, 564–576 e516. 10.1016/j.cell.2017.06.010.

107. Pan, J., Meyers, R.M., Michel, B.C., Mashtalir, N., Sizemore, A.E., Wells, J.N., Cassel, S.H., Vazquez, F., Weir, B.A., Hahn, W.C., et al. (2018). Interrogation of Mammalian Protein Complex Structure, Function, and Membership Using Genome-Scale Fitness Screens. Cell Systems 6, 555–568.e557. 10.1016/j.cels.2018.04.011.

108. McDonald, E.R., III, de Weck, A., Schlabach, M.R., Billy, E., Mavrakis, K.J., Hoffman, G.R., Belur, D., Castelletti, D., Frias, E., Gampa, K., et al. (2017). Project DRIVE: A Compendium of Cancer Dependencies and Synthetic Lethal Relationships Uncovered by Large-Scale, Deep RNAi Screening. Cell 170, 577–592.e510. 10.1016/j.cell.2017.07.005.

109. Pederson, T. (1998). The plurifunctional nucleolus. Nucleic Acids Research 26, 3871–3876. 10.1093/nar/26.17.3871.

110. Boisvert, F.M., van Koningsbruggen, S., Navascues, J., and Lamond, A.I. (2007). The multifunctional nucleolus. Nat Rev Mol Cell Biol 8, 574–585. 10.1038/nrm2184.

111. Frottin, F., Schueder, F., Tiwary, S., Gupta, R., Korner, R., Schlichthaerle, T., Cox, J., Jungmann, R., Hartl, F.U., and Hipp, M.S. (2019). The nucleolus functions as a phase-separated protein quality control compartment. Science 365, 342–347. 10.1126/science.aaw9157.

112. Scott, D.D., and Oeffinger, M. (2016). Nucleolin and nucleophosmin: nucleolar proteins with multiple functions in DNA repair. Biochemistry and Cell Biology 94, 419–432. 10.1139/bcb-2016-0068.

113. Dorner, K., Ruggeri, C., Zemp, I., and Kutay, U. (2023). Ribosome biogenesis factors-from names to functions. EMBO J 42, e112699. 10.15252/embj.2022112699.

114. Otto, E., Culakova, E., Meng, S., Zhang, Z., Xu, H., Mohile, S., and Flannery, M.A. (2022). Overview of Sankey flow diagrams: Focusing on symptom trajectories in older adults with advanced cancer. Journal of Geriatric Oncology 13, 742–746. 10.1016/j.jgo.2021.12.017.

115. Seal, R.L., Braschi, B., Gray, K., Jones, T.E.M., Tweedie, S., Haim-Vilmovsky, L., and Bruford, E.A. (2023). Genenames.org: the HGNC resources in 2023. Nucleic Acids Res 51, D1003–D1009. 10.1093/nar/gkac888.

116. Seiler, M., Peng, S., Agrawal, A.A., Palacino, J., Teng, T., Zhu, P., Smith, P.G., Cancer Genome Atlas Research, N., Buonamici, S., and Yu, L. (2018). Somatic Mutational Landscape of Splicing Factor Genes and Their Functional Consequences across 33 Cancer Types. Cell Reports 23, 282–296 e284. 10.1016/j.celrep.2018.01.088.

117. Galganski, L., Urbanek, M.O., and Krzyzosiak, W.J. (2017). Nuclear speckles: molecular organization, biological function and role in disease. Nucleic Acids Research 45, 10350–10368. 10.1093/nar/gkx759.

118. Shevtsov, S.P., and Dundr, M. (2011). Nucleation of nuclear bodies by RNA. Nature Cell Biology 13, 167–173. 10.1038/ncb2157.

119. Ilik, I.A., Malszycki, M., Lubke, A.K., Schade, C., Meierhofer, D., and Aktas, T. (2020). SON and SRRM2 are essential for nuclear speckle formation. Elife 9. 10.7554/eLife.60579.

120. Wen, J., Lv, R., Ma, H., Shen, H., He, C., Wang, J., Jiao, F., Liu, H., Yang, P., Tan, L., et al. (2018). Zc3h13 Regulates Nuclear RNA m(6)A Methylation and Mouse Embryonic Stem Cell Self-Renewal. Molecular Cell 69, 1028–1038 e1026. 10.1016/j.molcel.2018.02.015.

121. Kanehisa, M., Furumichi, M., Sato, Y., Kawashima, M., and Ishiguro-Watanabe, M. (2023). KEGG for taxonomy-based analysis of pathways and genomes. Nucleic Acids Research 51, D587–D592. 10.1093/nar/gkac963.

122. Rodrigues, K.S., Petroski, L.P., Utumi, P.H., Ferrasa, A., and Herai, R.H. (2023). IARA: a complete and curated atlas of the biogenesis of spliceosome machinery during RNA splicing. Life Sci Alliance 6. 10.26508/lsa.202201593.

123. Li, D., Yu, W., and Lai, M. (2023). Towards understandings of serine/arginine-rich splicing factors. Acta Pharm Sin B 13, 3181–3207. 10.1016/j.apsb.2023.05.022.

124. Boija, A., Klein, I.A., Sabari, B.R., Dall’Agnese, A., Coffey, E.L., Zamudio, A.V., Li, C.H., Shrinivas, K., Manteiga, J.C., Hannett, N.M., et al. (2018). Transcription Factors Activate Genes through the Phase-Separation Capacity of Their Activation Domains. Cell 175, 1842–1855 e1816. 10.1016/j.cell.2018.10.042.

125. Otto, J.E., Ursu, O., Wu, A.P., Winter, E.B., Cuoco, M.S., Ma, S., Qian, K., Michel, B.C., Buenrostro, J.D., Berger, B., et al. (2023). Structural and functional properties of mSWI/SNF chromatin remodeling complexes revealed through single-cell perturbation screens. Molecular Cell 83, 1350–1367 e1357. 10.1016/j.molcel.2023.03.013.

126. Szentirmay, M.N., and Sawadogo, M. (2000). Spatial organization of RNA polymerase II transcription in the nucleus. Nucleic Acids Res 28, 2019–2025. 10.1093/nar/28.10.2019.

127. Cenik, B.K., and Shilatifard, A. (2021). COMPASS and SWI/SNF complexes in development and disease. Nature Reviews Genetics 22, 38–58. 10.1038/s41576-020-0278-0.

128. Girbig, M., Misiaszek, A.D., and Muller, C.W. (2022). Structural insights into nuclear transcription by eukaryotic DNA-dependent RNA polymerases. Nature Reviews Molecular Cell Biology 23, 603–622. 10.1038/s41580-022-00476-9.

129. Helmlinger, D., and Tora, L. (2017). Sharing the SAGA. Trends in Biochemical Sciences 42, 850–861. 10.1016/j.tibs.2017.09.001.

130. Richter, W.F., Nayak, S., Iwasa, J., and Taatjes, D.J. (2022). The Mediator complex as a master regulator of transcription by RNA polymerase II. Nature Reviews Cell Biology 23, 732–749. 10.1038/s41580-022-00498-3.

131. Sun, Z., and Xu, Y. (2020). Nuclear Receptor Coactivators (NCOAs) and Corepressors (NCORs) in the Brain. Endocrinology 161. 10.1210/endocr/bqaa083.

132. Turowski, T.W., and Boguta, M. (2021). Specific Features of RNA Polymerases I and III: Structure and Assembly. Front Mol Biosci 8, 680090. 10.3389/fmolb.2021.680090.

133. Valencia, A.M., Sankar, A., van der Sluijs, P.J., Satterstrom, F.K., Fu, J., Talkowski, M.E., Vergano, S.A.S., Santen, G.W.E., and Kadoch, C. (2023). Landscape of mSWI/SNF chromatin remodeling complex perturbations in neurodevelopmental disorders. Nature Genetics 55, 1400–1412. 10.1038/s41588-023-01451-6.

134. Shrinivas, K., Sabari, B.R., Coffey, E.L., Klein, I.A., Boija, A., Zamudio, A.V., Schuijers, J., Hannett, N.M., Sharp, P.A., Young, R.A., and Chakraborty, A.K. (2019). Enhancer Features that Drive Formation of Transcriptional Condensates. Molecular Cell 75, 549–561 e547. 10.1016/j.molcel.2019.07.009.

135. Battistello, E., Hixon, K.A., Comstock, D.E., Collings, C.K., Chen, X., Rodriguez Hernaez, J., Lee, S., Cervantes, K.S., Hinkley, M.M., Ntatsoulis, K., et al. (2023). Stepwise activities of mSWI/SNF family chromatin remodeling complexes direct T cell activation and exhaustion. Mol Cell 83, 1216–1236 e1212. 10.1016/j.molcel.2023.02.026.

136. Ma, L., Gao, Z., Wu, J., Zhong, B., Xie, Y., Huang, W., and Lin, Y. (2021). Co-condensation between transcription factor and coactivator p300 modulates transcriptional bursting kinetics. Molecular Cell 81, 1682–1697 e1687. 10.1016/j.molcel.2021.01.031.

137. Sabari, B.R., Dall’Agnese, A., Boija, A., Klein, I.A., Coffey, E.L., Shrinivas, K., Abraham, B.J., Hannett, N.M., Zamudio, A.V., Manteiga, J.C., et al. (2018). Coactivator condensation at super-enhancers links phase separation and gene control. Science 361. 10.1126/science.aar3958.

138. Wei, M.T., Chang, Y.C., Shimobayashi, S.F., Shin, Y., Strom, A.R., and Brangwynne, C.P. (2020). Nucleated transcriptional condensates amplify gene expression. Nat Cell Biol 22, 1187–1196. 10.1038/s41556-020-00578-6.

139. Guo, Y.E., Manteiga, J.C., Henninger, J.E., Sabari, B.R., Dall’Agnese, A., Hannett, N.M., Spille, J.H., Afeyan, L.K., Zamudio, A.V., Shrinivas, K., et al. (2019). Pol II phosphorylation regulates a switch between transcriptional and splicing condensates. Nature 572, 543–548. 10.1038/s41586-019-1464-0.

140. Jonas, F., Carmi, M., Krupkin, B., Steinberger, J., Brodsky, S., Jana, T., and Barkai, N. (2023). The molecular grammar of protein disorder guiding genome-binding locations. Nucleic Acids Res 51, 4831–4844. 10.1093/nar/gkad184.

141. Kumar, D.K., Jonas, F., Jana, T., Brodsky, S., Carmi, M., and Barkai, N. (2023). Complementary strategies for directing in vivo transcription factor binding through DNA binding domains and intrinsically disordered regions. Molecular Cell 83, 1462–1473 e1465. 10.1016/j.molcel.2023.04.002.

142. Lambert, S.A., Jolma, A., Campitelli, L.F., Das, P.K., Yin, Y., Albu, M., Chen, X., Taipale, J., Hughes, T.R., and Weirauch, M.T. (2018). The Human Transcription Factors. Cell 175, 598–599. 10.1016/j.cell.2018.09.045.

143. Schmeier, S., Alam, T., Essack, M., and Bajic, V.B. (2017). TcoF-DB v2: update of the database of human and mouse transcription co-factors and transcription factor interactions. Nucleic Acids Res 45, D145–D150. 10.1093/nar/gkw1007.

144. Panne, D., Maniatis, T., and Harrison, S.C. (2007). An atomic model of the interferon-beta enhanceosome. Cell 129, 1111–1123. 10.1016/j.cell.2007.05.019.

145. Davis, R.B., Supakar, A., Ranganath, A.K., Moosa, M.M., and Banerjee, P.R. (2024). Heterotypic interactions can drive selective co-condensation of prion-like low-complexity domains of FET proteins and mammalian SWI/SNF complex. Nat Commun 15, 1168. 10.1038/s41467-024-44945-5.

146. Zeigler, T.M., Chung, M.C., Narayan, O.P., and Guan, J. (2021). Protein phase separation: physical models and phase-separation-mediated cancer signaling. Advances in Physics: X 6, 1936638. 10.1080/23746149.2021.1936638.

147. Polyansky, A.A., Gallego, L.D., Efremov, R.G., Kohler, A., and Zagrovic, B. (2023). Protein compactness and interaction valency define the architecture of a biomolecular condensate across scales. Elife 12. 10.7554/eLife.80038.

148. Gestwicki, J.E., Cairo, C.W., Strong, L.E., Oetjen, K.A., and Kiessling, L.L. (2002). Influencing Receptor−Ligand Binding Mechanisms with Multivalent Ligand Architecture. Journal of the American Chemical Society 124, 14922–14933. 10.1021/ja027184x.

149. Grob, A., Colleran, C., and McStay, B. (2014). Construction of synthetic nucleoli in human cells reveals how a major functional nuclear domain is formed and propagated through cell division. Genes Dev 28, 220–230. 10.1101/gad.234591.113.

150. Ferrolino, M.C., Mitrea, D.M., Michael, J.R., and Kriwacki, R.W. (2018). Compositional adaptability in NPM1-SURF6 scaffolding networks enabled by dynamic switching of phase separation mechanisms. Nat Commun 9, 5064. 10.1038/s41467-018-07530-1.

151. Mitrea, D.M., Cika, J.A., Guy, C.S., Ban, D., Banerjee, P.R., Stanley, C.B., Nourse, A., Deniz, A.A., and Kriwacki, R.W. (2016). Nucleophosmin integrates within the nucleolus via multi-modal interactions with proteins displaying R-rich linear motifs and rRNA. Elife 5. 10.7554/eLife.13571.

152. Mitrea, D.M., Cika, J.A., Stanley, C.B., Nourse, A., Onuchic, P.L., Banerjee, P.R., Phillips, A.H., Park, C.G., Deniz, A.A., and Kriwacki, R.W. (2018). Self-interaction of NPM1 modulates multiple mechanisms of liquid-liquid phase separation. Nat Commun 9, 842. 10.1038/s41467-018-03255-3.

153. Feric, M., Vaidya, N., Harmon, T.S., Mitrea, D.M., Zhu, L., Richardson, T.M., Kriwacki, R.W., Pappu, R.V., and Brangwynne, C.P. (2016). Coexisting Liquid Phases Underlie Nucleolar Subcompartments. Cell 165, 1686–1697. 10.1016/j.cell.2016.04.047.

154. Xu, S., Lai, S.K., Sim, D.Y., Ang, W.S.L., Li, H.Y., and Roca, X. (2022). SRRM2 organizes splicing condensates to regulate alternative splicing. Nucleic Acids Res 50, 8599–8614. 10.1093/nar/gkac669.

155. Park, Y.K., Lee, J.E., Yan, Z., McKernan, K., O’Haren, T., Wang, W., Peng, W., and Ge, K. (2021). Interplay of BAF and MLL4 promotes cell type-specific enhancer activation. Nat Commun 12, 1630. 10.1038/s41467-021-21893-y.

156. Wang, L.H., Aberin, M.A.E., Wu, S., and Wang, S.P. (2021). The MLL3/4 H3K4 methyltransferase complex in establishing an active enhancer landscape. Biochem Soc Trans 49, 1041–1054. 10.1042/BST20191164.

157. Mendiratta, G., Ke, E., Aziz, M., Liarakos, D., Tong, M., and Stites, E.C. (2021). Cancer gene mutation frequencies for the U.S. population. Nat Commun 12, 5961. 10.1038/s41467-021-26213-y.

158. Zhao, Z., Aoi, Y., Philips, C.N., Meghani, K.A., Gold, S.R., Yu, Y., John, L.S., Qian, J., Zeidner, J.M., Meeks, J.J., and Shilatifard, A. (2023). Somatic mutations of MLL4/COMPASS induce cytoplasmic localization providing molecular insight into cancer prognosis and treatment. Proc Natl Acad Sci U S A 120, e2310063120. 10.1073/pnas.2310063120.

159. Fasciani, A., D’Annunzio, S., Poli, V., Fagnocchi, L., Beyes, S., Michelatti, D., Corazza, F., Antonelli, L., Gregoretti, F., Oliva, G., et al. (2020). MLL4-associated condensates counterbalance Polycomb-mediated nuclear mechanical stress in Kabuki syndrome. Nat Genet 52, 1397–1411. 10.1038/s41588-020-00724-8.

160. Ling, Y.H., Ye, Z., Liang, C., Yu, C., Park, G., Corden, J.L., and Wu, C. (2024). Disordered C-terminal domain drives spatiotemporal confinement of RNAPII to enhance search for chromatin targets. Nat Cell Biol 26, 581–592. 10.1038/s41556-024-01382-2.

161. Martinez-Jimenez, F., Muinos, F., Sentis, I., Deu-Pons, J., Reyes-Salazar, I., Arnedo-Pac, C., Mularoni, L., Pich, O., Bonet, J., Kranas, H., et al. (2020). A compendium of mutational cancer driver genes. Nature Reviews Cancer 20, 555–572. 10.1038/s41568-020-0290-x.

162. Gao, Q., Liang, W.W., Foltz, S.M., Mutharasu, G., Jayasinghe, R.G., Cao, S., Liao, W.W., Reynolds, S.M., Wyczalkowski, M.A., Yao, L., et al. (2018). Driver Fusions and Their Implications in the Development and Treatment of Human Cancers. Cell Rep 23, 227–238 e223. 10.1016/j.celrep.2018.03.050.

163. Latysheva, N.S., and Babu, M.M. (2019). Molecular Signatures of Fusion Proteins in Cancer. ACS Pharmacol Transl Sci 2, 122–133. 10.1021/acsptsci.9b00019.

164. Tripathi, S., Shirnekhi, H.K., Gorman, S.D., Chandra, B., Baggett, D.W., Park, C.G., Somjee, R., Lang, B., Hosseini, S.M.H., Pioso, B.J., et al. (2023). Defining the condensate landscape of fusion oncoproteins. Nature Ciommunications 14, 6008. 10.1038/s41467-023-41655-2.

165. Davis, R.B., Supakar, A., Ranganath, A.K., Moosa, M.M., and Banerjee, P.R. (2024). Heterotypic interactions can drive selective co-condensation of prion-like low-complexity domains of FET proteins and mammalian SWI/SNF complex. Nature Communications 15, 1168. 10.1038/s41467-024-44945-5.

166. Boulay, G., Sandoval, G.J., Riggi, N., Iyer, S., Buisson, R., Naigles, B., Awad, M.E., Rengarajan, S., Volorio, A., McBride, M.J., et al. (2017). Cancer-Specific Retargeting of BAF Complexes by a Prion-like Domain. Cell 171, 163–178 e119. 10.1016/j.cell.2017.07.036.

167. Passet, M., Kim, R., Gachet, S., Sigaux, F., Chaumeil, J., Galland, A., Sexton, T., Quentin, S., Hernandez, L., Larcher, L., et al. (2022). Concurrent CDX2 cis-deregulation and UBTF::ATXN7L3 fusion define a novel high-risk subtype of B-cell ALL. Blood 139, 3505–3518. 10.1182/blood.2021014723.

168. Alzofon, N., Koc, K., Panwell, K., Pozdeyev, N., Marshall, C.B., Albuja-Cruz, M., Raeburn, C.D., Nathanson, K.L., Cohen, D.L., Wierman, M.E., et al. (2021). Mastermind Like Transcriptional Coactivator 3 (MAML3) Drives Neuroendocrine Tumor Progression. MOlecular Cancer Research 19, 1476–1485. 10.1158/1541-7786.MCR-20-0992.

169. Stott, K., Watson, M., Bostock, M.J., Mortensen, S.A., Travers, A., Grasser, K.D., and Thomas, J.O. (2014). Structural Insights into the Mechanism of Negative Regulation of Single-box High Mobility Group Proteins by the Acidic Tail Domain *. Journal of Biological Chemistry 289, 29817–29826. 10.1074/jbc.M114.591115.

170. Göbel, U., Sander, C., Schneider, R., and Valencia, A. (1994). Correlated mutations and residue contacts in proteins. Proteins: Structure, Function, and Bioinformatics 18, 309–317. 10.1002/prot.340180402.

171. Shindyalov, I.N., Kolchanov, N.A., and Sander, C. (1994). Can three-dimensional contacts in protein structures be predicted by analysis of correlated mutations? Protein Engineering, Design and Selection 7, 349–358. 10.1093/protein/7.3.349.

172. Morcos, F., Pagnani, A., Lunt, B., Bertolino, A., Marks, D.S., Sander, C., Zecchina, R., Onuchic, J.N., Hwa, T., and Weigt, M. (2011). Direct-coupling analysis of residue coevolution captures native contacts across many protein families. Proceedings of the National Academy of Sciences 108, E1293–E1301. doi:10.1073/pnas.1111471108.

173. Morcos, F., Jana, B., Hwa, T., and Onuchic, J.N. (2013). Coevolutionary signals across protein lineages help capture multiple protein conformations. Proceedings of the National Academy of Sciences 110, 20533–20538. doi:10.1073/pnas.1315625110.

174. Baxter-Koenigs, A.R., El Nesr, G., and Barrick, D. (2022). Singular value decomposition of protein sequences as a method to visualize sequence and residue space. Protein Science 31, e4422. 10.1002/pro.4422.

175. Soranno, A., Buchli, B., Nettels, D., Cheng, R.R., Müller-Späth, S., Pfeil, S.H., Hoffmann, A., Lipman, E.A., Makarov, D.E., and Schuler, B. (2012). Quantifying internal friction in unfolded and intrinsically disordered proteins with single-molecule spectroscopy. Proceedings of the National Academy of Sciences 109, 17800–17806. 10.1073/pnas.1117368109.

176. Brucale, M., Schuler, B., and Samorì, B. (2014). Single-Molecule Studies of Intrinsically Disordered Proteins. Chemical Reviews 114, 3281–3317. 10.1021/cr400297g.

177. Soranno, A., Koenig, I., Borgia, M.B., Hofmann, H., Zosel, F., Nettels, D., and Schuler, B. (2014). Single-molecule spectroscopy reveals polymer effects of disordered proteins in crowded environments. Proceedings of the National Academy of Sciences 111, 4874–4879. 10.1073/pnas.1322611111.

178. Borgia, A., Zheng, W., Buholzer, K., Borgia, M.B., Schüler, A., Hofmann, H., Soranno, A., Nettels, D., Gast, K., Grishaev, A., et al. (2016). Consistent View of Polypeptide Chain Expansion in Chemical Denaturants from Multiple Experimental Methods. Journal of the American Chemical Society 138, 11714–11726. 10.1021/jacs.6b05917.

179. König, I., Soranno, A., Nettels, D., and Schuler, B. (2021). Impact of In-Cell and In-Vitro Crowding on the Conformations and Dynamics of an Intrinsically Disordered Protein. Angewandte Chemie International Edition 60, 10724–10729. 10.1002/anie.202016804.

180. Chowdhury, A., Nettels, D., and Schuler, B. (2023). Interaction Dynamics of Intrinsically Disordered Proteins from Single-Molecule Spectroscopy. Annual Review of Biophysics 52, 433–462. 10.1146/annurev-biophys-101122-071930.

181. Jensen, M.R., Ruigrok, R.W.H., and Blackledge, M. (2013). Describing intrinsically disordered proteins at atomic resolution by NMR. Current Opinion in Structural Biology 23, 426–435. 10.1016/j.sbi.2013.02.007.

182. Salvi, N., Abyzov, A., and Blackledge, M. (2016). Multi-Timescale Dynamics in Intrinsically Disordered Proteins from NMR Relaxation and Molecular Simulation. The Journal of Physical Chemistry Letters 7, 2483–2489. 10.1021/acs.jpclett.6b00885.

183. Adamski, W., Salvi, N., Maurin, D., Magnat, J., Milles, S., Jensen, M.R., Abyzov, A., Moreau, C.J., and Blackledge, M. (2019). A Unified Description of Intrinsically Disordered Protein Dynamics under Physiological Conditions Using NMR Spectroscopy. Journal of the American Chemical Society 141, 17817–17829. 10.1021/jacs.9b09002.

184. Lazar, T., Martínez-Pérez, E., Quaglia, F., Hatos, A., Chemes, Lucía B., Iserte, J.A., Méndez, N.A., Garrone, N.A., Saldaño, Tadeo E., Marchetti, J., et al. (2020). PED in 2021: a major update of the protein ensemble database for intrinsically disordered proteins. Nucleic Acids Research 49, D404–D411. 10.1093/nar/gkaa1021.

185. Kriwacki, R.W., Hengst, L., Tennant, L., Reed, S.I., and Wright, P.E. (1996). Structural studies of p21Waf1/Cip1/Sdi1 in the free and Cdk2-bound state: conformational disorder mediates binding diversity. Proceedings of the National Academy of Sciences 93, 11504–11509. 10.1073/pnas.93.21.11504.

186. Berlow, R.B., Dyson, H.J., and Wright, P.E. (2017). Hypersensitive termination of the hypoxic response by a disordered protein switch. Nature 543, 447-451. 10.1038/nature21705.

187. Shnitkind, S., Martinez-Yamout, M.A., Dyson, H.J., and Wright, P.E. (2018). Structural Basis for Graded Inhibition of CREB:DNA Interactions by Multisite Phosphorylation. Biochemistry 57, 6964–6972. 10.1021/acs.biochem.8b01092.

188. Sun, X., Dyson, H.J., and Wright, P.E. (2021). A phosphorylation-dependent switch in the disordered p53 transactivation domain regulates DNA binding. Proceedings of the National Academy of Sciences 118, e2021456118. 10.1073/pnas.2021456118.

189. Borg, M., Mittag, T., Pawson, T., Tyers, M., Forman-Kay, J.D., and Chan, H.S. (2007). Polyelectrostatic interactions of disordered ligands suggest a physical basis for ultrasensitivity. Proceedings of the National Academy of Sciences 104, 9650–9655. 10.1073/pnas.0702580104.

190. Bah, A., Vernon, R.M., Siddiqui, Z., Krzeminski, M., Muhandiram, R., Zhao, C., Sonenberg, N., Kay, L.E., and Forman-Kay, J.D. (2015). Folding of an intrinsically disordered protein by phosphorylation as a regulatory switch. Nature 519, 106–109. 10.1038/nature13999.

191. Das, R.K., Crick, S.L., and Pappu, R.V. (2012). N-Terminal Segments Modulate the α-Helical Propensities of the Intrinsically Disordered Basic Regions of bZIP Proteins. Journal of Molecular Biology 416, 287–299. 10.1016/j.jmb.2011.12.043.

192. Fuertes, G., Banterle, N., Ruff, K.M., Chowdhury, A., Mercadante, D., Koehler, C., Kachala, M., Estrada Girona, G., Milles, S., Mishra, A., et al. (2017). Decoupling of size and shape fluctuations in heteropolymeric sequences reconciles discrepancies in SAXS vs. FRET measurements. Proceedings of the National Academy of Sciences 114, E6342–E6351. doi:10.1073/pnas.1704692114.

193. Warner, J.B.I.V., Ruff, K.M., Tan, P.S., Lemke, E.A., Pappu, R.V., and Lashuel, H.A. (2017). Monomeric Huntingtin Exon 1 Has Similar Overall Structural Features for Wild-Type and Pathological Polyglutamine Lengths. Journal of the American Chemical Society 139, 14456–14469. 10.1021/jacs.7b06659.

194. Dogan, J., Gianni, S., and Jemth, P. (2014). The binding mechanisms of intrinsically disordered proteins. Physical Chemistry Chemical Physics 16, 6323–6331. 10.1039/C3CP54226B.

195. Jemth, P., Karlsson, E., Vögeli, B., Guzovsky, B., Andersson, E., Hultqvist, G., Dogan, J., Güntert, P., Riek, R., and Chi, C.N. (2018). Structure and dynamics conspire in the evolution of affinity between intrinsically disordered proteins. Science Advances 4, eaau4130. doi:10.1126/sciadv.aau4130.

196. Toto, A., Malagrinò, F., Visconti, L., Troilo, F., Pagano, L., Brunori, M., Jemth, P., and Gianni, S. (2020). Templated folding of intrinsically disordered proteins. Journal of Biological Chemistry 295, 6586–6593. 10.1074/jbc.REV120.012413.

197. Newcombe, E.A., Due, A.D., Sottini, A., Elkjær, S., Theisen, F.F., Fernandes, C.B., Staby, L., Delaforge, E., Bartling, C.R.O., Brakti, I., et al. (2024). Stereochemistry in the disorder–order continuum of protein interactions. Nature 636, 762–768. 10.1038/s41586-024-08271-6.

198. Theisen, F.F., Staby, L., Tidemand, F.G., O’Shea, C., Prestel, A., Willemoës, M., Kragelund, B.B., and Skriver, K. (2021). Quantification of Conformational Entropy Unravels Effect of Disordered Flanking Region in Coupled Folding and Binding. Journal of the American Chemical Society 143, 14540–14550. 10.1021/jacs.1c04214.

199. Elkjær, S., Due, A.D., Christensen, L.F., Theisen, F.F., Staby, L., Kragelund, B.B., and Skriver, K. (2023). Evolutionary fine-tuning of residual helix structure in disordered proteins manifests in complex structure and lifetime. Communications Biology 6, 63. 10.1038/s42003-023-04445-6.

200. Skriver, K., Theisen, F.F., and Kragelund, B.B. (2023). Conformational entropy in molecular recognition of intrinsically disordered proteins. Current Opinion in Structural Biology 83, 102697. 10.1016/j.sbi.2023.102697.

201. Mukhopadhyay, S., Krishnan, R., Lemke, E.A., Lindquist, S., and Deniz, A.A. (2007). A natively unfolded yeast prion monomer adopts an ensemble of collapsed and rapidly fluctuating structures. Proceedings of the National Academy of Sciences 104, 2649–2654. doi:10.1073/pnas.0611503104.

202. Yu, M., Heidari, M., Mikhaleva, S., Tan, P.S., Mingu, S., Ruan, H., Reinkemeier, C.D., Obarska-Kosinska, A., Siggel, M., Beck, M., et al. (2023). Visualizing the disordered nuclear transport machinery in situ. Nature 617, 162–169. 10.1038/s41586-023-05990-0.

203. Yu, M., Gruzinov, A.Y., Ruan, H., Scheidt, T., Chowdhury, A., Giofrè, S., Mohammed, A.S.A., Caria, J., Sauter, P.F., Svergun, D.I., and Lemke, E.A. (2024). A genetically encoded anomalous SAXS ruler to probe the dimensions of intrinsically disordered proteins. Proceedings of the National Academy of Sciences 121, e2415220121. doi:10.1073/pnas.2415220121.

204. Rogers, J.M., Wong, C.T., and Clarke, J. (2014). Coupled Folding and Binding of the Disordered Protein PUMA Does Not Require Particular Residual Structure. Journal of the American Chemical Society 136, 5197–5200. 10.1021/ja4125065.

205. Kim, J.-Y., and Chung, H.S. (2020). Disordered proteins follow diverse transition paths as they fold and bind to a partner. Science 368, 1253–1257. doi:10.1126/science.aba3854.

206. Appadurai, R., Koneru, J.K., Bonomi, M., Robustelli, P., and Srivastava, A. (2023). Clustering Heterogeneous Conformational Ensembles of Intrinsically Disordered Proteins with t-Distributed Stochastic Neighbor Embedding. Journal of Chemical Theory and Computation 19, 4711–4727. 10.1021/acs.jctc.3c00224.

207. Koren, G., Meir, S., Holschuh, L., Mertens, H.D.T., Ehm, T., Yahalom, N., Golombek, A., Schwartz, T., Svergun, D.I., Saleh, O.A., et al. (2023). Intramolecular structural heterogeneity altered by long-range contacts in an intrinsically disordered protein. Proceedings of the National Academy of Sciences 120, e2220180120. doi:10.1073/pnas.2220180120.

208. Baul, U., Chakraborty, D., Mugnai, M.L., Straub, J.E., and Thirumalai, D. (2019). Sequence Effects on Size, Shape, and Structural Heterogeneity in Intrinsically Disordered Proteins. The Journal of Physical Chemistry B 123, 3462–3474. 10.1021/acs.jpcb.9b02575.

209. Fuxreiter, M. (2018). Fuzziness in Protein Interactions—A Historical Perspective. Journal of Molecular Biology 430, 2278–2287. 10.1016/j.jmb.2018.02.015.

210. Song, J., Gomes, G.-N., Shi, T., Gradinaru, C.C., and Chan, H.S. (2017). Conformational Heterogeneity and FRET Data Interpretation for Dimensions of Unfolded Proteins. Biophysical Journal 113, 1012–1024. 10.1016/j.bpj.2017.07.023.

211. Hatos, A., Monzon, A.M., Tosatto, S.C.E., Piovesan, D., and Fuxreiter, M. (2021). FuzDB: a new phase in understanding fuzzy interactions. Nucleic Acids Research 50, D509–D517. 10.1093/nar/gkab1060.

212. Emenecker, R.J., Guadalupe, K., Shamoon, N.M., Sukenik, S., and Holehouse, A.S. (2023). Sequence-ensemble-function relationships for disordered proteins in live cells. bioRxiv, 2023.2010.2029.564547. 10.1101/2023.10.29.564547.

213. Krueger, R., Brenner, M.P., and Shrinivas, K. (2024). Generalized design of sequence-ensemble-function relationships for intrinsically disordered proteins. bioRxiv, 2024.2010.2010.617695. 10.1101/2024.10.10.617695.

214. Simard, R., and L’Ecuyer, P. (2011). Computing the Two-Sided Kolmogorov-Smirnov Distribution. Journal of Statistical Software 39, 1–18. 10.18637/jss.v039.i11.

215. Piovesan, D., Del Conte, A., Clementel, D., Monzon, A.M., Bevilacqua, M., Aspromonte, M.C., Iserte, J.A., Orti, F.E., Marino-Buslje, C., and Tosatto, S.C.E. (2023). MobiDB: 10 years of intrinsically disordered proteins. Nucleic Acids Research 51, D438–D444. 10.1093/nar/gkac1065.

216. Potenza, E., Di Domenico, T., Walsh, I., and Tosatto, S.C. (2015). MobiDB 2.0: an improved database of intrinsically disordered and mobile proteins. Nucleic Acids Res 43, D315–320. 10.1093/nar/gku982.

217. Nagaraj, N., Wisniewski, J.R., Geiger, T., Cox, J., Kircher, M., Kelso, J., Paabo, S., and Mann, M. (2011). Deep proteome and transcriptome mapping of a human cancer cell line. Molecular Systems Biology 7, 548. 10.1038/msb.2011.81.

218. Vitalis, A., and Pappu, R.V. (2009). ABSINTH: a new continuum solvation model for simulations of polypeptides in aqueous solutions. J Comput Chem 30, 673–699. 10.1002/jcc.21005.

219. Berman, H.M., Westbrook, J., Feng, Z., Gilliland, G., Bhat, T.N., Weissig, H., Shindyalov, I.N., and Bourne, P.E. (2000). The Protein Data Bank. Nucleic Acids Research 28, 235–242. 10.1093/nar/28.1.235.

220. Lalmansingh, J.M., Keeley, A.T., Ruff, K.M., Pappu, R.V., and Holehouse, A.S. (2023). SOURSOP: A Python Package for the Analysis of Simulations of Intrinsically Disordered Proteins. Journal of Chemical Theory and Computation 19, 5609–5620. 10.1021/acs.jctc.3c00190.

221. Edgar, R.C. (2024). Protein structure alignment by Reseek improves sensitivity to remote homologs. Bioinformatics 40. 10.1093/bioinformatics/btae687.

222. Marchler-Bauer, A., Derbyshire, M.K., Gonzales, N.R., Lu, S., Chitsaz, F., Geer, L.Y., Geer, R.C., He, J., Gwadz, M., Hurwitz, D.I., et al. (2015). CDD: NCBI’s conserved domain database. Nucleic Acids Research 43, D222–226. 10.1093/nar/gku1221.

